# Comparative mortality of dominant *Staphylococcus aureus* lineages in human bacteremia and animal infection models

**DOI:** 10.1101/2025.03.11.642576

**Authors:** Miquel Sánchez-Osuna, Marc Bravo, María-Alexandra Cañas, Andrómeda-Celeste Gómez, Inmaculada Gómez-Sánchez, José M. Miró, Isidre Gibert, Oriol Gasch, Cristina García-de-la-Mària, Daniel Yero, Oscar Q. Pich

## Abstract

*Staphylococcus aureus* is a major cause of severe infections including infective endocarditis, but lineage-specific virulence determinants remain unclear. We analyzed 77 *S. aureus* bacteremic isolates from major lineages using phenotypic assays, infection models, and transcriptomics. Our results revealed significant heterogeneity in *S. aureus* pathogenicity. ST398 isolates exhibited heightened virulence, characterized by increased hemolysin production, whereas CC30 strains showed reduced growth, biofilm formation, and infectivity. Notably, the ST398 *agrC* mutant Sau7 exhibited unique phenotypic behavior, with high biofilm production and decreased virulence in *Galleria mellonella* larvae model. Infection studies in the rabbit experimental endocarditis model showed increased vegetation size and bacterial load in Sau7-infected animals, highlighting the role of *agr* system in *S. aureus* colonization and biofilm formation. Transcriptomic analysis identified key pathways, including quorum sensing systems and hemolysins, driving virulence in ST398 strains. These findings provide insights into the lineage-specific virulence mechanisms and the multifaceted nature of *S. aureus* pathogenicity.

## INTRODUCTION

*Staphylococcus aureus* is a cocci-shaped bacterium that acts both as a commensal microbe and a human major pathogen^1^. It is generally found in the skin and nasal mucosa in 20-40% of the population^2^. However, when these barriers are damaged (e.g. wounds, intravenous drug use or surgery), the bacterium can gain access to the bloodstream and underlying tissues causing infection^3^. Due to its clinical relevance, bacteremia is one of the well-studied manifestations of *S. aureus* infections, with an estimated incidence of around 40 episodes per 100,000 people per year^4^. Bacteremia is a life-threatening clinical condition associated with high morbidity and mortality that often results in metastasis, causing complications such as infective endocarditis (IE), discitis, osteomyelitis or muscular abscesses^1^.

The development of *S. aureus* infections depends on both host characteristics and bacterial factors^5^. Severity of bacterial infections is influenced by host factors such as age, comorbidities, genetics, previous infections, nutrition, and the overall environment^6^. Poor clinical outcomes of *S. aureus* infections have been associated with the patient’s age, which is the most consistent predictor of mortality, the presence of certain comorbidities and the antimicrobial therapy used^7^. However, the impact of bacterial factors in *S. aureus* infection’s outcome is scarcely known. Although methicillin and vancomycin resistance may correlate with mortality, this has been associated with nosocomial strains affecting patients with significant comorbidities^7,8^. Conversely, several lineages have been associated with increased risk of complications^9,10^, while more recent studies did not find any apparent association between clonality and adverse outcomes^10,11^.

This lack of consistency underscores the necessity of studies evaluating *S. aureus* factors that genuinely influence infection outcomes. The use of animal models enables testing multiple specimens ensuring a consistent genetic background and comorbidities, under controlled conditions. This approach is highly effective for analyzing bacterial infections, as it reduces the potential influence of host-related variables. Different vertebrate and invertebrate animal models have been used for *S. aureus* so far; including mice (*Mus musculus*)^12^, rats (*Rattus norvegicus*)^13^, rabbits (*Leporidae spp.*)^11^, wax worm (*Galleria mellonella)*^14^ and nematodes (*Caenorhabditis elegans*)^15^. However, these studies have primarily focused on the treatment or infectious traits of specific strains, rather than conducting high-throughput analyses with a wide collection of isolates.

Herein, we used two experimental infection models to evaluate the pathogenicity of 77 *S. aureus* isolates from bacteremia, representing the dominant lineages in a hospital from the Barcelona metropolitan area^16^. Lethality of all the clinical isolates was assessed in *G. mellonella* larvae. These analyses allowed the identification of hypervirulent and attenuated lineages, as well as outliner strains within specific lineages. The contribution of specific virulence determinants to the mortality observed in *G. mellonella* was further explored by combining comparative genomics, transcriptomics, determination of the hemolysis type and biofilm formation capacity. Then, strains representative of the lineages with the more extreme virulent phenotypes in *G. mellonella* were selected for infection assays in an experimental endocarditis (EE) model in rabbits, including a detailed bacterial quantification across different organs. Overall, our results suggest that virulence in *G. mellonella* is influenced by the *agr* type and hemolytic activity. While these factors are still important in the EE model in rabbits, strains with a deficient *agr* function are still capable of causing death, which correlates with the formation of large vegetations in key organs. Therefore, our study confirms that animal models are essential to reveal *bona fide* virulence factors, which could otherwise be difficult to recognize in human infections due to the inherent heterogeneity of the hosts. Moreover, our study demonstrates the usefulness of the *G. mellonella* to conduct large-scale infection studies and to identify factors of mortality by *S. aureus*.

## MATERIAL AND METHODS

### Bacterial isolates

This study examined a cohort of 77 *S. aureus* isolates, selected as a representative panel from a larger collection of 339 *S. aureus* strains isolated from patients with bacteremia at the Parc Taulí University Hospital (Sabadell, Spain) between July 2014 and December 2022^16^. Specifically, *S. aureus* isolates were randomly selected to include about half of the strains from the most prevalent lineages in the large collection. All these isolates are sequenced and lineages, antibiotic resistance genes and virulence factors are already characterized. The computational workflow for these analyses has been previously described in detail^17^. Whole-genome sequencing data are available at National Center for Biotechnology Information (NCBI) under the accession number PRJNA1055690. Patient’s clinical information is also recorded.

### *S. aureus* phenotypic characterization

#### Growth curves

*S. aureus* growth curves were performed on 96-well microtiter plates with 0.5X Tryptic Soy Broth (TSB) supplemented with 1% glucose and were determined using a Multiskan FC microplate photometer (Thermo Fisher Scientific) under constant shaking at 37°C for 24 hours. OD_550_ was recorded every 15 min. Growth rate (h^−1^) was computed using the best-fit values of the exponential growth model, where the growth rate corresponds to the parameter *k* in the equation 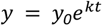. The doubling time was then calculated using the formula *ln*(2)/*k*.

#### Biofilm quantification

Quantitative analysis of *S. aureus* biofilm formation was performed with crystal violet assay, similar to previously described^14^. *S. aureus* overnight cultures grown on Columbia Agar with 5% Sheep Blood plates (Becton Dickinson) were collected and adjusted to an OD_550_ of 0.05 in 0.5X TSB supplemented with 1% glucose. Then, 96-well flat-bottom microtiter plates (non-treated and sterile) were inoculated with 200 μL of bacterial suspensions and incubated at 37°C under 5% CO_2_ for 24 hours. The amount of biofilm was quantified by measuring the OD_550_ of dissolved crystal violet using a Multiskan FC microplate photometer (Thermo Fisher Scientific).

#### Hemolysis activity assays

Hemolytic activity of *S. aureus* was inferred using the Christie-Atkins-Munch-Peterson (CAMP) synergistic hemolysis test against *S. aureus* strain RN4220 on Columbia agar plates^18^. *S. aureus* overnight cultures grown on Columbia Agar plates were collected and adjusted to a 0.5 McFarland in Phoenix™ ID Broth (Becton Dickinson) using a PhoenixSpec™ nephelometer (Becton Dickinson). After incubation at 37°C for 24 hours, the plates were refrigerated to increase hemolytic activity. All assays were performed three times in independent experiments.

### *In vivo* studies in the *G. mellonella* model

Infection studies of *G. mellonella* (wax worm) larvae were performed as previously described, with slight modifications^14^. Larvae were obtained from an in-house hatchery, and only healthy-looking larvae weighing 200-300 mg were selected for further experiments. Bacterial inoculums were prepared in Phosphate Buffered Saline (PBS) using *S. aureus* cells grown overnight on Columbia agar plates, and adjusted to OD_550_ of 0.125. Fifteen larvae were individually infected per isolate by injecting 10 μL of bacterial suspension (~5×10^6^ cfu/larva) into the left proleg, and incubated in darkness at 37°C. Mortality was determined every 24 h, and larvae were considered dead when totally melanized and/or no longer responded to touch. The *S. aureus* strains Newman and SAR5091 were included as the reproducibility controls in all experiments. Inoculum bacterial counts were confirmed by plating on Columbia agar plates. Kaplan-Meier survival curves were plotted using GraphPad Prism. A logistic regression model was fitted using the proportion of dead individuals over time, and Lethal time 50 (LT_50_) was estimated using the “dose.p” function from the MASS package in R. LT_50_ was defined as the time required for the inoculum to kill 50% of the infected larvae. Mann-Whitney U (MWU) test was used for comparing distributions of LT_50_ values.

### *In vivo* studies in the EE rabbit model

#### Study animals

New Zealand white rabbits (body weight, 2.5 kg) provided by San Bernardo farm (Pamplona, Spain) were used. Housing took place in the animal facilities of the University of Barcelona, School of Medicine, which is equipped with a high-efficiency particulate air filter in an automatic air exchange system, as well as a circadian light cycle. Rabbits were nourished *ad libitum*. All animal experimentation in this study was approved by the Animal Ethics and Research Committee of the University of Barcelona and the Generalitat de Catalunya (Barcelona, Spain).

#### Strains selected

Based on the results obtained from the *G. mellonella* model, two strains (Sau22, Sau65) from CC30 (presenting the highest survival rate) and two strains (Sau56, Sau164) from ST398 (with the lowest survival rate) were selected for study in the EE model. The *S. aureus* Sau7 (ST398) isolate with an *agrC* disruption was also included.

#### Experimental endocarditis model

The experimental aortic valve IE model was induced according to the method described by Garrison and Freedman^19^. Briefly, the animals were anaesthetized and a polyethylene catheter was inserted through the right carotid artery into the left ventricle to produce an aortic valve injury. Twenty-four hours later, animals were randomized into the different infecting strain groups. They were inoculated via the marginal ear vein with 1 mL of 5 × 10^6^ colony forming units [cfu/mL] of one of the selected strains (Sau7, Sau22, Sau65, Sau56 or Sau164). Twenty-four hours after inoculation, 2 mL of blood were obtained to confirm bacteremia. Animals were supervised every 8 hours. Post-mortem necropsy was performed. The vegetations of the aortic valve, spleen, the left kidney and the brain were removed under sterile conditions. The middle third of the spleen and kidney and the left hemisphere of the brain were selected for culture. Samples were weighed, homogenised in 2 mL of saline solution and cultured quantitatively. The number of colony forming units (cfu) recovered from tissues or vegetation was expressed as the number of log_10_ cfu per gram of tissue^20^. Each group contained ten evaluable animals: those with positive blood cultures 24 hours after the inoculum and positive bacterial growth in vegetation at the time of necropsy.

#### Statistical analysis

Survival time from inoculation was measured in hours. Bacterial density in the different tissues was expressed as the median and the interquartile range (IQR) of the number of log_10_ cfu/g tissue or vegetation, as well as the vegetation weight. The non-parametric Mann Whitney test was used to compare the log_10_ cfu tissue values and the survival time. The Log-rank (Mantel-Cox) test was used to compare the survival rates among the different groups in the Kaplan-Meier survival curve.

### Transcriptomic analysis

#### RNA extraction

Exponential cultures of *S. aureus* were grown on TSB at 37°C. After 3 hours, total RNA was extracted using the RNeasy mini-kit (Qiagen) followed by a DNAse treatment with the TURBO-DNA free kit (Thermo Fisher Scientific). Cell lysis was achieved by pre-incubating culture pellets in PBS containing 100 μg/mL lysostaphin (Merck) at 37°C for 15 minutes. RNA quality was assessed using a RNA Nano Chip (Agilent Technologies).

#### RNA sequencing

RNA library preparation was performed using the Illumina Stranded Total RNA Prep Ligation with Ribo-Zero Plus kit and 10bp unique dual indices. Sequencing was done on a NovaSeq X Plus (SeqCenter, USA), producing paired-end 150bp reads. RNA sequencing data is available at National Center for Biotechnology Information (NCBI) under the accession number PRJNA1212992.

#### Differential expression analysis

Raw sequencing data quality was evaluated with FastQC (https://github.com/s-andrews/FastQC, accessed in May 2024), filtered with TrimGalore (https://github.com/FelixKrueger/TrimGalore, accessed in May 2024) and residual rRNA reads were removed with the SortMeRNA software^21^. Filtered reads were then mapped using Bowtie2^22^ against the *S. aureus* NCTC 8325 genome [GCF_000013425] and alignments were quantified with FADU^23^. Differential expression gene (DEG) analysis was finally performed with DESeq2^24^. Genes showing −1.0 <= log2 fold-change >= 1.0 and adjusted *p*-value <0.05 were considered as DEGs. DEG products were then analyzed using ShinyGO for gene ontology (GO) enrichment^25^ and mapped to eggNOG^26^ and to Virulence Factor Database^27^. Genes with 0 FADU counts in one of the genomes were removed for further analysis.

### *S. aureus* ST398 comparative genomics

To elucidate the genetic causes governing the differential virulence observed in *S. aureus* ST398 Sau7 strain, pairwise orthologs detection of all ST398 isolates was performed. Orthologs were compiled from *S. aureus* Sau7 strain via reciprocal BLASTP^28^ using all protein-coding gene sequences encoded in ST398 genomes, using a stringent threshold of a conservative e-value of <1e–20 and query coverage of >75%. Protein-coding genes present in all ST398 genomes but absent in the *S. aureus* Sau7 genome were selected for further investigation. This analysis was implemented using a custom Python script^29^. Transposon insertion within *agrC* gene was verified via PCR using appropriate primers (Table S1).

### Phylogenetic inference

Phylogenetic analysis was performed using core genome alignments based on single nucleotide polymorphisms (SNPs) called by snippy (https://github.com/tseemann/snippy, accessed in May 2024) against the *S. aureus* NCTC 8325 genome [GCF_000013425]. A Maximum Likelihood (ML) tree was constructed with IQ-TREE 2^30^ using 1000 bootstrap replicates and TVM+F+G4 as the substitution model, as deduced from ModelFinder^31^. Tree visualization and annotation were performed with iTOL^32^.

### Ethics statement

The Ethics Committee for Investigation with medicinal products (CEIm) of the Parc Taulí University Hospital approved the implementation of this study (2025/4001, approved on 28 January 2025). The requirement for informed written consent was waived given the retrospective nature of the study. Patient identification was encoded, complying with the requirements of the Spanish Organic Law on Data Protection 15/1999.

## RESULTS

### Genetic and clinical data associated with the *S. aureus* isolates under study

This study was conducted using a panel of seventy-seven *S. aureus* clinical isolates, selected from a larger collection^16^, from patients with bacteremia treated at the Parc Taulí University Hospital. The isolates comprised strains from the most prevalent lineages and it was representative of *S. aureus* bacteremia cases at the hospital (Figure 1). Specifically, the selected isolates belonged to the following lineages: CC30 (n=14), CC1 (n=11), CC5 (n=10), ST398 (n=10), CC8 (n=8), CC45 (n=8), CC22 (n=5), CC15 (n=4), ST7 (n=3), CC121 (n=2), and ST291 (n=2). Regarding clinical outcomes, 28 cases were fatal (36.3%), 23 resulted in sepsis (29.9%), 19 led to persistent bacteremia (24.7%), 18 resulted in septic embolism (23.4%) and two strains caused IE (2.6%) in our patients. Eight of the isolates (10.3%) were methicillin-resistant *S. aureus* (MRSA) according to antibiotic susceptibility tests. Whole-genome sequencing revealed that the most prevalent *S. aureus* virulence factors in our dataset included adhesion (*clfAB, sdrC, fnbA*) and immune evasion factors (*sbi, scn*); hemolysins (*hla, hlb, hld, hlgABC*), and biofilm-associated genes (*icaABCD*). Iron acquisition (*isd* genes) and secretion system components (*esa, ess, esxA*) were also highly conserved. Toxins like the Panton-Valentine leucocidin, TSST-1, and enterotoxins (*sea, seb*) were rare. Clinical and molecular data of all *S. aureus* strains are summarized in Data S1. Bioinformatic analyses conducted by our group did not reveal significant associations between the genetic features of the isolates and the clinical outcomes^16^.

**Figure 1.**
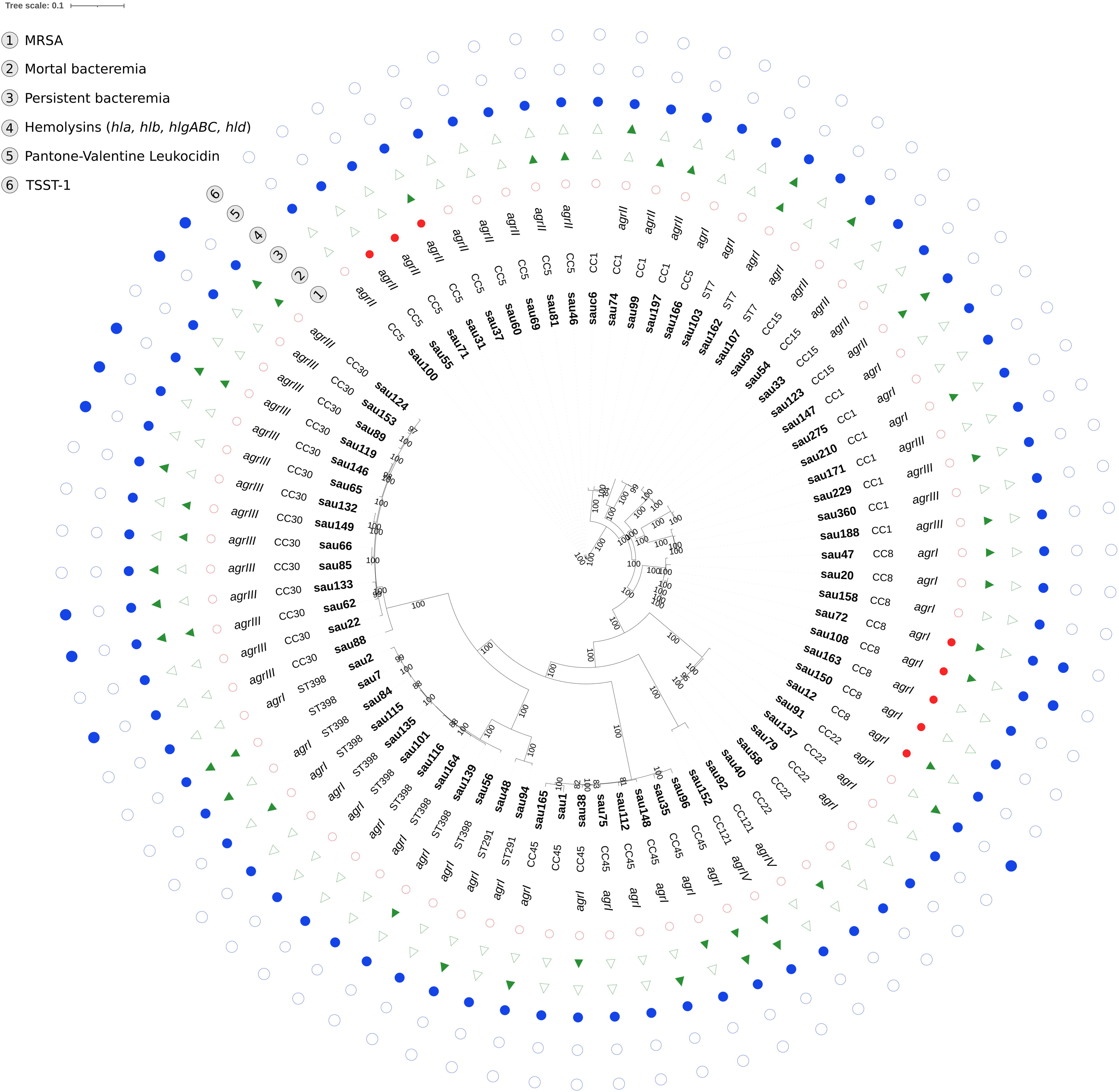
Core genome phylogenetic tree of *S. aureus* isolates analyzed in this study. Branch support values are provided as the percent of bootstrap replicates in which the branching was observed. Support values are only shown for branches with at least 80% support. The tree includes annotations for lineage, *agr* type, MRSA strains, bacteremia outcome, and the presence of hemolysins, Panton-Valentine Leukocidin (PVL), and TSST-1 toxin.

### Growth rate, biofilm formation capacity and hemolytic activity

To expand the phenotypic data, we determined the growth rate, biofilm formation capacity and hemolytic activity of all the isolates under study. Wilcoxon rank-sum test was performed to determine whether any lineage exhibited significant differences compared to the overall dataset (Figure 2AB, Table S2). The growth rate analysis showed an overall mean of 1.5 h^−1^ ± 0.2 SD across all isolates, with slight variations among lineages. Briefly, ST398 isolates exhibited the highest mean growth rate (1.7 h^−1^), followed by ST7 (1.6 h^−1^) and CC22 (1.6 h^−1^). In contrast, growth rate of CC30 (1.4 h^−1^) and ST291 (1.4 h^−1^) isolates was moderately below the average. For biofilm formation, the global mean was 0.9 ± 1.3 SD. Specifically, CC15 isolates displayed the highest mean biofilm formation (2.9), followed by CC1 (1.3) and ST7 (1.6). Conversely, CC22 (0.5) and CC30 (0.4) showed the lowest biofilm formation capacity. Notably, CC30 was the only lineage showing a statistically significant difference (MWU, *p* < 0.05) for both phenotypes, displaying significantly lower biofilm formation and a reduced growth rate compared to other lineages (Figure 2AB).

**Figure 2.**
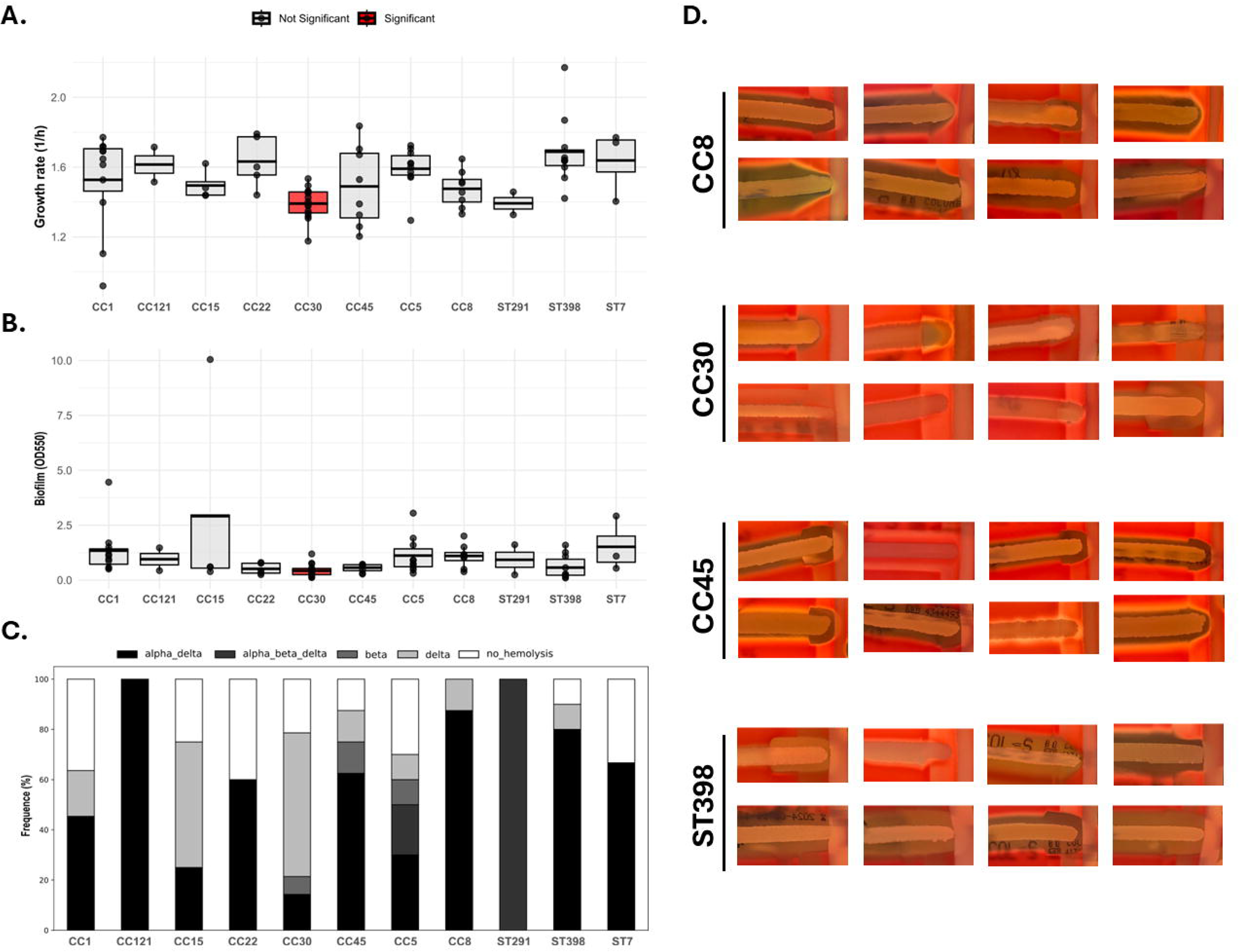
Summary of the phenotypic characterization of *S. aureus* isolates analyzed in this study. The figure includes box plots depicting the growth rate (A) and biofilm formation (B) across different *S. aureus* lineages. Lineages exhibiting significant differences are highlighted in red. (C) Stacked bar plot illustrating the distribution of hemolysis types across all lineages. (D) Hemolysis activity assay results for representative isolates from diverse *S. aureus* lineages.

To assess the hemolytic activity of *S. aureus* isolates, we conducted the CAMP synergistic hemolysis test on Columbia agar plates using the reference strain RN4220. The results revealed variations in hemolysis types among isolates, with certain lineages displaying distinct hemolytic patterns (Figure 2C, Figure S1). The majority of the CC30 isolates exhibited δ-hemolysis (57.2%), although 14.3% showed α-hemolysis, 7.1% β-hemolysis and 21.4% were non-hemolytic. ST398 showed predominantly α- and δ-hemolysis (80%), followed by δ-hemolysis (10%) or no hemolysis (10%). CC45 isolates were predominantly α- and δ-hemolytic 62.5%, while 12.5% displayed β- or δ-hemolysis, and 12.5% were non-hemolytic. CC8 was characterized by a strong α- and δ-hemolytic phenotype (87.5%), with the remaining 12.5% showing δ-hemolysis. In contrast, CC5 displayed a more balanced distribution, with 30% α- and δ-hemolysis; 20% α-, β- and δ-hemolysis; 10% β- and δ-hemolysis each; and 30% non-hemolytic isolates. Altogether, the phenotypic characterization conducted confirmed the heterogeneity of the*S. aureus* isolates under study.

### Infection assays in *G. mellonella* larvae

Infection studies in *G. mellonella* revealed a wide range of LT_50_ values across the analyzed *S. aureus* isolates, including strains causing larval mortality in less than 24 hours and strains for which larvae survived for the full 5-day duration of the experiment (Figure 3, Figure S2, Table S2). The consistent LT_50_ values observed for the control strains *S. aureus* Newman (53.1 h ± 2.7 SD) and SAR5091 (24.7 h ± 1.1 SD) in all infection experiments underscored the high reproducibility and robustness of the method.

**Figure 3.**
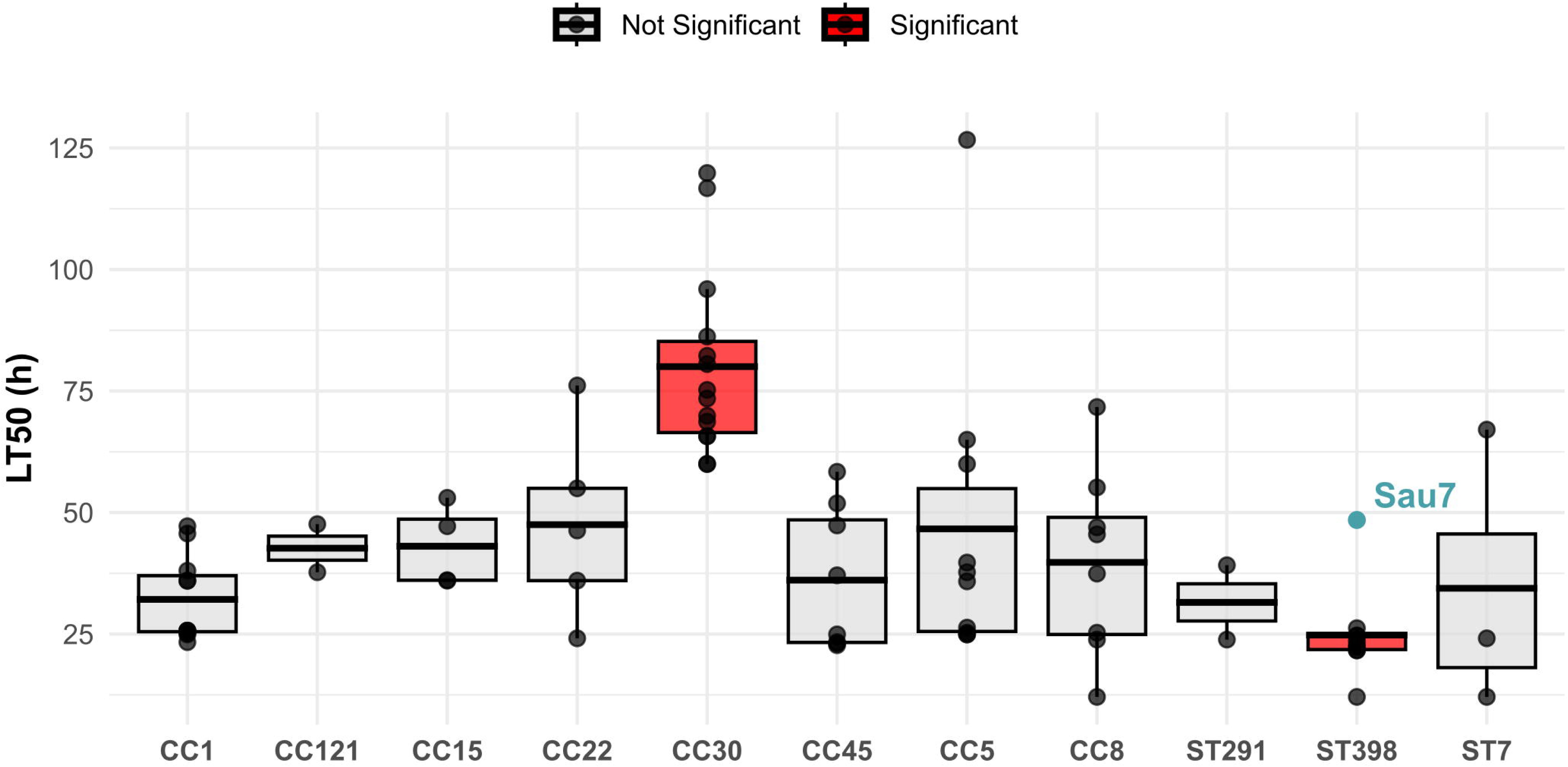
Box plot showing the LT_50_ (h) distribution across all analyzed *S. aureus* lineages tested in the *G. mellonella* model. Each dot represents an individual isolate. Lineages with significant differences are highlighted in red. The Sau7 isolate, showing a transposon insertion within *agrC* gene, is highlighted in blue.

Analysis of mean LT_50_ values across lineages revealed distinct patterns in larvae mortality (One-way ANOVA, *p* < 0.05). As shown in Figure 3, isolates of the CC30 lineage exhibited low mortality values, with an average LT_50_ of 80.0 h ± 18.4 SD. Specifically, we observed a significant difference in LT_50_ distribution of CC30 isolates when compared to the rest of lineages (MWU *p* < 0.05). Conversely, ST398 strains were the most lethal strains in *G. mellonella* larvae (LT_50_ = 24.8 h ± 8.7 SD), showing significantly different LT_50_ distributions compared to CC1, CC5, CC15, CC22 and CC30 lineages (MWU *p* < 0.05), underscoring a pronounced virulence of ST398 in this model. Of note, the *S. aureus* isolate Sau7 (ST398) displayed a low mortality (LT_50_ = 48.5 h) when compared to the other ST398 strains. Comparative genomics analysis revealed an IS30-like transposase insertion at the position 534 of the *agrC* coding sequence in Sau7, which was further confirmed by PCR (Figure S3). The Sau7 isolate was also the highest biofilm producer within the ST398 lineage (Table S2). Core genome phylogenetic analysis revealed that strains from the CC30 and ST398 lineages form closely related clusters, suggesting that they share a recent common ancestor and indicating a potential shared evolutionary trajectory (Figure 1). Accordingly, the reservoir of virulence factors of these two lineages is remarkably similar (Data S1), with 39 overlapping genes, including *hly/hla*, *hlgABC*, *hlb* and *hld* among others. However, CC30 exhibits higher frequencies of *clfB*, *sak, sea*, *tsst-1, vWbp and cap8* genes while ST398 harbors *coa, fnbAB* and *cap5* genes, which are absent in CC30.

The remaining lineages displayed intermediate lethality, with mean LT_50_ values ranging from 31.5 to 47.5 hours, and considerable heterogeneity in LT_50_ values within each lineage. No significant differences in LT_50_ were observed among these lineages. However, we found that two additional isolates from our study with truncated *agrC* genes (Data S1), Sau58 (LT_50_ = 54.99) and Sau40 (LT_50_ = 76.14) were the least lethal CC22 isolates (CC22 mean LT_50_ = 47.5) (Table S2). In contrast, double *agrBC* disruption, detected in Sau1 (CC45) and Sauc6 (CC1) isolates (Data S1), was not associated with reduced lethality in our study (Table S2).

In order to identify possible phenotypic features associated with the lethality observed in the *G. mellonella* model, we computed the Pearson Correlation Coefficient (PCC) between growth rate and biofilm formation with the LT_50_ observed in *G. mellonella* larvae. Despite the previously observed phenotypic differences in CC30 strains, including lower growth rates and reduced biofilm formation, LT_50_ values did not show a global correlation with either growth rate (PCC = −0.26) or biofilm formation (PCC = 0.0) in this study (Figure 4AB). In contrast, the analysis of LT_50_ and hemolysis types revealed significant differences (One-way ANOVA, *p* < 0.05). Specifically, the results showed a significant increase in LT_50_ in *G. mellonella* for non-hemolytic strains compared to those exhibiting α- and δ-hemolysis (Figure 4C). These results were similar when comparing LT_50_ and *agr* type (One-way ANOVA, *p* < 0.05), with *agrIII* being associated with higher LT_50_ values compared to both *agrI* and *agrII* types (Figure 4D).

**Figure 4.**
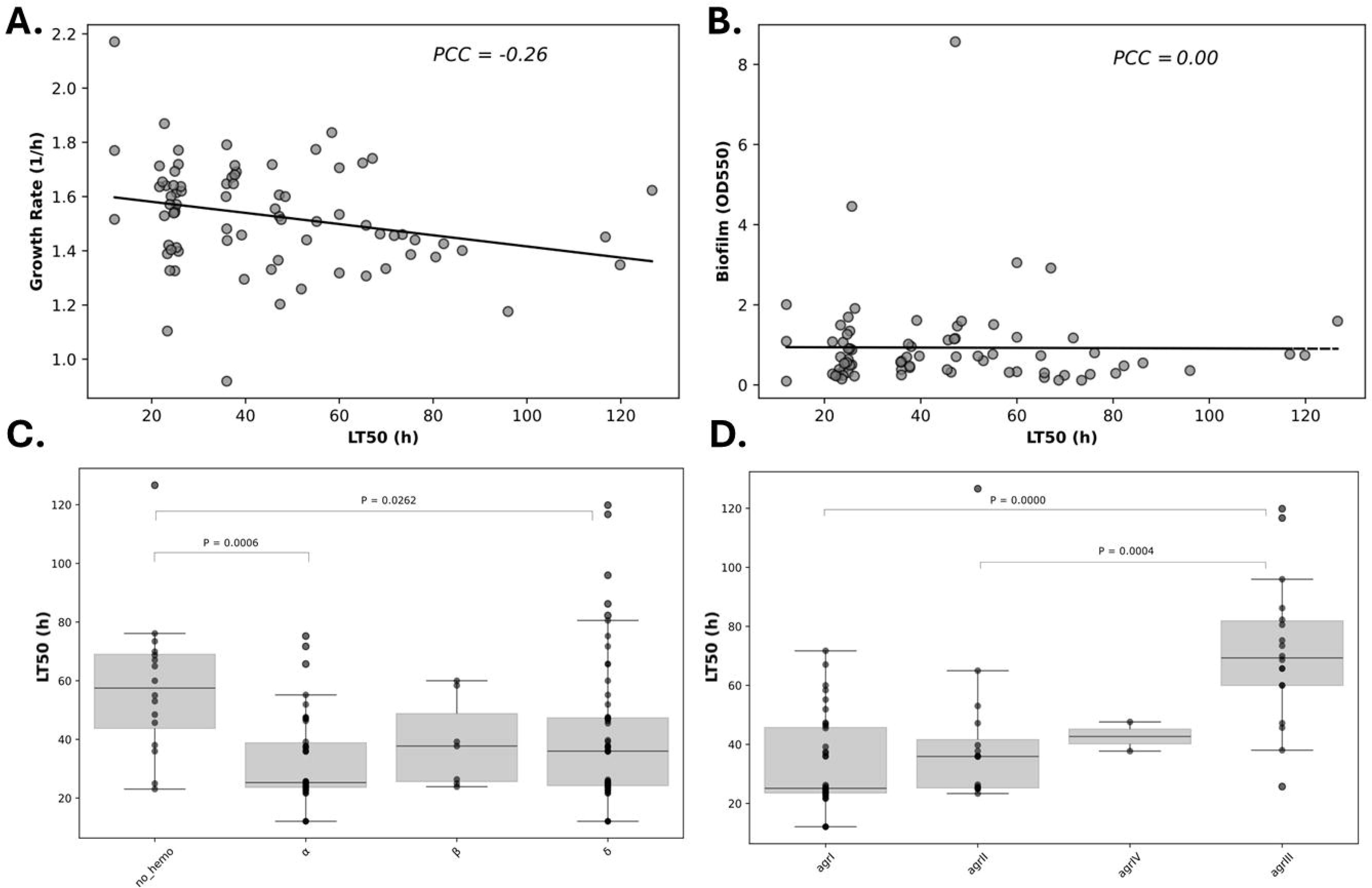
Plots showing the correlation between the growth rate (A), biofilm formation (B), hemolysis (C) and *agr* type (D) with LT_50_ in the *G. mellonella* model.

### *In vivo* results in the EE model in rabbits

The infectivity of selected ST398 and CC30 strains was evaluated in the rabbit EE model by using the following parameters: survival time, bacterial density in aortic vegetation and in peripheral organs (spleen, kidney and brain). The size of the vegetation was also taken into account.

#### Survival rates

The results of the survival rates for the different study strains are shown in Figure 5. The highest survival rates were obtained when animals were infected with Sau22 (CC30), which showed significantly reduced lethality compared to all the other groups. These differences were statistically significant in all cases, as shown in Figure 5A. Consequently, a significant difference (*p* = 0.002) was detected in the overall comparison (Figure 5A) and in pairwise comparison of the strains using Kaplan-Meier curves (Figure 5B). Although animals infected with ST398 strains had a lower survival rate (median 72 h for Sau56 and Sau164), no significant differences were found when it was compared to Sau65 (CC30) or the mutant ST398 strain (Sau7) groups with median 90 h and 85 h of survival respectively.

**Figure 5.**
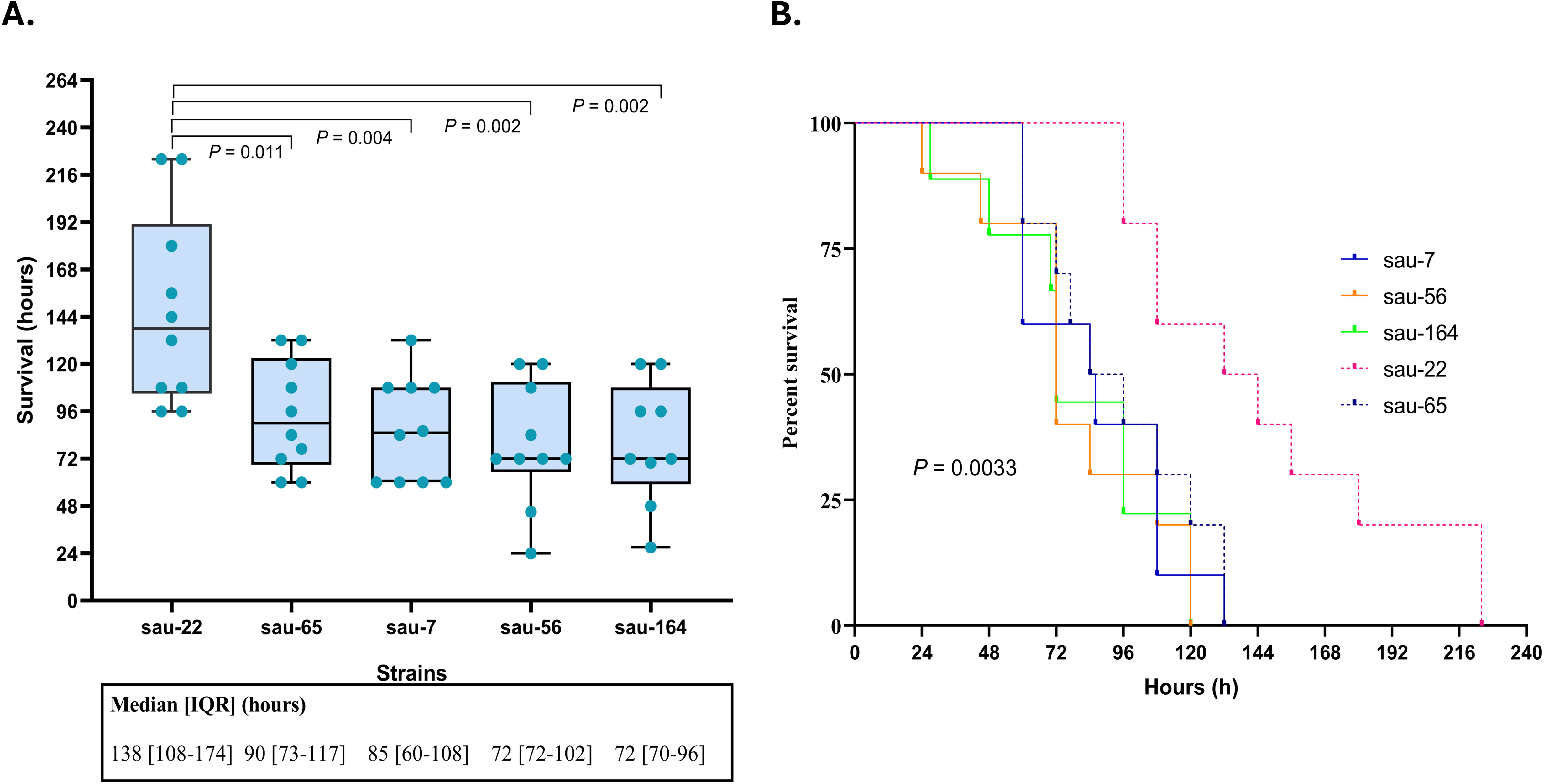
Summary of the IE rabbit model results. (A) Box plot depicting survival rates of different *S. aureus* strains over time in an experimental endocarditis model. Animals infected with Sau22 (CC30) showed the highest survival rates compared to all other groups. These differences were statistically significant in all cases. (B) The survival curves of rabbits following infection with *S. aureus* strains Sau22, Sau65, Sau7, Sau56, and Sau164 is represented by a Kaplan-Meier survival plot and expressed as percentage of survival vs. time. (p < 0.0033).

In the Sau65-infected group (CC30), it is of note that all samples obtained for two animals were sterile at the time of necropsy after euthanasia (two weeks after inoculation), despite presenting positive blood cultures 24 h after inoculation. These two animals were not included in the mean survival of the group as they did not develop endocarditis, probably due to the lower infectivity of the strain.

#### Bacterial CFU/g tissue density comparisons

The bacterial densities in aortic vegetations, spleen, kidney, and brain are presented in Table 1. Bacterial growth in vegetation was comparable across all groups, with a median density of approximately log10 = 10 CFU/g of vegetation. However, a significant increase in bacterial load was observed in animals infected with *S. aureus* strain Sau65 (CC30) compared to the strains Sau56 and Sau164 (ST398) (*p* < 0.022 and *p* < 0.018, respectively). Notably, the Sau7-infected group showed a similar density to Sau22 and Sau65 and exhibited significantly higher bacterial densities than the Sau56-infected group (*p* = 0.033) as it is displayed in Figure S4. Regarding the bacterial densities observed in peripheral organs such as the spleen and kidney, it is noteworthy that strains belonging to the clonal complex CC30, as well as strain Sau7 (ST398 mutant), exhibited higher densities than the two ST398 strains. However, the difference was statistically significant only between Sau65 and Sau56 and Sau164, as shown in Table 1 and Figure S4. As expected, the bacterial density in the brain was lower than in other organs, except in animals infected with Sau7. In these cases, a significantly higher number was observed, with a median of 8 log_10_ CFU/g, like the densities found in the spleen and kidney. This represents an increase of 1.5 to 2.5 log_10_ compared to other strains. These results are depicted in Table 1 and Figure S4.

**Table 1.**
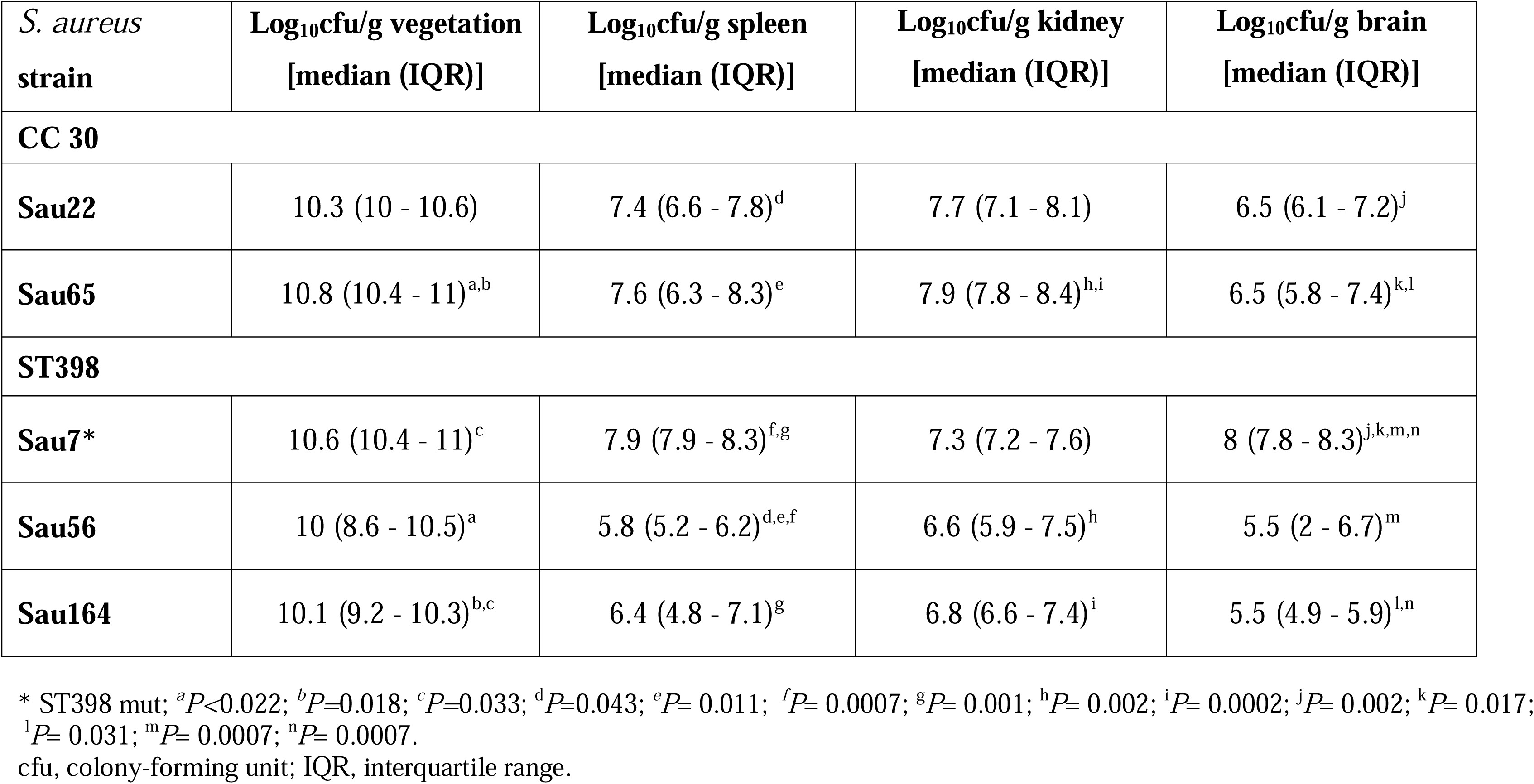
Results of bacterial growth of the selected *S. aureus* strains in vegetations, spleen, kidney and brain in a rabbit model of experimental endocarditis.

| *S. aureus*<br>strain | Log<sub>10</sub>cfu/g vegetation<br>[median (IQR)] | Log<sub>10</sub>cfu/g spleen<br>[median (IQR)] | Log<sub>10</sub>cfu/g kidney<br>[median (IQR)] | Log<sub>10</sub>cfu/g brain<br>[median (IQR)] |
| --- | --- | --- | --- | --- |
| **CC 30** |  |  |  |  |
| **Sau22** | 10.3 (10 - 10.6) | 7.4 (6.6 - 7.8)<sup>d</sup> | 7.7 (7.1 - 8.1) | 6.5 (6.1 - 7.2)<sup>j</sup> |
| **Sau65** | 10.8 (10.4 - 11)<sup>a,b</sup> | 7.6 (6.3 - 8.3)<sup>e</sup> | 7.9 (7.8 - 8.4)<sup>h,i</sup> | 6.5 (5.8 - 7.4)<sup>k,l</sup> |
| **ST398** |  |  |  |  |
| **Sau7\*** | 10.6 (10.4 - 11)<sup>c</sup> | 7.9 (7.9 - 8.3)<sup>f,g</sup> | 7.3 (7.2 - 7.6) | 8 (7.8 - 8.3)<sup>j,k,m,n</sup> |
| **Sau56** | 10 (8.6 - 10.5)<sup>a</sup> | 5.8 (5.2 - 6.2)<sup>d,e,f</sup> | 6.6 (5.9 - 7.5)<sup>h</sup> | 5.5 (2 - 6.7)<sup>m</sup> |
| **Sau164** | 10.1 (9.2 - 10.3)<sup>b,c</sup> | 6.4 (4.8 - 7.1)<sup>g</sup> | 6.8 (6.6 - 7.4)<sup>i</sup> | 5.5 (4.9 - 5.9)<sup>l,n</sup> |

\* ST398 mut; <sup>a</sup>*P*<0.022; <sup>b</sup>*P*=0.018; <sup>c</sup>*P*=0.033; <sup>d</sup>*P*=0.043; <sup>e</sup>*P*= 0.011; <sup>f</sup>*P*= 0.0007; <sup>g</sup>*P*= 0.001; <sup>h</sup>*P*= 0.002; <sup>i</sup>*P*= 0.0002; <sup>j</sup>*P*= 0.002; <sup>k</sup>*P*= 0.017;
<sup>l</sup>*P*= 0.031; <sup>m</sup>*P*= 0.0007; <sup>n</sup>*P*= 0.0007.
cfu, colony-forming unit; IQR, interquartile range.

#### Vegetation size comparison

When studying the weight of vegetations in the different groups, we observed that similar weights were obtained for the strains, regardless of whether the infecting strain belonged to CC30 or ST398 (Figure 6). But when the infecting strain was Sau7, significantly larger vegetations were obtained (*p* < 0.001 in all comparisons). This fact, as already mentioned, is consistent with the high biofilm production ability of this strain (Table S2). It is noteworthy that the ST398 strain Sau7, in contrast to the results obtained in the *G. mellonella* model, showed a similar mortality rate to the other ST398 strains, with mean LT_50_ values ranging from 78.9 h to 86.6 h (Figure 5).

**Figure 6.**
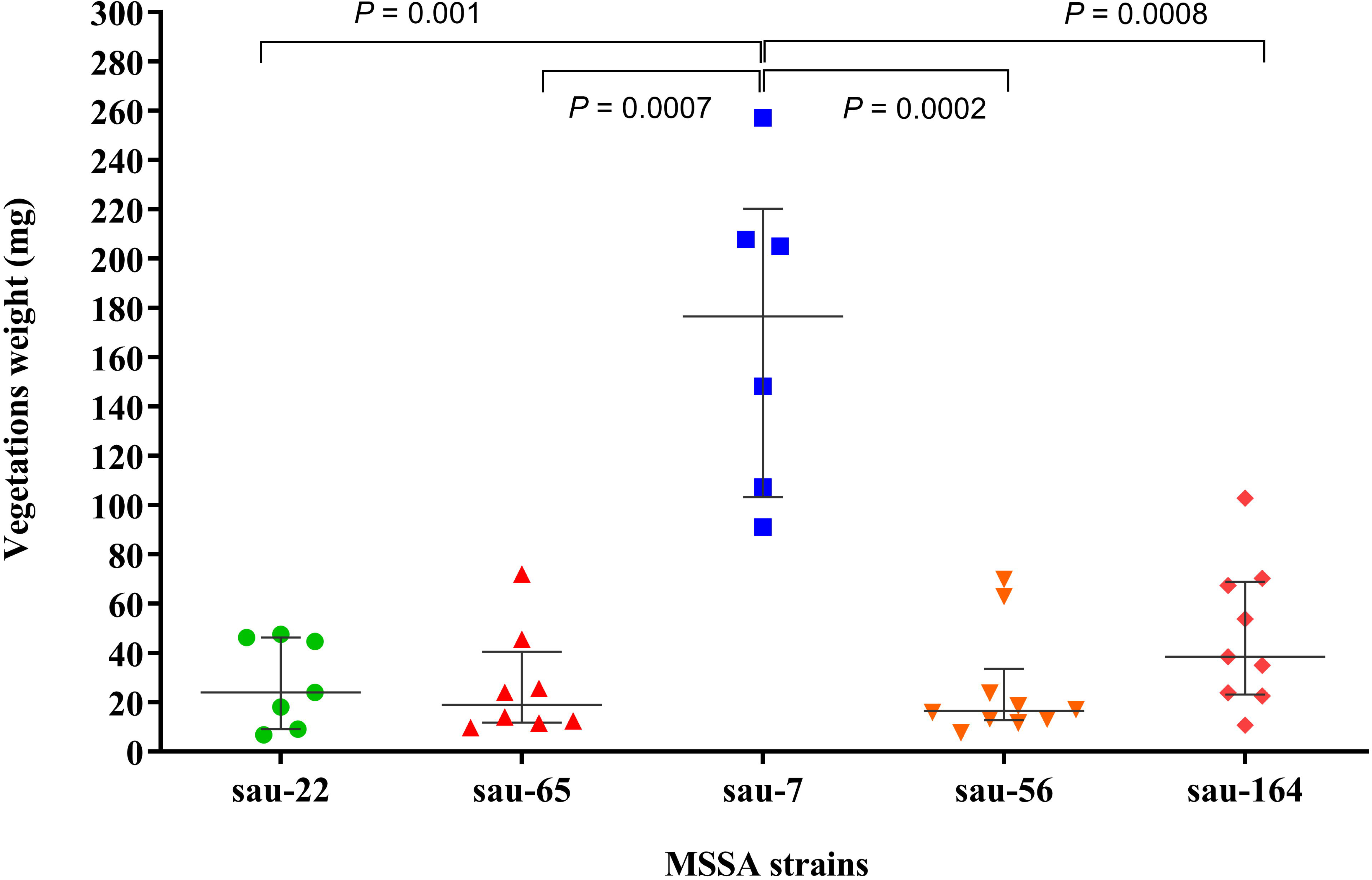
Aortic vegetation size in rabbits at the final time point. Aortic vegetation size in the IE model after infections with 10^6^ CFU/mL of *S. aureus* strains (Sau22, Sau65, Sau7, Sau56, and Sau164). Each dot represents an individual animal. Horizontal black bars indicate the mean and interquartile range of vegetation weight (mg).

### Comparative transcriptomics of CC30 and ST398 isolates

In order to elucidate the molecular mechanisms underlying the contrasting virulence displayed by CC30 and ST398 lineages in the animal models, global changes in gene expression between Sau22 and Sau65 (both CC30), Sau56 and Sau164 (both ST398) and Sau7 (ST398 with a truncated agrC) were assessed by RNA-seq. First, gene expression differences between strains of the same lineage were compared, excluding the Sau7 strain due to its distinct phenotypic behavior in animal models. Based on two-fold differences of gene expression, CC30 strains (Sau22 and Sau65) exhibited four DEGs (Table S3) and ST398 strains (Sau56 and Sau164) six DEGs (Table S4). These results indicate that strains within the same lineage exhibit very similar expression patterns. Based on these results, RNA-seq counts for the two CC30 isolates (Sau22 and Sau65), and the two ST398 isolates (Sau56 and Sau164) were treated as replicates in further DESeq2 analyses.

As depicted in Figure 7A, the RNA-seq study revealed 143 DEGs in *S. aureus* CC30 lineage when compared to ST398 with 75 up-regulated and 68 down-regulated genes (Table S5). GO enrichment analysis of the up-regulated genes was conducted to identify pathways putatively associated with enhanced lethality in ST398 isolates. This analysis identifies gene ontologies that are more prevalent than expected when compared to the entire genome. Specifically, we found that toxins were significantly enriched in the up-regulated gene set (fold enrichment = 16.7, FDR < 0.05), and included the *hly/hla* (SAOUHSC_01121, log2-FC = 4.6), *hld* (SAOUHSC_02260, log2-FC = 3.2) and *hlgB* (SAOUHSC_02710, log2-FC = 2.6) hemolysin-encoding genes. Other virulence factors were also found to be up-regulated in the ST398 isolates, including *agrB* (SAOUHSC_02261, log2-FC = 5.9), *agrC* (SAOUHSC_02264, log2-FC = 4.1), *scn* (SAOUHSC_02167, log2-FC = 3.5), *hysA* (SAOUHSC_02463, log2-FC = 3.3), *geh* (log2-FC = 2.2), *coa* (SAOUHSC_00192, log2-FC = 2.0), *harA* (SAOUHSC_01843, log2-FC = 1.5) and *ebp* (SAOUHSC_01501, log2-FC = 1.3) genes (Figure 7C). No significant enriched pathways were found in the ST398 down-regulated gene set.

**Figure 7.**
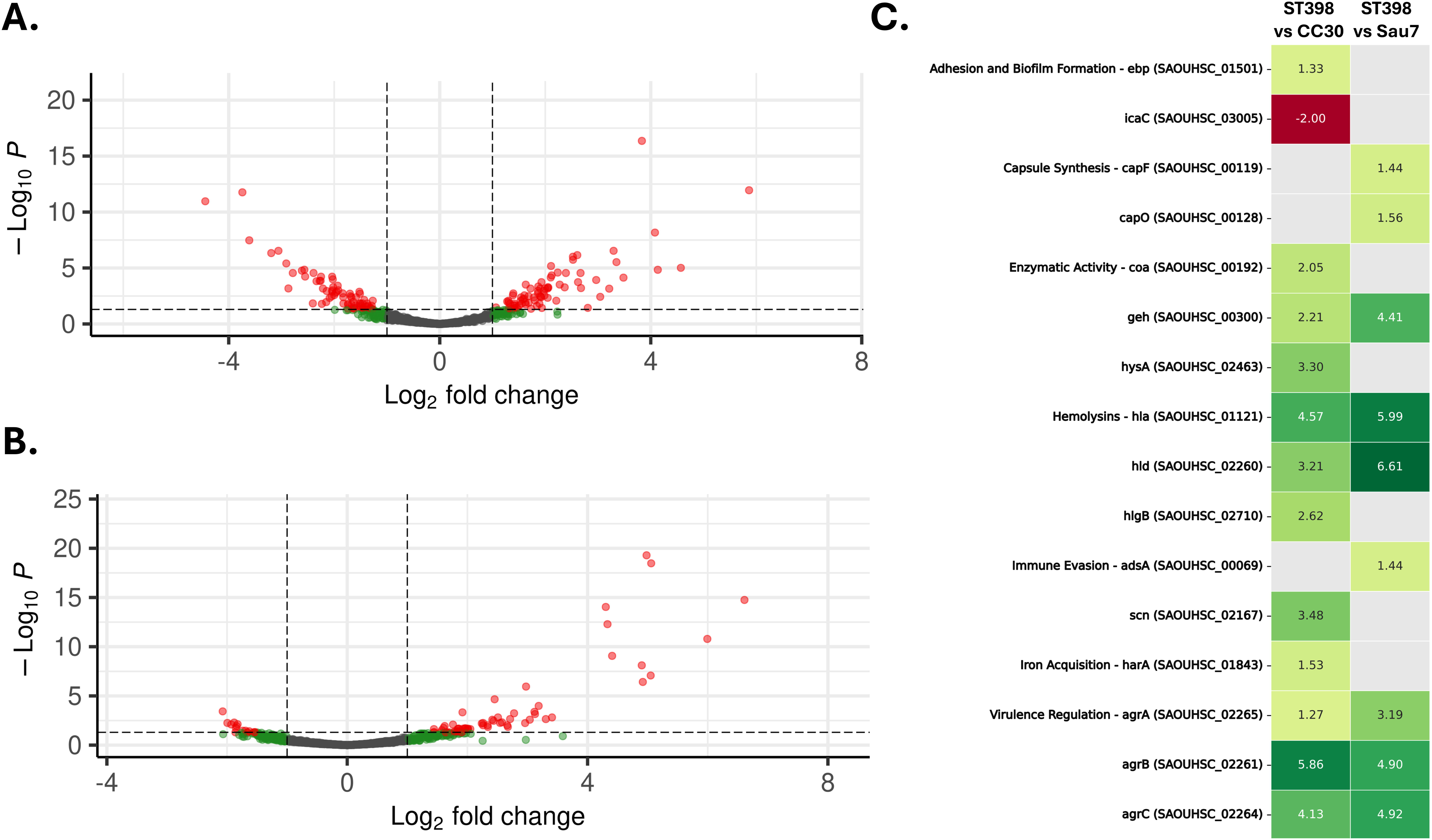
RNA-seq results overview. The figure includes volcano plots showing differentially expressed genes when comparing ST398 and CC30 isolates (A), and ST398 isolates and Sau7 strain (B). Each point on the plot represents a gene, with significantly differentially expressed genes (DEGs) highlighted in red (log2 fold-change ≥ ±1.0 and adjusted *p*-value < 0.05). Non-significant genes are shown in green (with a log2 fold-change between ±1.0, but no statistical significance) or gray (when no threshold is met). (C) Heatmap for differentially expressed virulence factor genes between ST398 and CC30 isolates, and between ST398 isolates and the Sau7 strain. The heatmap includes locus tags, preferred gene names, and virulence factor (VF) types. Non-significant genes are depicted as gray cells.

To gain insights into the distinctive lethality in *G. mellonella* larvae observed between *S. aureus* Sau7 (Figure S3) and the other ST398 strains, differential expression analysis was performed with DESeq2, comparing Sau7 to Sau56 and Sau164 strains. As shown in Figure 7B, the results revealed 77 DEGs, 62 of which were up-regulated and 15 down-regulated in the *S. aureus* ST398 strains (Table S6). GO analysis revealed that ST398 up-regulated genes significantly enriched pathways (fold enrichment = 37.3; FDR < 0.05) involved in quorum sensing, including *agrA* (SAOUHSC_02265, log2-FC = 3.2), *agrB* (log2-FC = 4.9), *agrC* (log2-FC = 4.9) and *agrD* (SAOUHSC_02262, log2-FC = 5.1) genes. Other virulence genes were also identified as up-regulated in *S. aureus* ST398 strains: *hld* (log2-FC = 6.6), *hly/hla* (log2-FC = 6.0), *geh* (SAOUHSC_00300, log2-FC = 4.4), *capO* (SAOUHSC_00128, log2-FC = 1.5), *capF* (SAOUHSC_00119, log2-FC = 1.4) and *asdA* (SAOUHSC_00069, log2-FC = 1.4) (Figure 7C). Again, no virulence genes were detected in the ST398 down-regulated genes set.

## DISCUSSION

*S. aureus* is a highly aggressive human pathogen and a leading cause of bacteremia and IE. The risk of complications and mortality associated with *S. aureus* infections is notably high; however, the underlying bacterial genetic factors contributing to these outcomes remain poorly understood. Consistent with previous studies, bioinformatics analyses conducted by our group revealed no significant correlation between the genetic features of a collection of 339 *S. aureus* isolates from bacteremic patients and the infection outcomes^16^. Host factors, including age, comorbidities, and the specific antibiotic treatments administered, significantly influence the progression of bacteremia^7^, likely complicating the identification of bacterial genetic markers associated with mortality. This limitation can be addressed using animal models that allow infection studies in genetically similar individuals under controlled conditions. The *G. mellonella* larvae infection model has emerged as a reliable and promising tool for studying *S. aureus* infections^33^.

### Lethality of *S. aureus* in *G. mellonella* larvae is partially aligned with mortality in patients with bacteremia

In this study, we observed notable discrepancies between the mortality rates associated with *S. aureus* infections in our cohort of bacteremic patients^16^ and the lethality observed in *G. mellonella* larvae. Specifically, methicillin-susceptible *S. aureus* (MSSA) ST398 exhibited the highest lethality in *G. mellonella*. However, the mortality associated with ST398 isolates in our patient cohort (30%) closely matched the average mortality rate across all patients (27.4%)^16^. These results suggest that the ST398 lineage demonstrates superior lethality in the *G. mellonella* model, indicating a genuine virulence potential for this clone. While previous studies have demonstrated the hypervirulence of MRSA-ST398 in animal models^34^, the lethality of MSSA-ST398 has been less well characterized. MSSA-ST398 is an emerging clone associated with severe infections, including bacteremia, IE, and bone and joint infections^35^. A previous study suggests that MSSA-ST398 strains exhibit greater virulence than non-ST398 strains, which is consistent with our findings^36^. Notably, evidence indicates that 30-day all-cause mortality was higher among patients with MSSA-ST398 bloodstream infections compared to those with non-ST398 MSSA. This suggests that a specific ST398 subtype may possess enhanced virulence, as reflected by its higher prevalence in bloodstream infection cases^36^. Interestingly, unlike the limited transmissibility of livestock-associated ST398-MRSA strains, MSSA-ST398 is readily transmitted between individuals^37^. The acquisition of prophages and other mobile genetic elements has driven the evolution of the ST398 lineage, contributing to its increased virulence and antibiotic resistance, making it a significant concern^38^.

Lineage CC1, which showed the highest mortality in our patient cohort (45.5%)^16^, also demonstrated high virulence in the *G. mellonella* model. CC1 is a predominant *S. aureus* clone involved in colonization, particularly among elderly nursing home residents, and is frequently linked to bloodstream infections^39^. Conversely, despite the relatively low mortality of the CC15 lineage in patients (15%)^16^, isolates from this lineage exhibited moderate lethality in *G. mellonella* larvae. On the other hand, lineages CC8 and CC22, which exhibited higher mortality in patients (40.0% and 37%, respectively)^16^, demonstrated intermediate lethality in *G. mellonella*. These lineages, along with CC5, are common MRSA clones known for their adaptability and capacity to acquire resistance genes, which allows them to persist in hospital settings and often affect patients with comorbidities^40^. This likely contributes to the elevated mortality rates observed in our patient cohort^41^.

The CC30 lineage exhibited the lowest lethality in the *G. mellonella* model, which was consistent with the low mortality observed in bacteremic patients infected with CC30 (21.2%)^16^ and its absence of relationship with 1-year mortality in MSSA IE^11^. Previous studies have reported the absence of sepsis-induced mortality in murine and *G. mellonella* models for CC30 isolates^42^, suggesting that this lineage elicits relatively low levels of pathogenicity compared to other clones^43^. A recent study on IE patients in Spain reported that CC30 is associated with persistent bacteremia (≥5 days) and inversely correlated with mortality^41^. However, some reports indicate that CC30 isolates have acquired metabolic traits that contribute to persistent and complicated infections, including IE^42^. Although the underlying mechanism linking CC30 to IE remains unclear, it has been suggested that specific characteristics of CC30 contribute to an attenuated immune response, facilitating bloodstream entry and prolonged persistence^44^.

Considering all lineages, the mean LT50 values for isolates from fatal and non-fatal bacteremia cases did not differ significantly in the *G. mellonella* model (MWU, p > 0.05), reinforcing the discrepancies in *S. aureus* pathogenesis between humans and the *G. mellonella* model (Figure 8A). Similarly, we observed no correlation between the phenotypic characteristics of *S. aureus* and mortality in humans. Notably, isolates with the highest biofilm-forming capacity were consistently non-fatal (Figure 8B). In contrast to our findings in *G. mellonella*, no correlation was observed between growth rate, hemolytic type, and mortality in our patient cohort (Figure 8CD).

**Figure 8.**
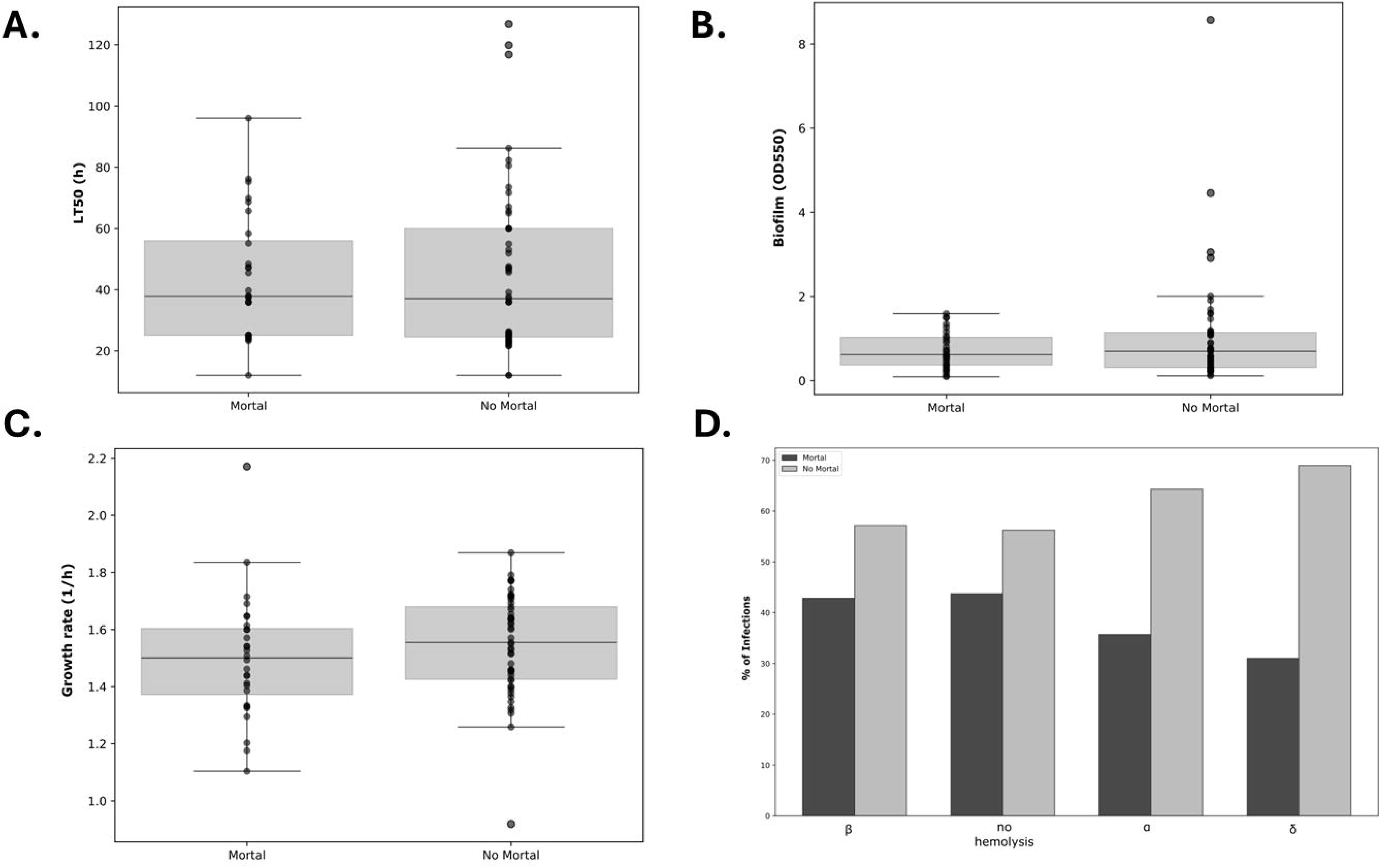
Correlation between mortal and non-mortal bacteremia in human patients and bacterial characteristics, including LT_50_ values in *G. mellonella* larvae (A), biofilm formation (B), growth rate (C), and hemolysis type (D).

### Role of the *agr* system in *S. aureus* virulence and mortality in *G. mellonella*

Our study did not reveal any significant correlation between the growth rate of *S. aureus* isolates or their ability to form biofilm and their lethality in *G. mellonella* larvae (Figure 4AB). In contrast, α- and δ-hemolysis were associated with an increased risk of mortality in *G. mellonella* (Figure 4C), while the presence of a type III *agr* system correlated with reduced virulence in *G. mellonella* (Figure 4D). It is well established that the *agr* system connects quorum sensing to the production of cytolytic toxins^45^. When the *agr* system is active, RNAIII is expressed, which regulates various virulence factors, including secreted toxins such as α- and δ-hemolysins, phenol-soluble modulins, and leucocidins like Panton-Valentine leucocidin^46^. Notably, α-hemolysin has been identified as a key factor contributing to mortality in pneumonia caused by MSSA-ST398^47^. Consistent with this, ST398 isolates exhibited significantly higher *in vitro* expression levels of α-toxin and δ-toxin compared to non-ST398 isolates, and MSSA-ST398 displayed a hypervirulent phenotype in mice^47^. Other studies have also highlighted the high hemolytic activity of the ST398 lineage as a hallmark of community-associated clones^48^.

Despite the high lethality of the ST398 lineage, disruption of the *agrC* gene in the Sau7 isolate led to an attenuated phenotype. Notably, biofilm formation in Sau7 (1.593) was significantly higher compared to the ST398 average (0.616 ± 0.537), while hemolytic activity was abolished. Similarly, inactivation of *agrC* in Sau58 and Sau40, both from the CC22 lineage, resulted in reduced lethality, increased biofilm formation, and a loss of hemolytic activity. However, *agrC* inactivation did not influence the bacteremia outcomes in the patients from our cohort^16^. In contrast, double *agrBC* disruption, observed in Sau1 (CC45) and Sauc6 (CC1) isolates, did not reduce virulence in *G. mellonella*. Furthermore, biofilm formation and hemolysis remained unchanged in Sau1 and Sauc6, suggesting that *agrB* inactivation might compensate for a*grC* disruption. The prevalence and potential impact of double *agrBC* disruption on infection outcomes in *S. aureus* remain areas for further investigation.

### Disruption of *agrC* and its role in *S. aureus* virulence across different infection models

Sau22 (CC30) was the least virulent isolate in the EE rabbit model. Similarly, two animals in the Sau65-infected (CC30) group did not develop endocarditis, despite showing initial bacteremia, suggesting that the Sau65 strain has lower infectivity. These findings are consistent with previous reports that describe low CC30 virulence in vertebrate models^42^ and align with the results observed in *G. mellonella* (Figure 3). In contrast, the three ST398 isolates tested exhibited similar survival rates in the EE model, which differs from the findings in *G. mellonella*, where the Sau7 strain demonstrated reduced lethality (Figure 3). This suggests that the factors influencing infection outcomes in the EE model in rabbits may differ from those in *G. mellonella* larvae.

Interestingly, Sau7 carries a disrupted *agrC* gene. Transposon insertions in the *agrC* gene have been reported in other *S. aureus* clinical isolates, suggesting that such inactivation may be driven by pathoadaptive processes^49^. *agrC* mutations, including inactivating variants, have frequently been identified during *S. aureus* colonization^50^. Notably, recent reports have linked *agrC* gene inactivation to persistent *S. aureus* infections^51^. In line with these findings, Sau7 was isolated from a patient with persistent bacteremia, hematogenous seeding, and spondylodiscitis. This patient had severely impaired functional reserve and a progressing lung neoplasm despite multiple chemotherapeutic regimens^16^.

Moreover, in the EE rabbit model, we observed high *S. aureus* Sau7 counts in peripheral organs and larger vegetation size, which may indicate enhanced colonization potentially linked to *agr* dysfunction due to *agrC* deficiency. This finding suggests that increased vegetation formation results in a more compact structure with less fragmentation, potentially decreasing the risk of septic embolism, as seen in human cases^52,53^. This observation is consistent with a recent study showing that vegetation formation inversely correlates with *agr* system expression in an EE rabbit model, where strains proficient in vegetation formation exhibit lower RNAIII expression^54^. The larger vegetations observed in the rabbits EE model may explain the increased lethality of Sau7 in this model, which was not seen in the *G. mellonella* model, likely due to the anatomical limitations of *G. mellonella* in developing endocarditis.

### Enhanced expression of virulence factors in isolates of the ST398 lineage

Although CC30 and ST398 isolates share a similar set of virulence factors, transcriptome analyses revealed notable differences between these two lineages (Figure 7A). ST398 strains exhibited increased expression of virulence factors, particularly those encoding hemolysins, which likely contribute to their superior pathogenic potential observed in animal models. These transcriptional patterns were consistent when comparing ST398 to the *agrC*-deficient isolate Sau7 (Figure 7B).

In ST398 isolates, genes from the *agr* operon were notably up-regulated. The *agr* system is a key regulator of virulence in *S. aureus*, with its role well-documented in animal infection models^55,56^. This quorum sensing mechanism controls the expression of approximately 200 genes, with RNAIII acting as the primary effector by regulating gene expression through antisense pairing^57^. Specifically, RNAIII positively regulates the expression of *hla*, which encodes α-hemolysin^58^, and was significantly overexpressed in ST398 isolates. The importance of α-hemolysin in *S. aureus* pathogenicity is widely recognized in animal models^59^. α-Hemolysin is a pore-forming toxin that disrupts cell membranes by altering transmembrane ion gradients, leading to cell lysis and tissue damage^60^. It also plays a role in immune activation and severe infection^61^. Additionally, RNAIII encodes δ-hemolysin, which exhibits both hemolytic and antimicrobial activities^62^. The reduced expression of the *agr* system in CC30 isolates and in the Sau7 is consistent with the observed decrease in both α- and δ-hemolysin expression and their diminished *in vitro* activity (Figure 9). However, the role of δ-hemolysin in *S. aureus* pathogenicity within the *G. mellonella* model remains underexplored. The *agr* system also regulates the *geh* gene^63^, which encodes a lipase critical for hydrolyzing host lipids, releasing fatty acids essential for bacterial metabolism^64^. Beyond the *agr* system, ST398 strains up-regulated additional virulence genes compared to Sau7 and CC30 isolates, suggesting the involvement of alternative regulatory mechanisms^63,65^. These include genes related to tissue invasion (*hysA*, *coa*), adhesion and immune modulation (*ebp*, *capO*, *capF*), immune evasion (*scn*), nutrient acquisition (*harA*), and metabolic processes (*asdA*).

**Figure 9.**
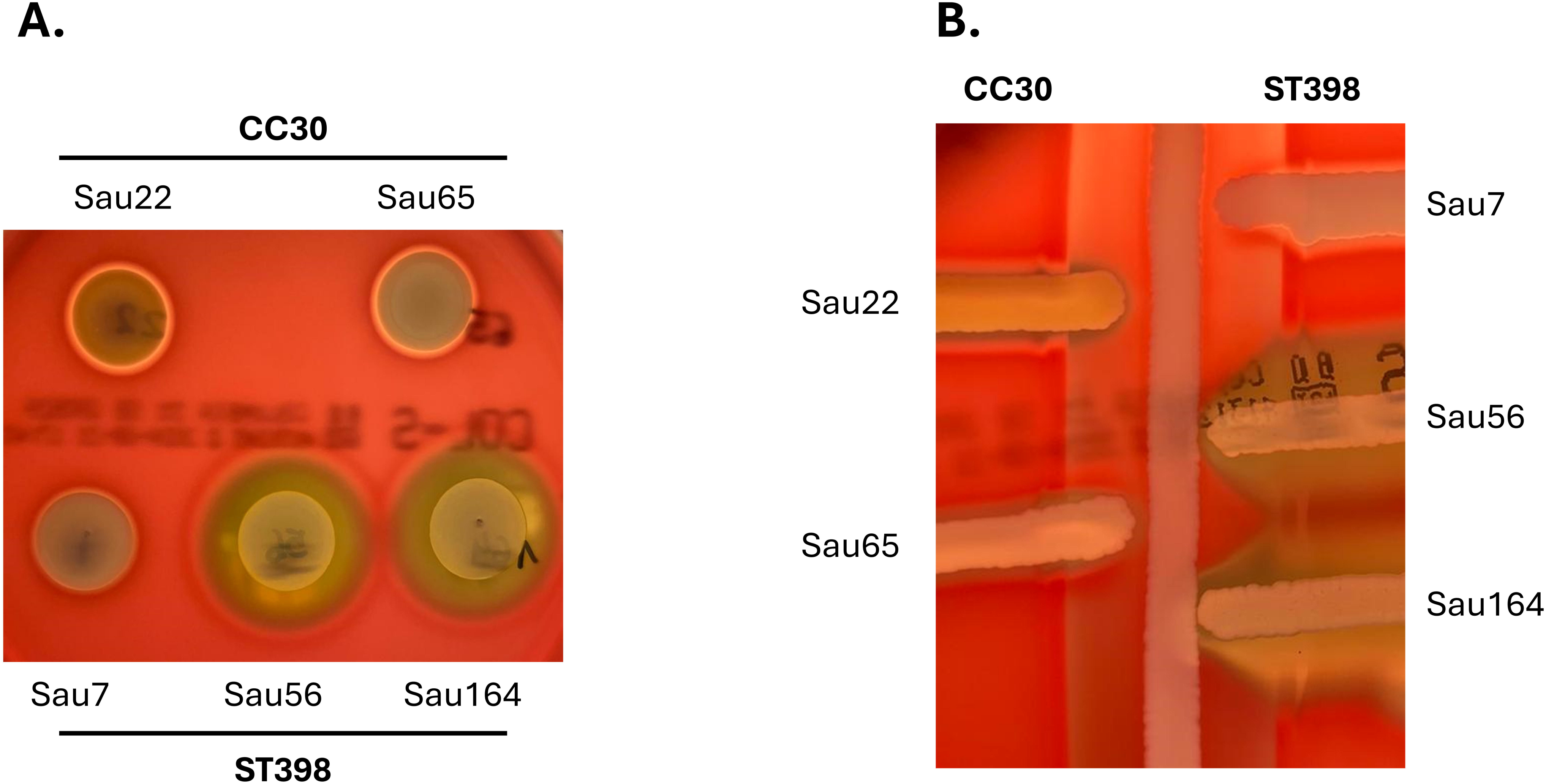
Hemolysis activity assay results for *S. aureus* isolates tested in the IE rabbit model and RNA-seqs. Hemolysis activities are shown as 10 µl drops (A) and CAMP assays (B) in Columbia agar plates.

This study underscores the variability in *S. aureus* virulence across different experimental models and human patients. While certain *S. aureus* lineages demonstrated enhanced virulence in these models, particularly through up-regulation of key virulence factors, these findings did not always correlate with patient outcomes. These discrepancies highlight the complexity of *S. aureus* pathogenesis and suggest that host-specific factors may play a critical role in shaping infection outcomes, emphasizing the need for further exploration of how experimental model results translate to human disease.

## ACKNOWLEDGMENTS

This work was supported by grant PI24/01294 from the Instituto de Salud Carlos III (ISCIII) and co-funded by the European Union. M.S.-O. is the recipient of a Margarita Salas fellowship from the Spanish Ministerio de Universidades. M.A.C received a Sara Borrell personal research grant (CD21/00125, 2022-24) from ISCIII. O.G. is the recipient of a research grant from the “Pla Estratègic de Recerca i Innovació en Salut (PERIS)” awarded by the Departament de Salut de la Generalitat de Catalunya. M.B., A.-C.G., I.G. and D.Y also acknowledge the funding from the Spanish MICINN (PID2023-150290OB-I00) and the Catalan AGAUR (2021 SGR00092). J.M.M. received a personal 80:20 research grant from Institut d’Investigacions Biomèdiques August Pi i Sunyer (IDIBAPS; Barcelona, Spain) during 2017–2025. We thank Dr. D. Scott Merrell (University of Arizona) for his critical review of the manuscript. We are grateful to SeqCenter (Pittsburgh, USA) for RNA sequencing. We also want to acknowledge the CERCA Programme / Generalitat de Catalunya.

## AUTHOR CONTRIBUTIONS

Conceptualization, M.S.-O., D.Y. and O.Q.P.; Investigation, M.S.-O., M.B., M.A.C., A.-C.G., I.G.-S. and C.G.-M.; Software, M.S.-O.; Formal Analysis, M.S.-O., M.B., M.A.C.; Resources, J.M.M., I.G, O.G., C.G.-M., D.Y. and O.Q.P.; Writing – original draft preparation, M.S.-O. and O.Q.P.; Writing – review and editing, M.S.-O., M.B., M.A.C., A.-C.G., I.G.-S., J.M.M., I.G., O.G., C.G.-M., D.Y. and O.Q.P; Funding acquisition, D.Y. and O.Q.P. All authors have read and agreed to the published version of the manuscript.

## Supplementary Material

**Data S1.** JSON-formatted file including the clinical data and molecular features for all *S. aureus* strains included in this study.

**Figure S1.** CAMP hemolysis results for all tested *S. aureus* isolates.

**Figure S2.** Kaplan-Meier survival curves for all tested *S. aureus* isolates.

**Figure S3.** (A) Graphical representation of *agr* operon in *S. aureus* ST398 isolates. (B) PCR verification showing the transposon insertion within the *agrC* gene in the Sau7 isolate.

**Figure S4.** Results of bacterial growth (CFU/g) of the selected MSSA isolates in vegetations (A), spleen (B), kidney (C) and brain (D) in a rabbit model of experimental endocarditis. Significant *p*-values are indicated.

**Table S1.** Oligonucleotides used in this work.

**Table S2.** Phenotypic characterization of all *S. aureus* strains analyzed in this study, including LT_50_, biofilm formation, growth rate and doubling time.

**Table S3.** Differentially expressed genes (DEGs) in Sau22 when compared to Sau56. EggNOG functional annotation is provided for each DEG.

**Table S4.** Differentially expressed genes (DEGs) in Sau56 when compared to Sau164. EggNOG functional annotation is provided for each DEG.

**Table S5.** Differentially expressed genes (DEGs) in Sau56 and Sau164 (ST398) when compared to Sau22 and Sau56 (CC30) isolates. EggNOG functional annotation is provided for each DEG.

**Table S6.** Differentially expressed genes (DEGs) in Sau56 and Sau164 (ST398) when compared to the Sau7 isolate. EggNOG functional annotation is provided for each DEG.

