## Supplementary material for "Comparative mortality of dominant *Staphylococcus aureus* lineages in human bacteremia and animal infection models": Figure S1

**Sau1 (CC45)**

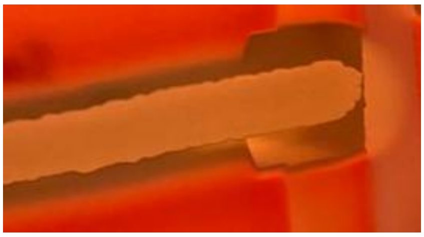

**Sau2 (ST398)**

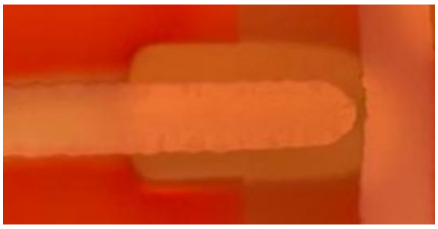

**Sau7 (ST398)**

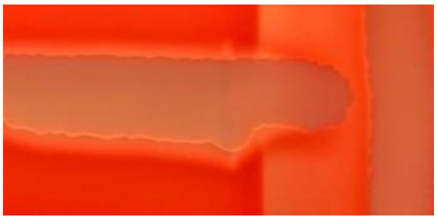

**Sau12 (CC8)**

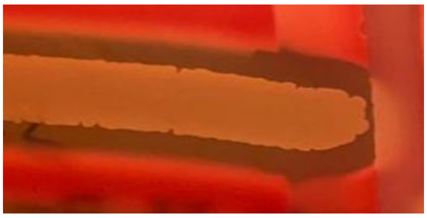

**Sau20 (CC8)**

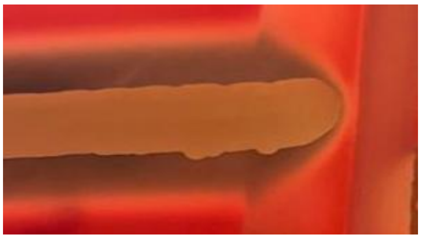

**Sau22 (CC30)**

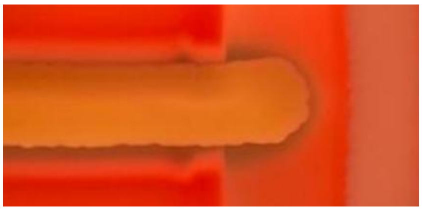

**Sau31 (CC5)**

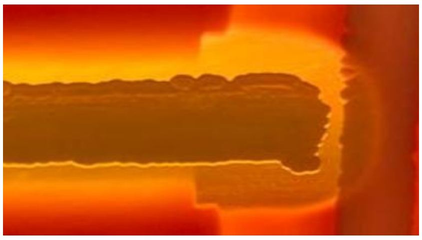

**Sau33 (CC15)**

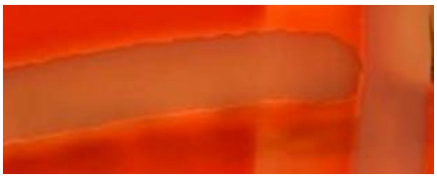

**Sau35 (CC45)**

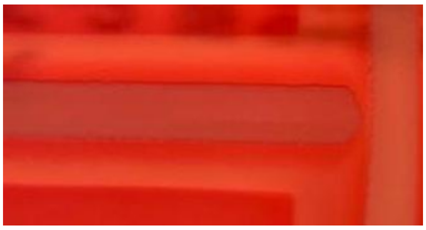

**Sau37 (CC5)**

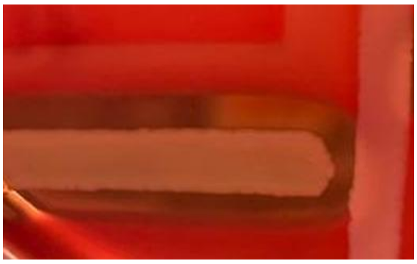

**Sau38 (CC45)**

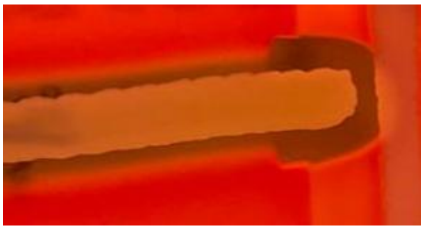

**Sau40 (CC22)**

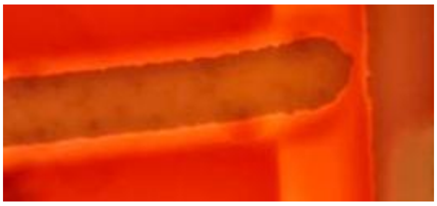

**Sau46 (CC5)**

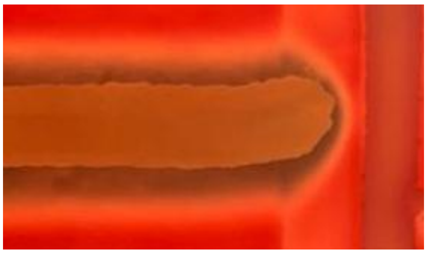

**Sau47 (CC8)**

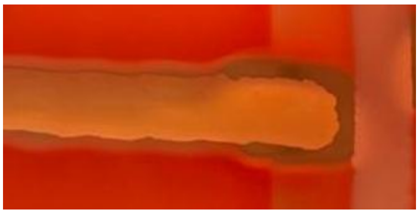

**Sau48 (ST291)**

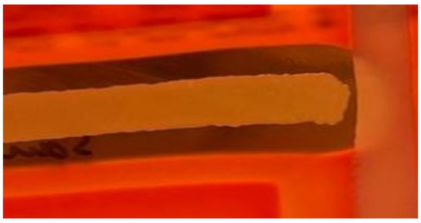

**Sau54 (CC15)**

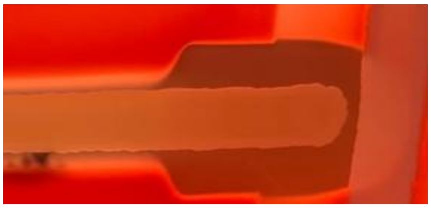

**Sau55 (CC5)**

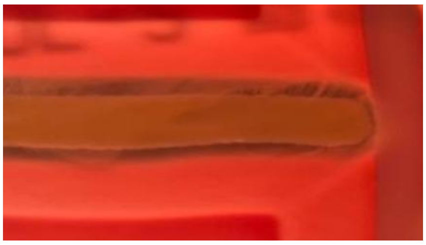

**Sau56 (ST398)**

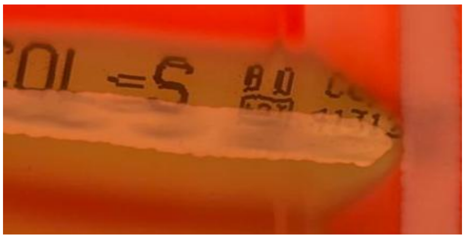

**Sau58 (CC22)**

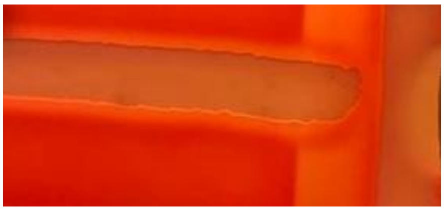

**Sau59 (CC15)**

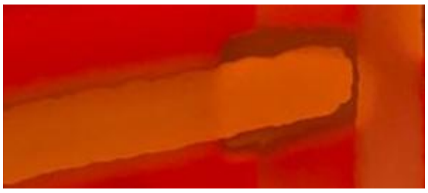

**Sau60 (CC5)**

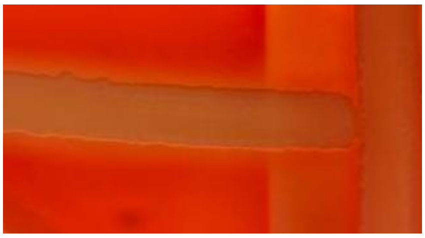

**Sau62 (CC30)**

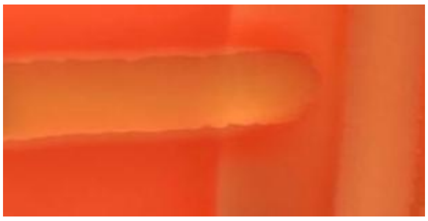

**Sau65 (CC30)**

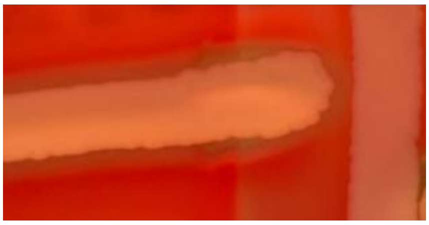

**Sau66 (CC30)**

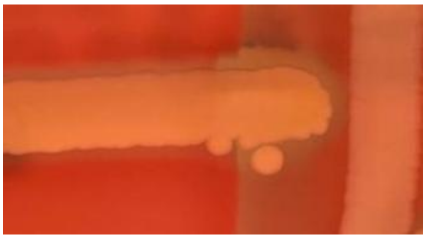

**Sau69 (CC5)**

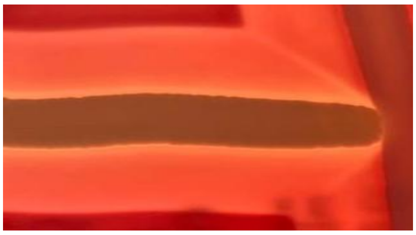

**Sau71 (CC5)**

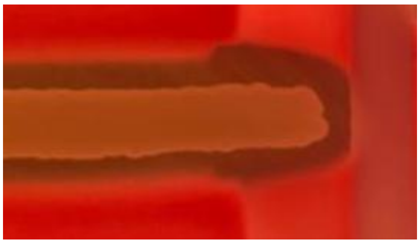

**Sau72 (CC8)**

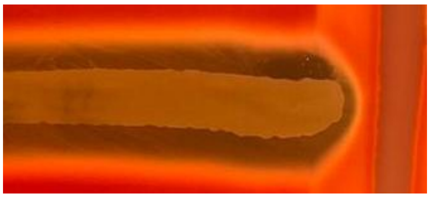

**Sau74 (CC1)**

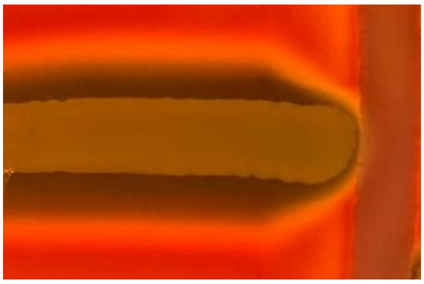

**Sau75 (CC45)**

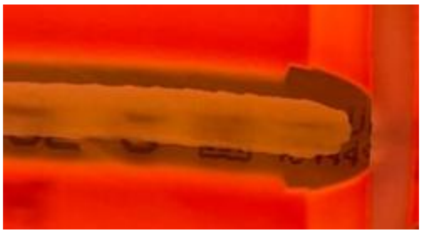

**Sau79 (CC22)**

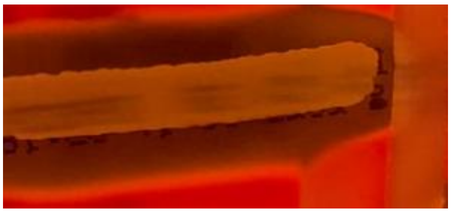

**Sau81 (CC5)**

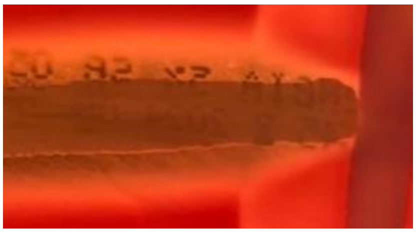

**Sau84 (ST398)**

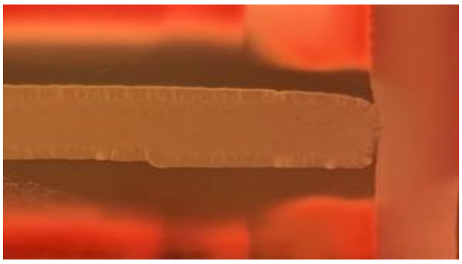

**Sau85 (CC30)**

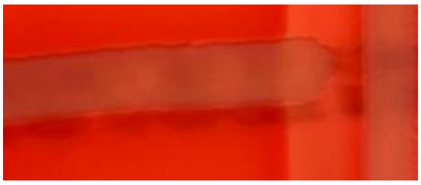

**Sau88 (CC30)**

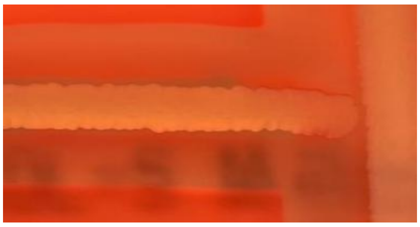

**Sau89 (CC30)**

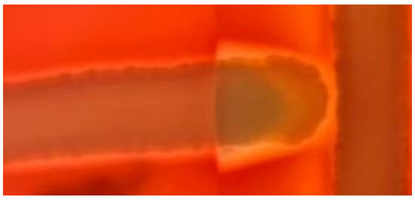

**Sau91 (CC22)**

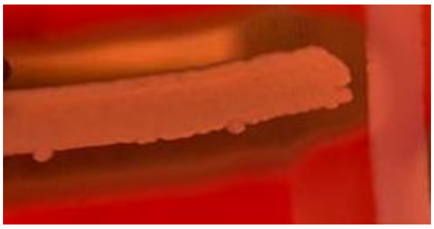

**Sau92 (CC121)**

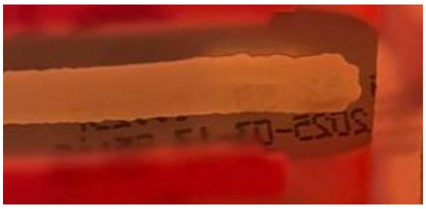

**Sau94 (ST291)**

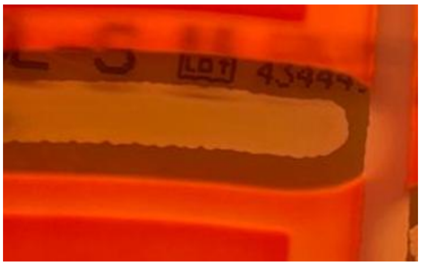

**Sau96 (CC45)**

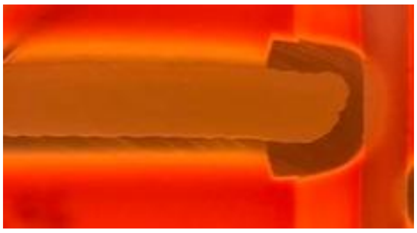

**Sau99 (CC1)**

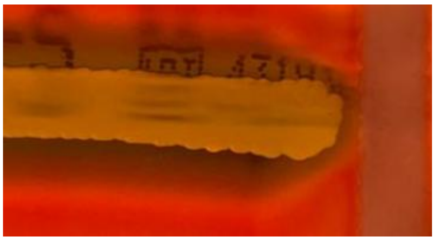

**Sau100 (CC5)**

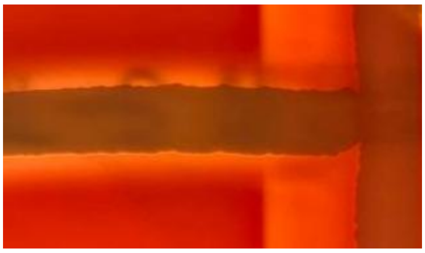

**Sau101 (ST398)**

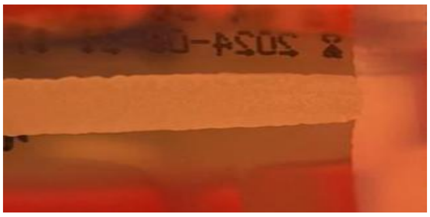

**Sau103 (ST7)**

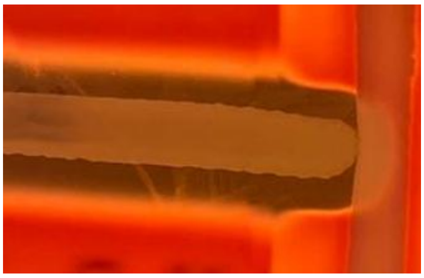

**Sau107 (ST7)**

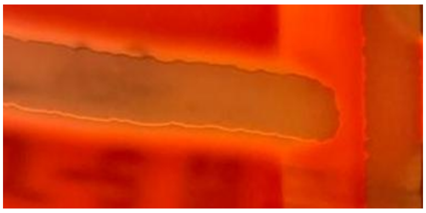

**Sau108 (CC8)**

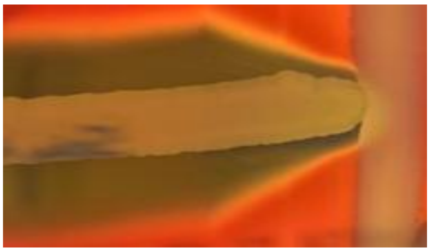

**Sau112 (CC45)**

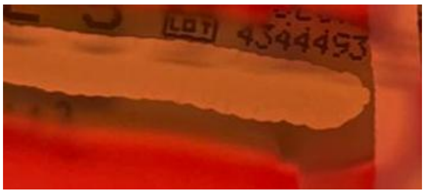


**Sau115 (ST398)**

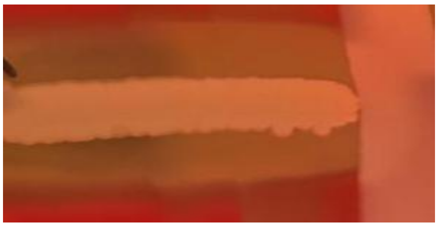

**Sau116 (ST398)**

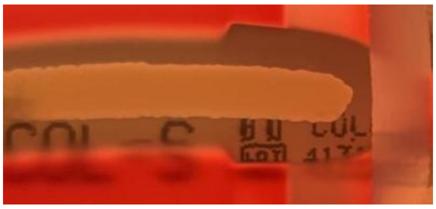


**Sau119 (CC30)**

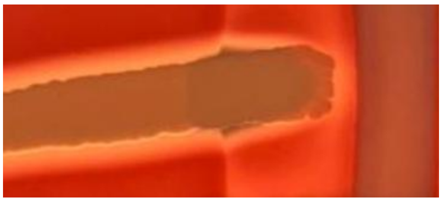

**Sau123 (CC15)**

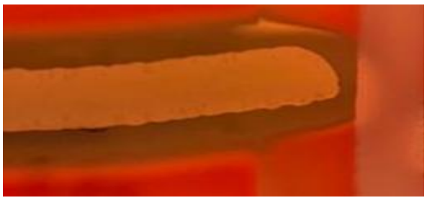

**Sau124 (CC30)**

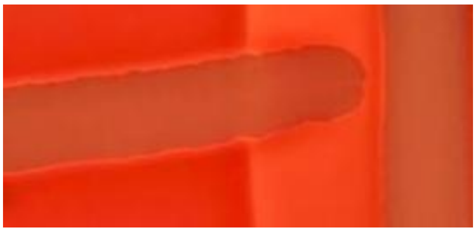

**Sau132 (CC30)**

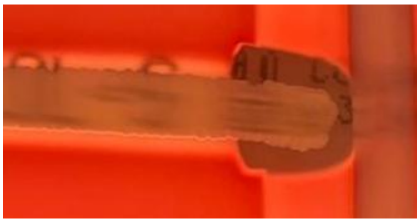

**Sau133 (CC30)**

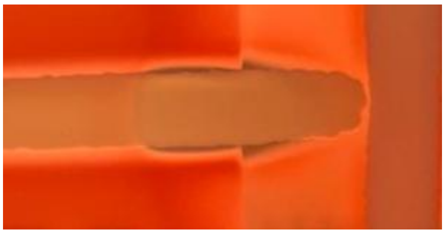

**Sau135 (ST398)**

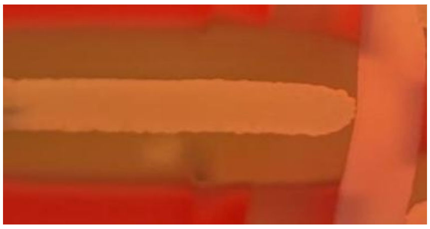

**Sau137 (CC22)**

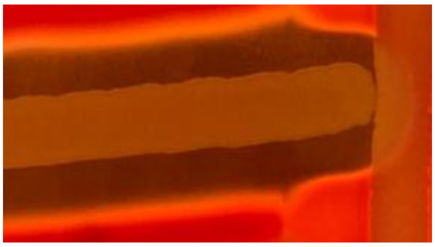

**Sau139 (ST398)**

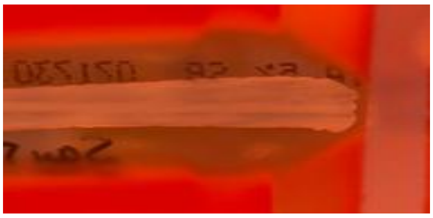

**Sau146 (CC30)**

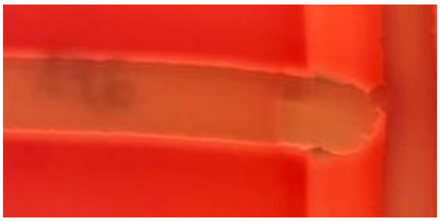

**Sau147 (CC1)**

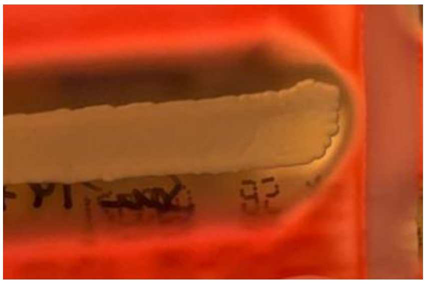

**Sau148 (CC45)**

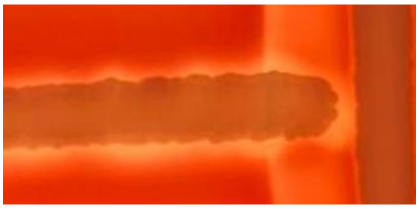

**Sau149 (CC30)**

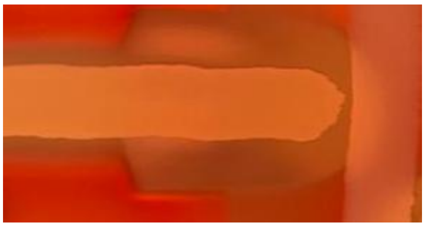

**Sau150 (CC8)**

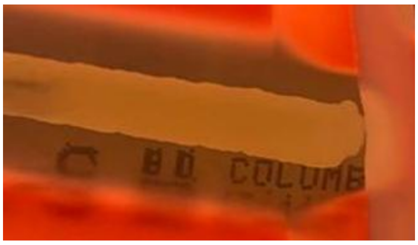

**Sau152 (CC121)**

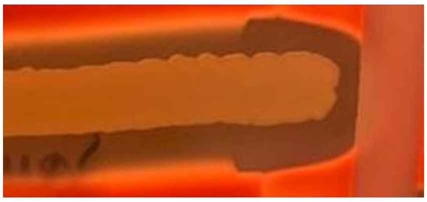

**Sau153 (CC30)**

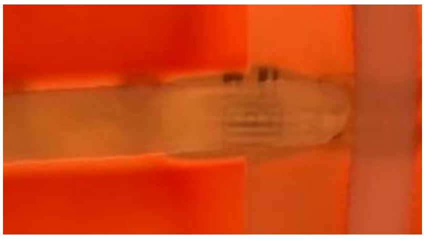

**Sau158 (CC8)**

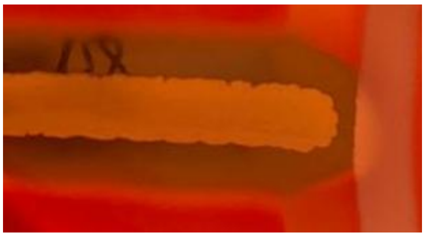

**Sau162 (ST7)**

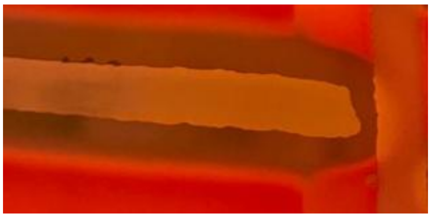

**Sau163 (CC8)**

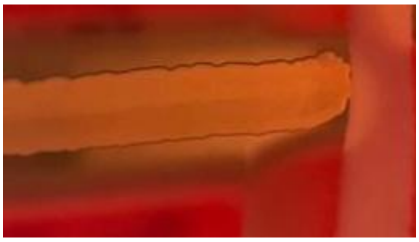

**Sau164 (ST398)**

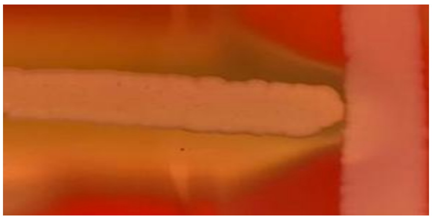

**Sau165 (CC45)**

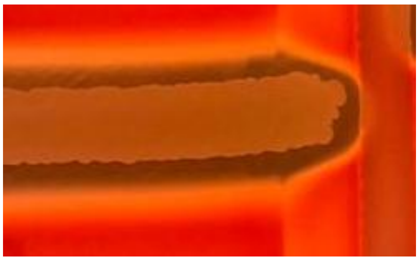

**Sau166 (CC5)**

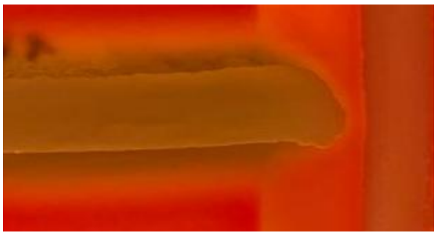

**Sau171 (CC1)**

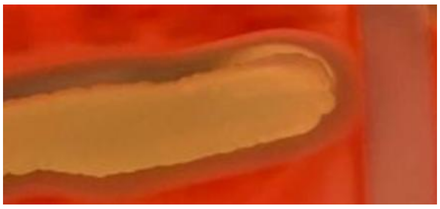

**Sau188 (CC1)**

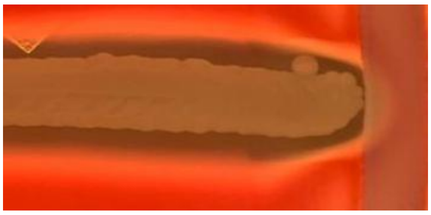

**Sau197 (CC1)**

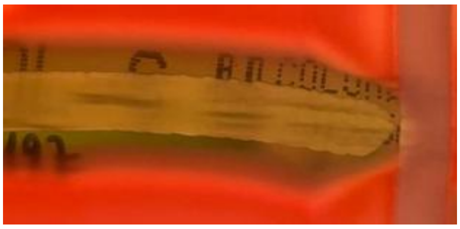

**Sau210 (CC1)**

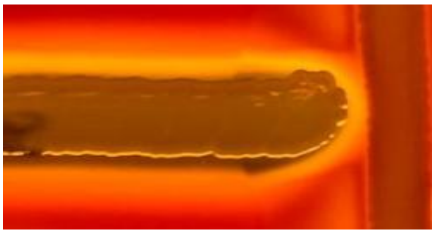

**Sau229 (CC1)**

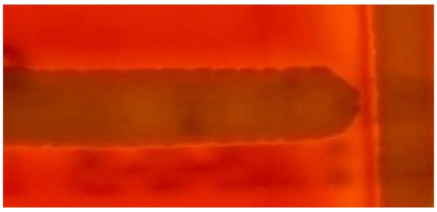

**Sau275 (CC1)**

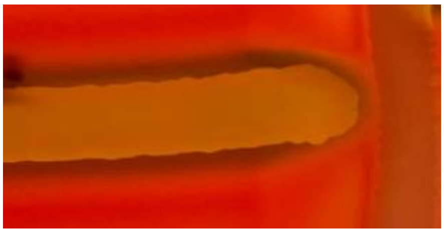

**Sau360 (CC1)**

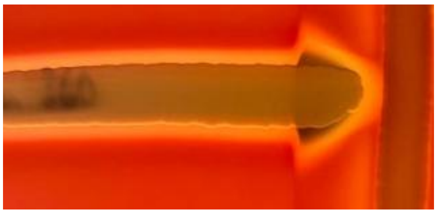

**Sauc6 (CC1)**

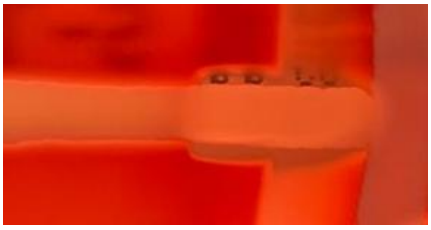
