## Supplementary material for "Comparative mortality of dominant *Staphylococcus aureus* lineages in human bacteremia and animal infection models": Figure S2

Sau1 (CC45)

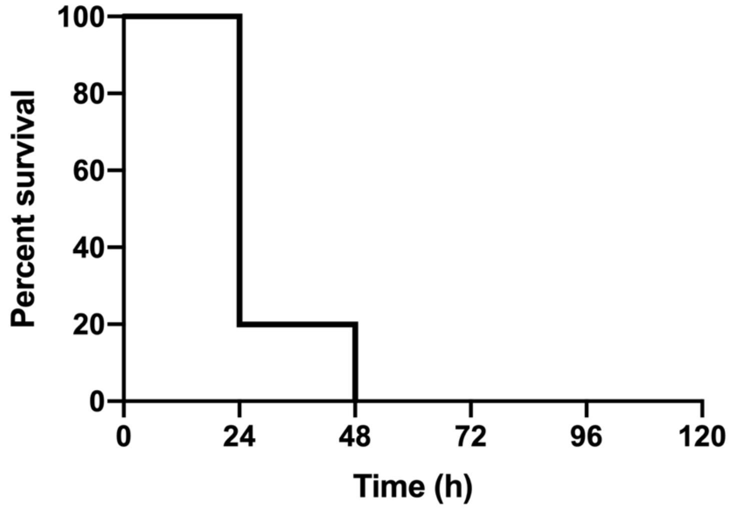


Sau2 (ST398)

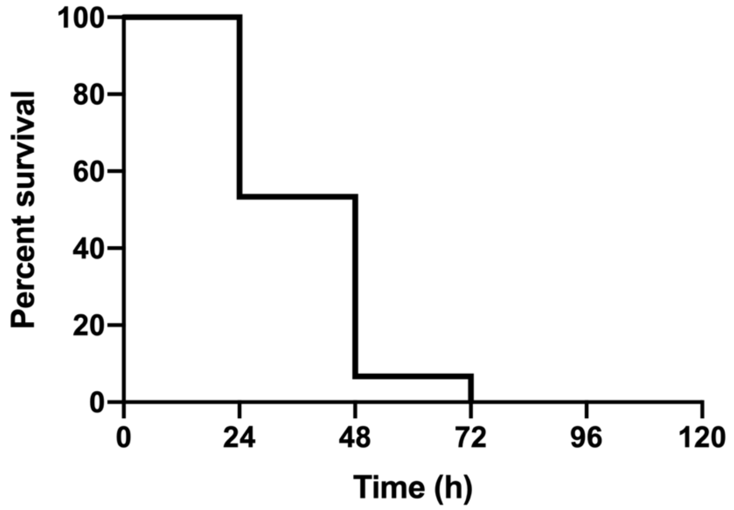


Sau7 (ST398)

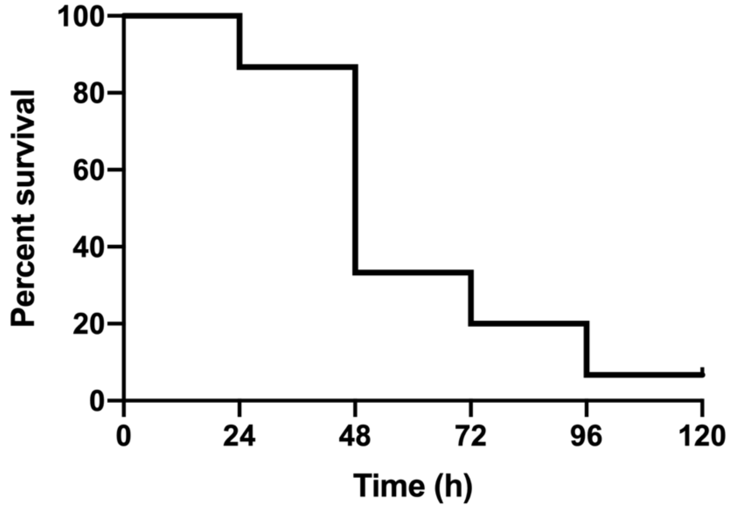

Sau12 (CC8)

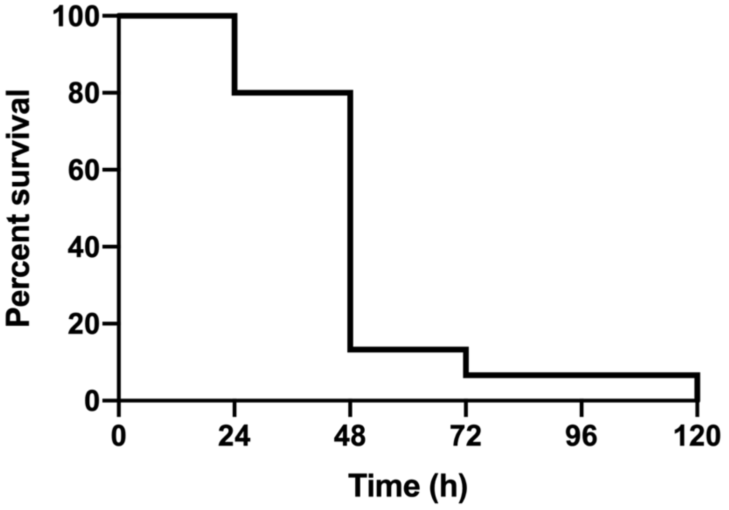


Sau20 (CC8)

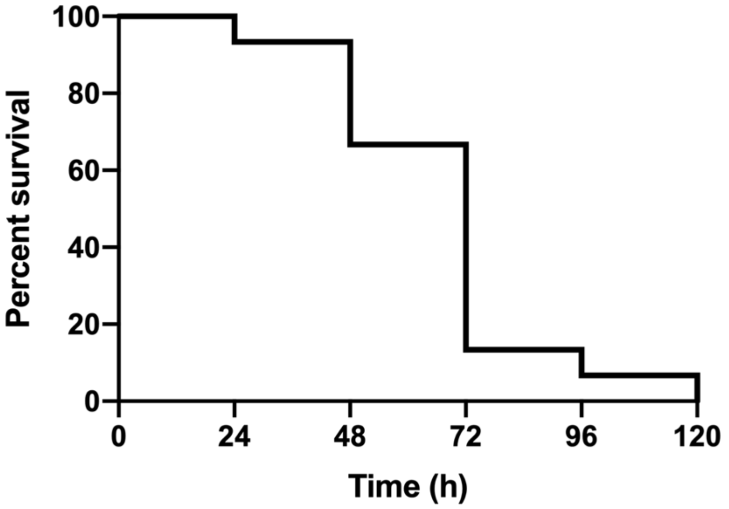


Sau22 (CC30)

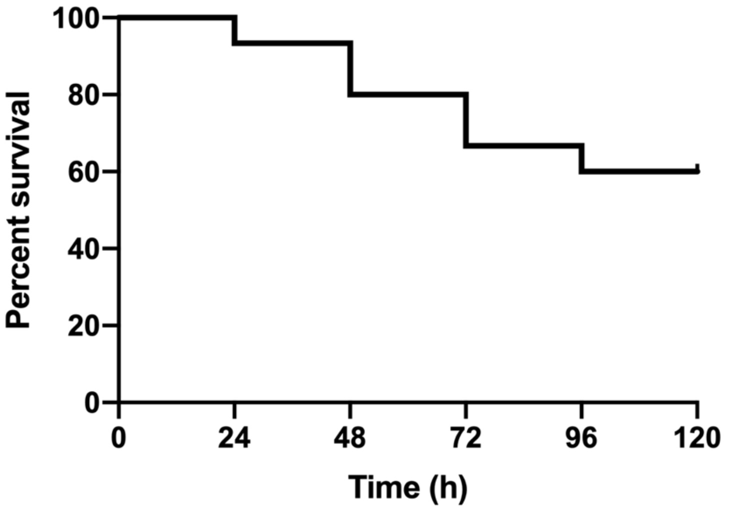

Sau31 (CC5)

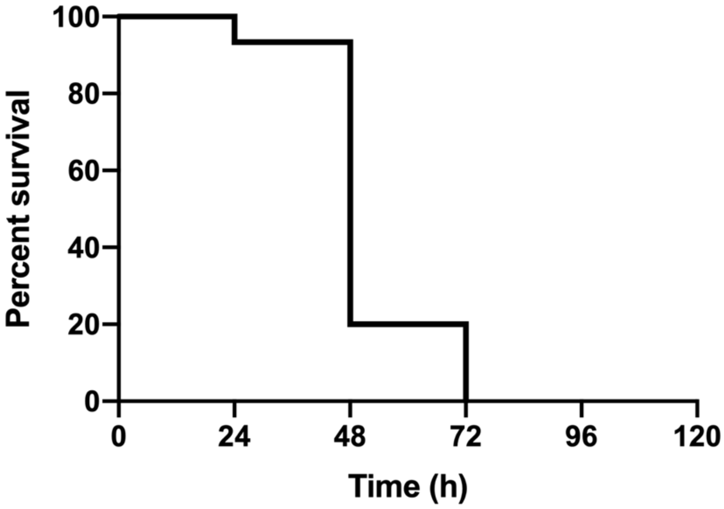


Sau33 (CC15)

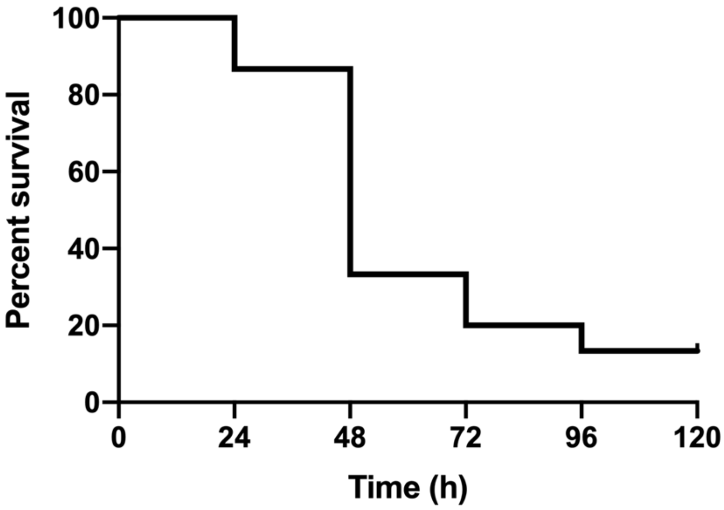


Sau35 (CC45)

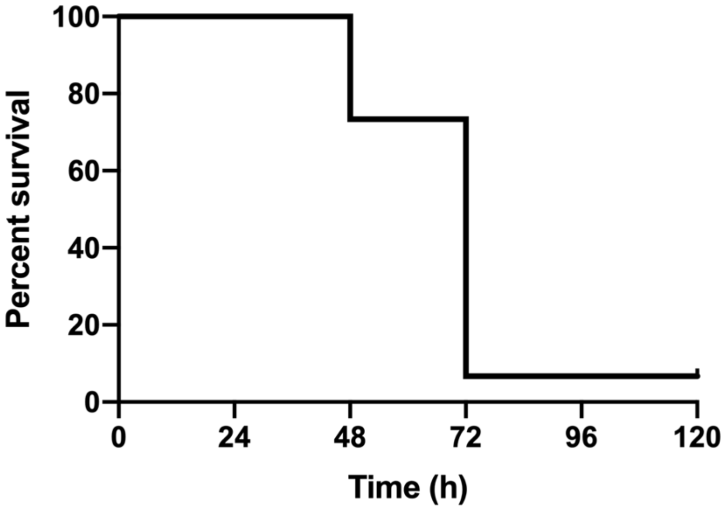

Sau37 (CC5)

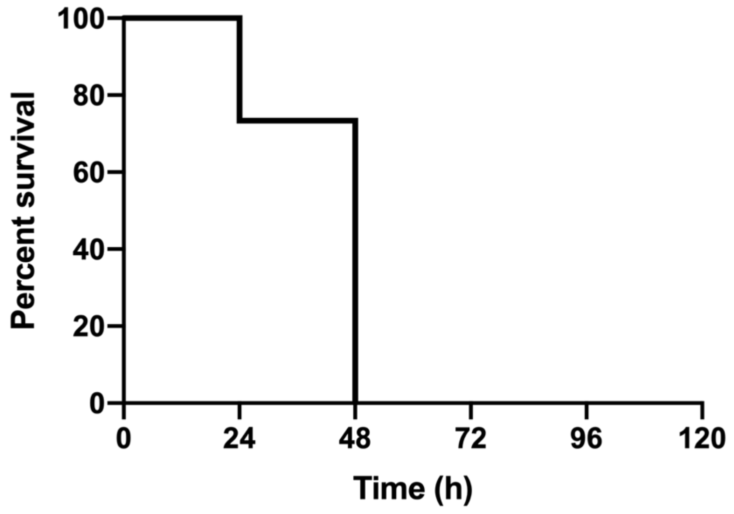


Sau38 (CC45

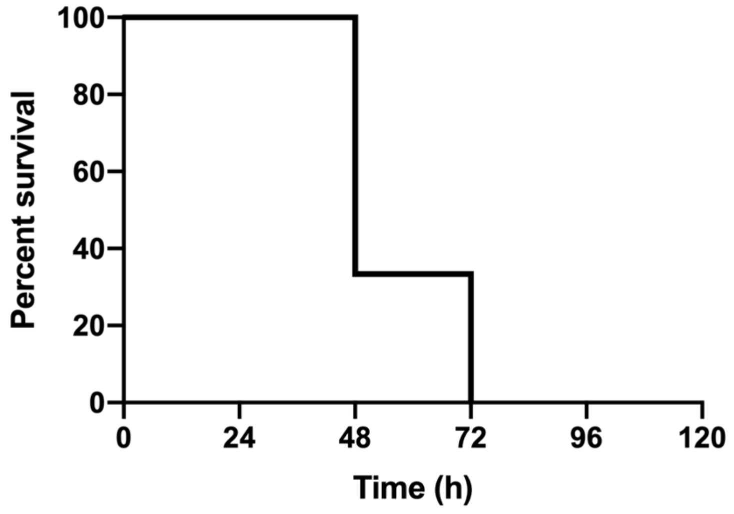


Sau40 (CC22)

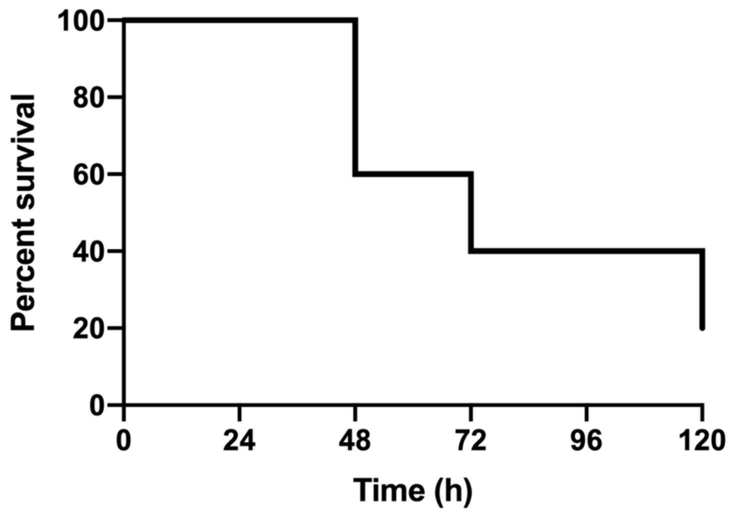

Sau46 (CC5)

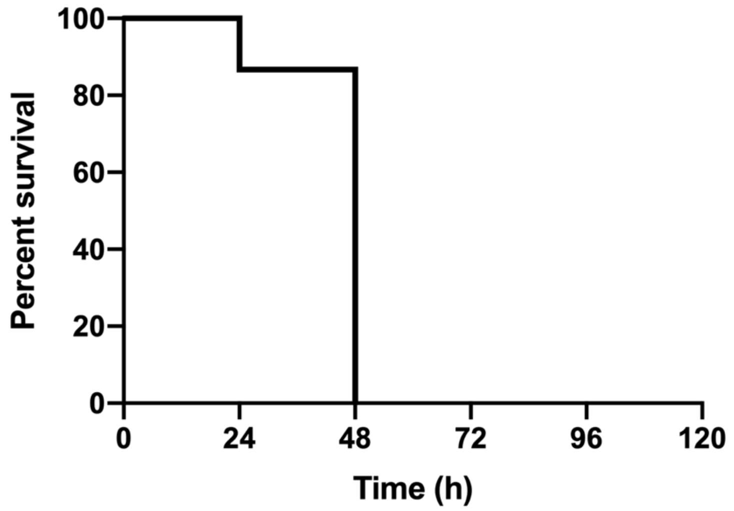


Sau47 (CC8)

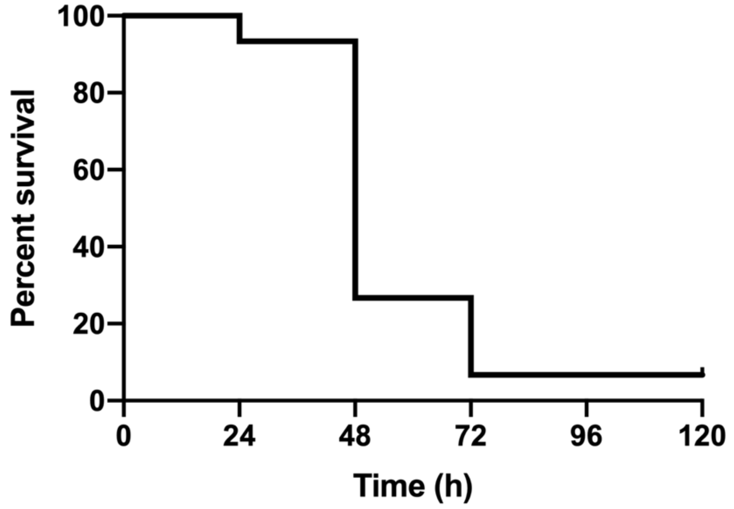


Sau48 (ST291)

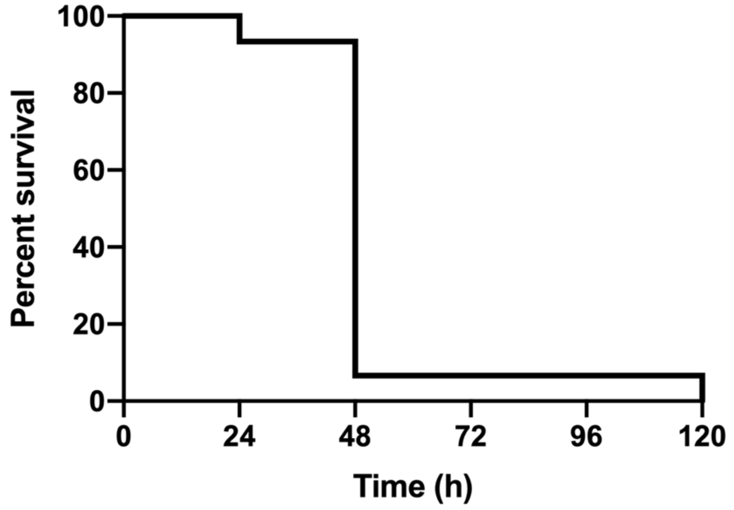

Sau54 (CC15)

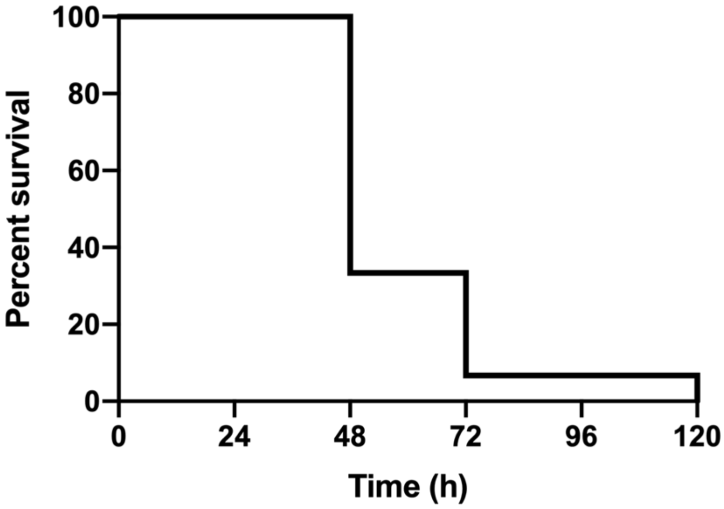


Sau55 (CC5)

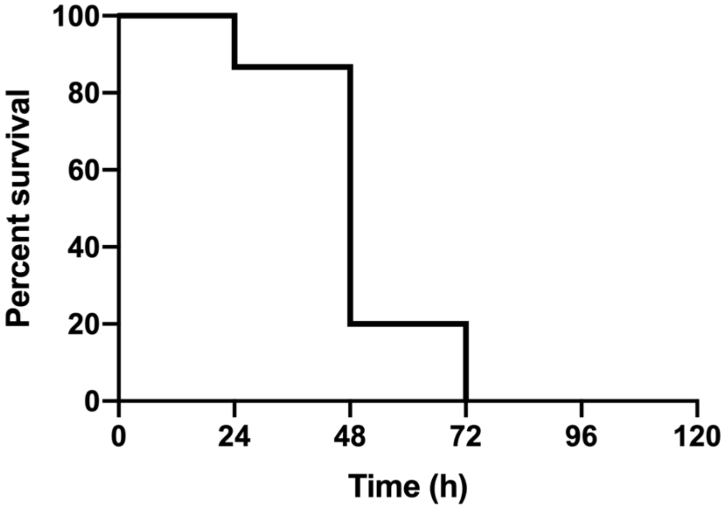


Sau56 (ST398)

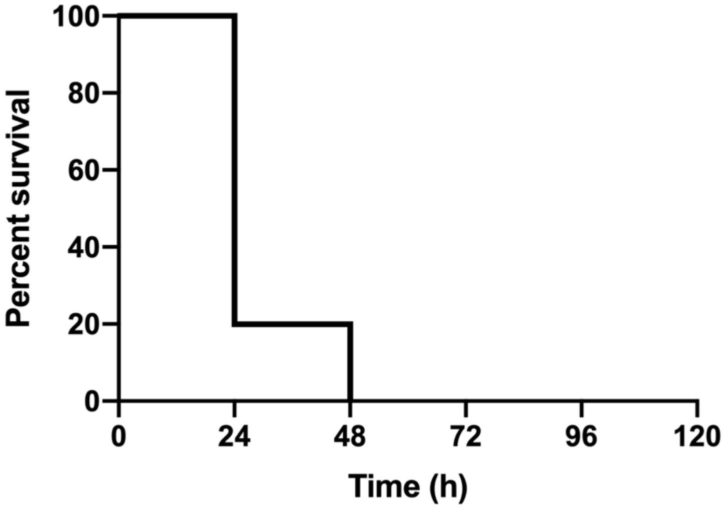

Sau58 (CC22)

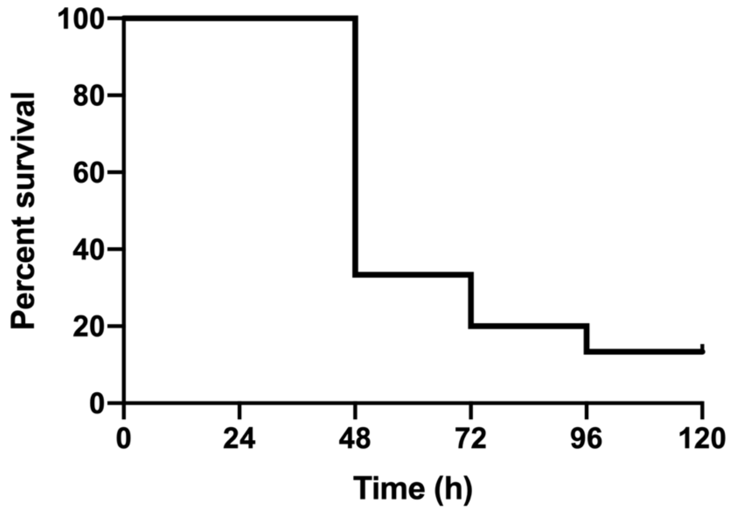


Sau59 (CC15)

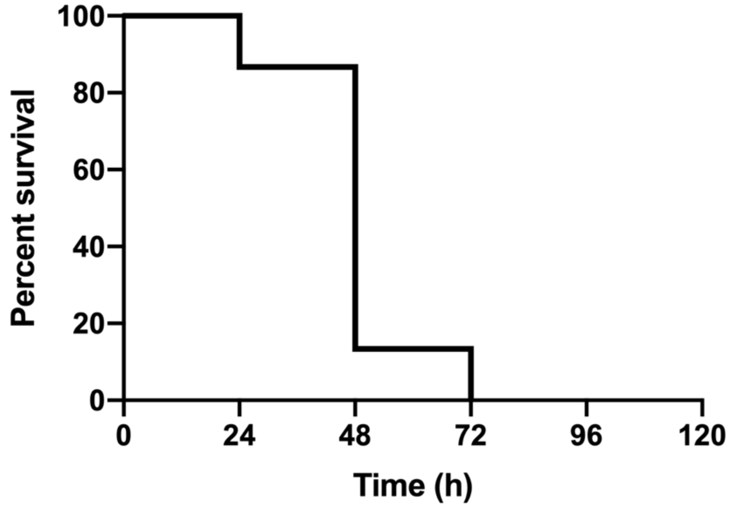


Sau60 (CC5)

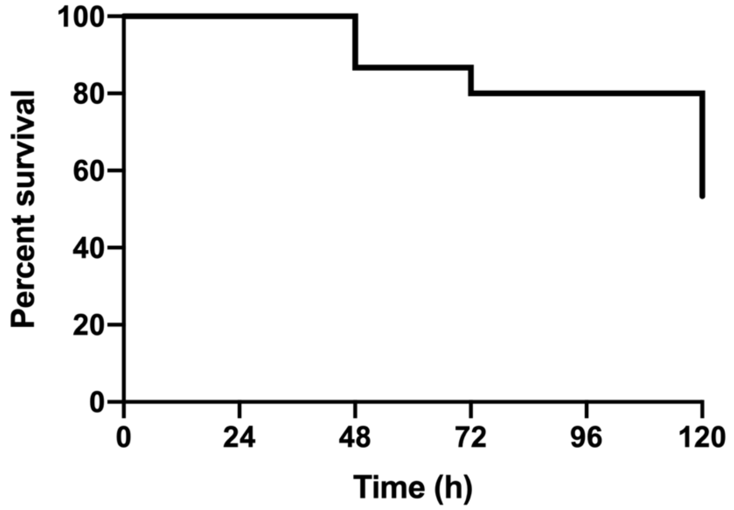

Sau62 (CC30)

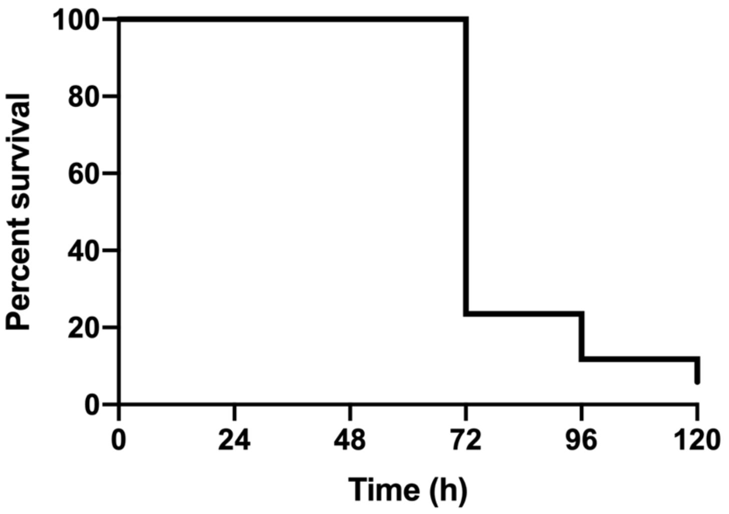


Sau65 (CC30)

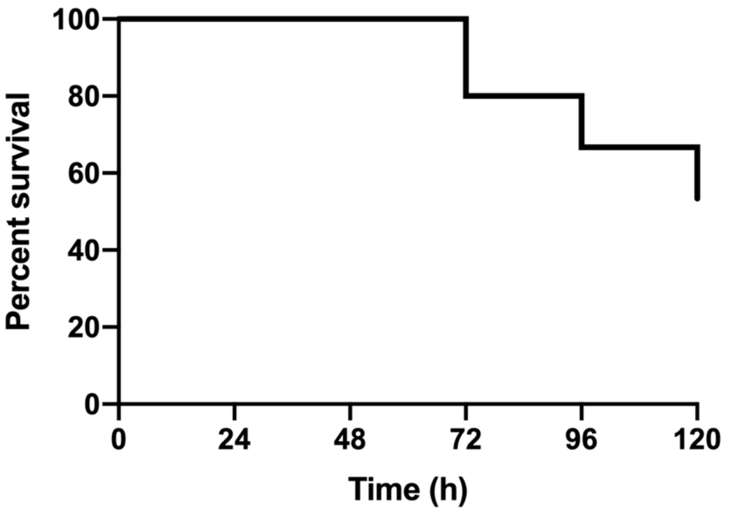


Sau66 (CC30)

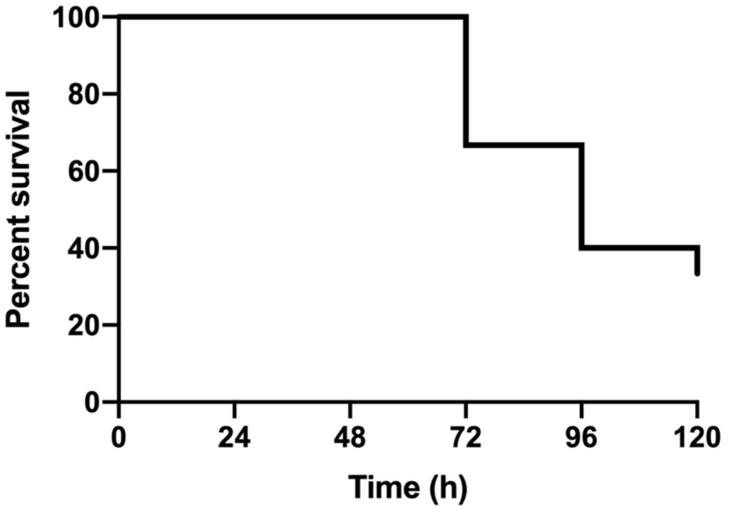

**Sau69 (CC5)**

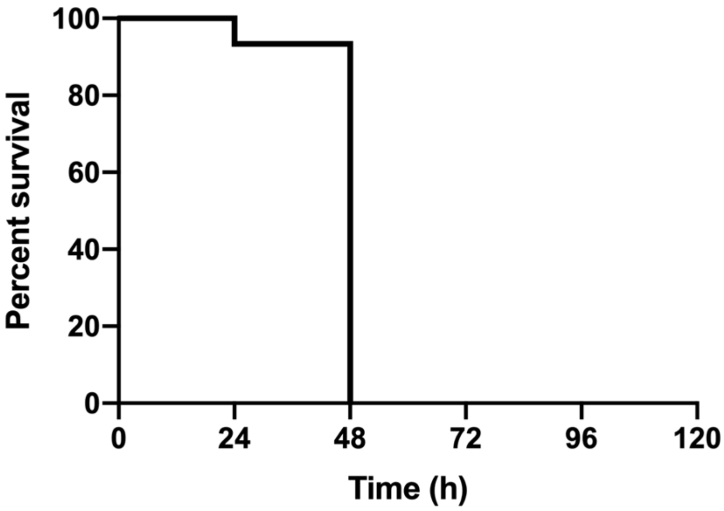


**Sau71 (CC5)**

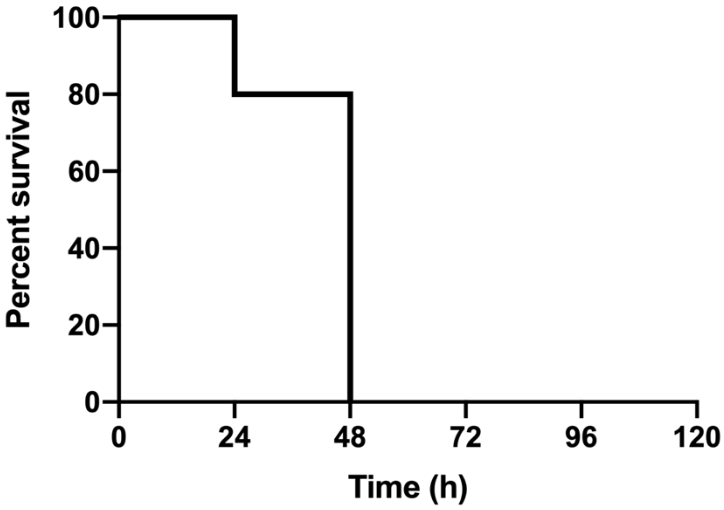


**Sau72 (CC8)**

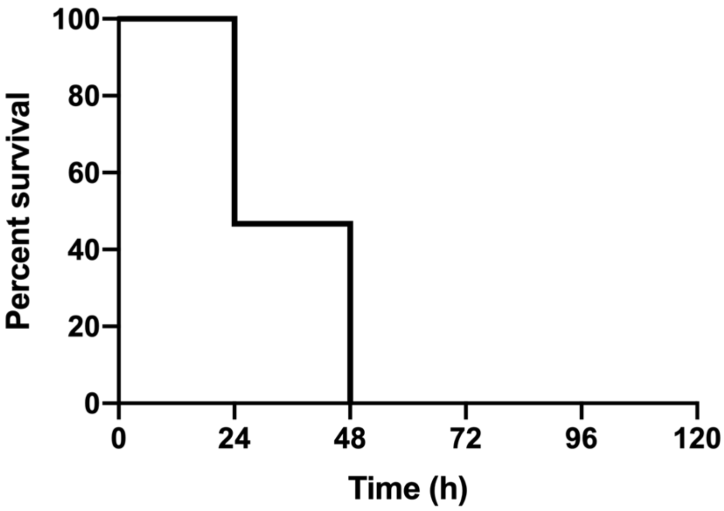

Sau74 (CC1)

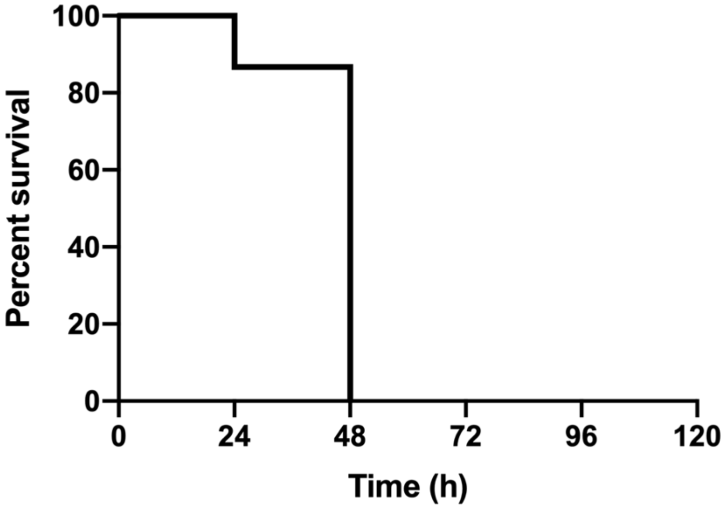


Sau75 (CC45)

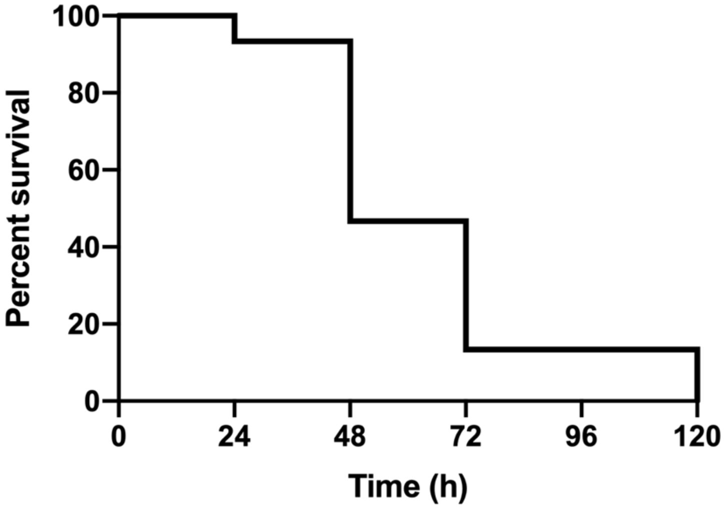


Sau79 (CC22)

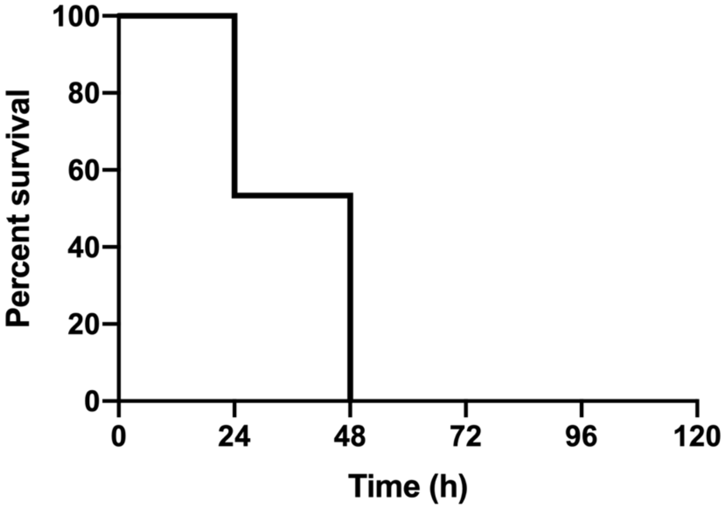

**Sau81 (CC5)**

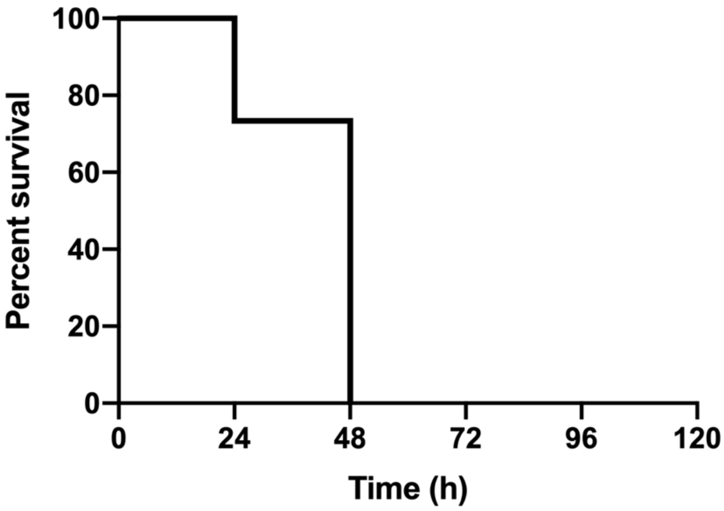


**Sau84 (ST398)**

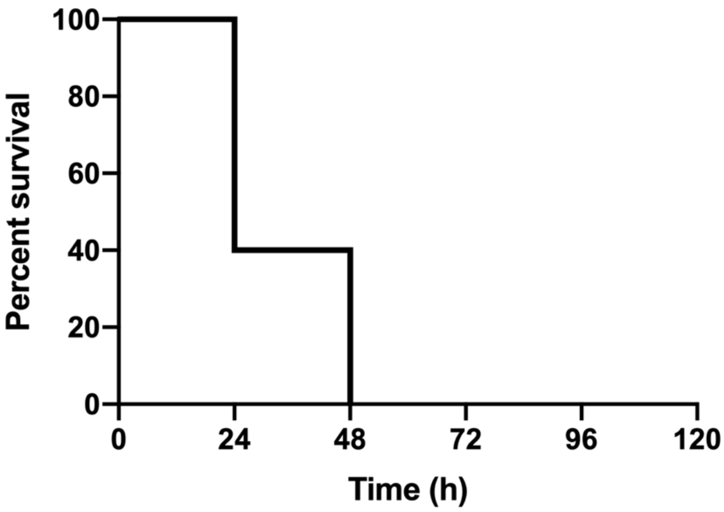


**Sau85 (CC30)**

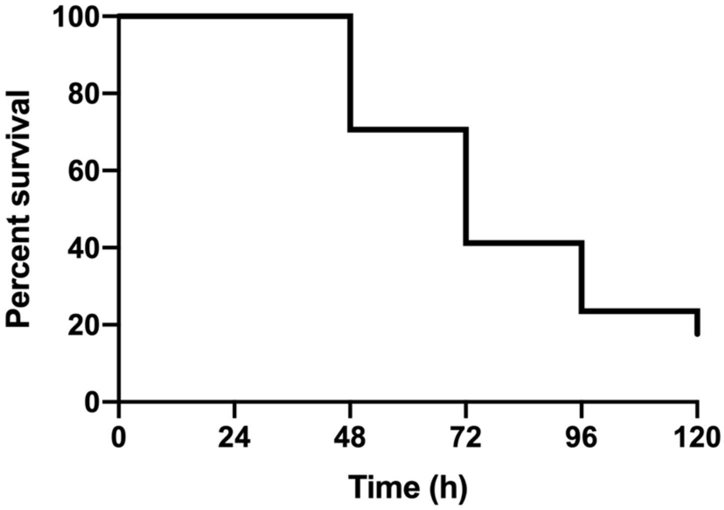

Sau88 (CC30)

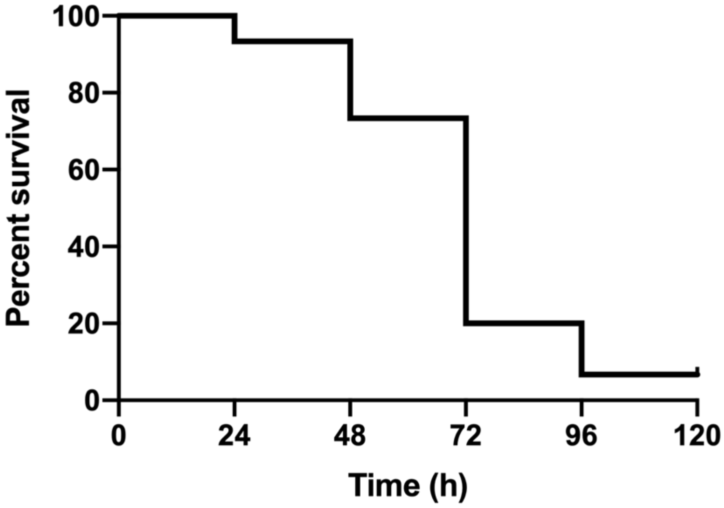


Sau89 (CC30)

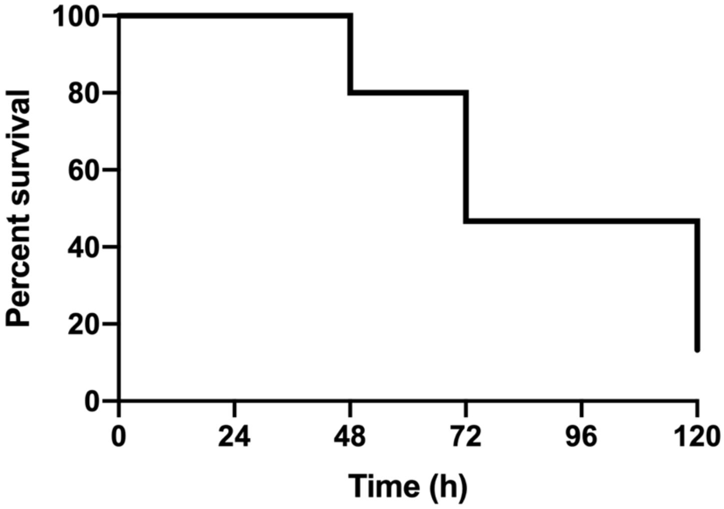


Sau91 (CC22)

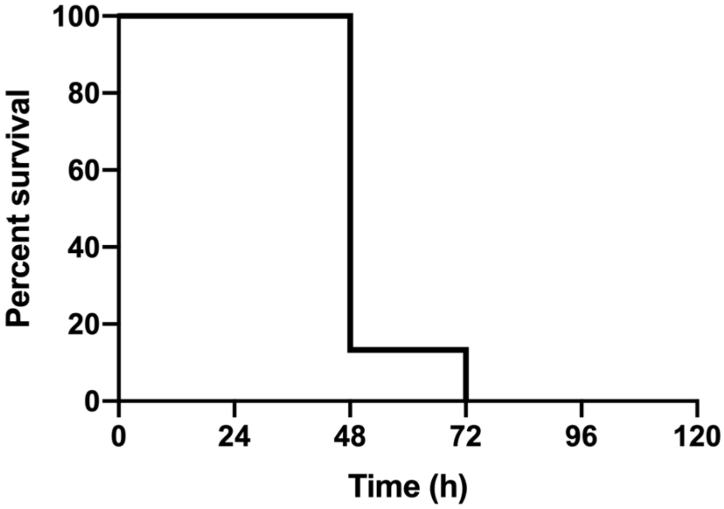

Sau92 (CC121)

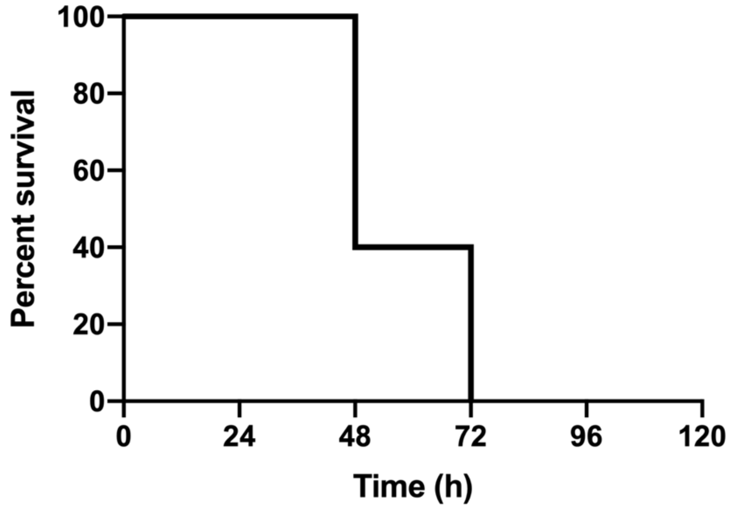


Sau94 (ST291)

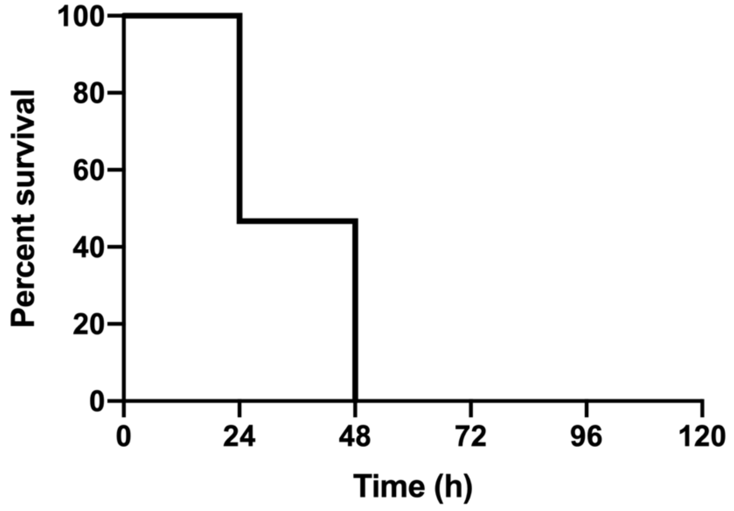


Sau96 (CC45)

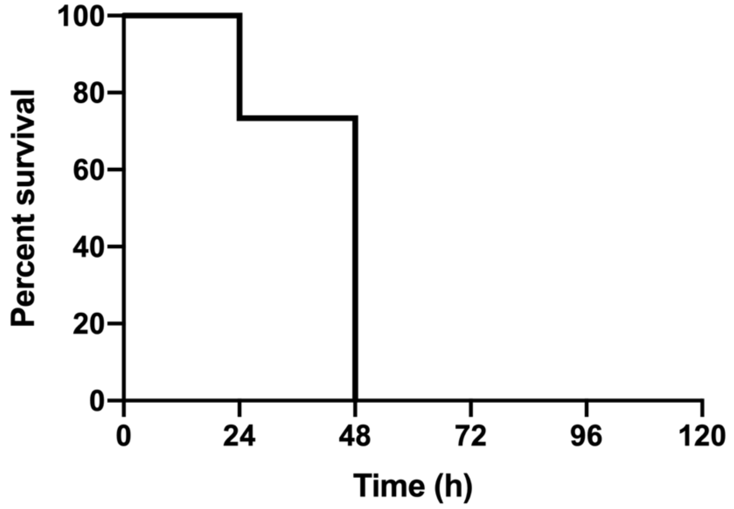

**Sau99 (CC1)**

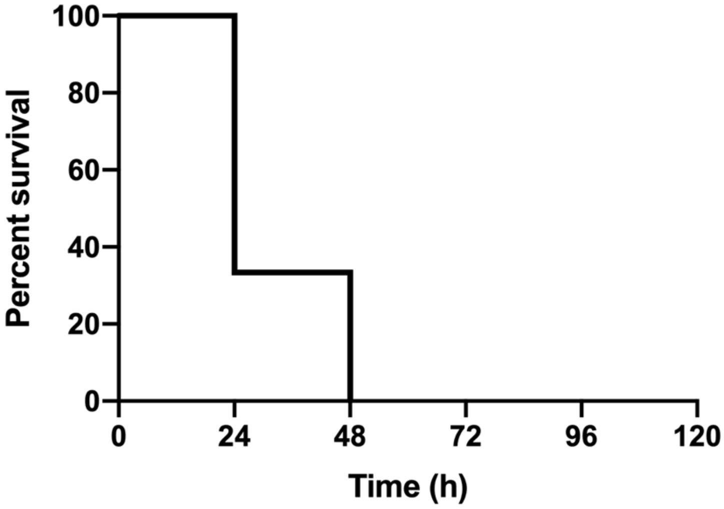


**Sau100 (CC5)**

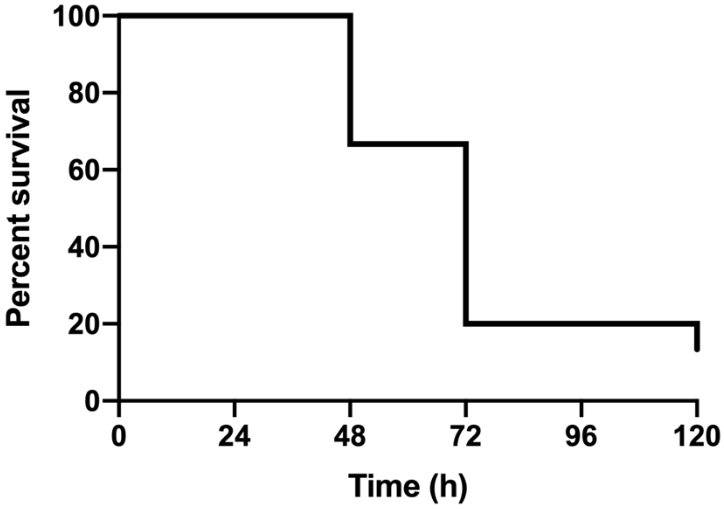


**Sau101 (ST398)**

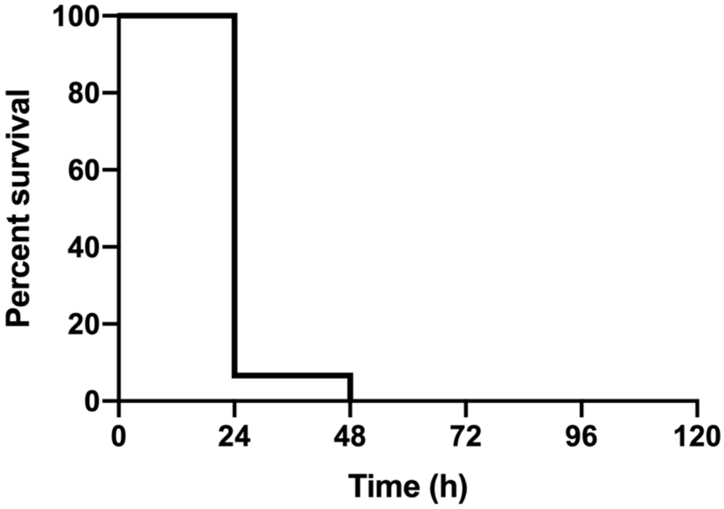

Sau103 (ST7)

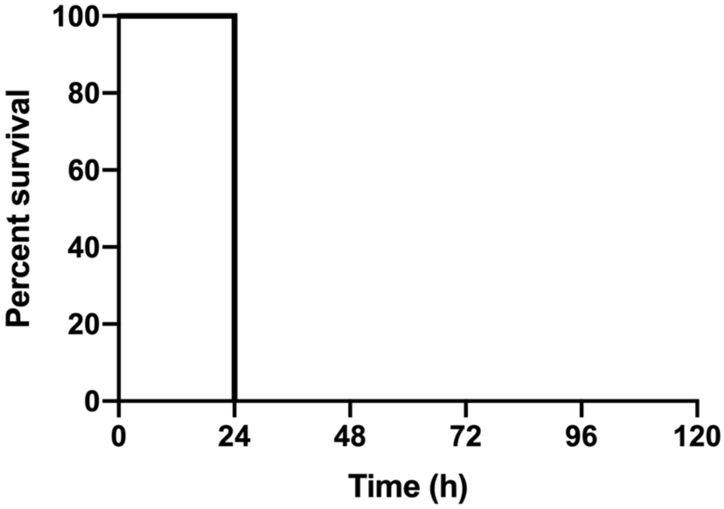


Sau107 (ST7)

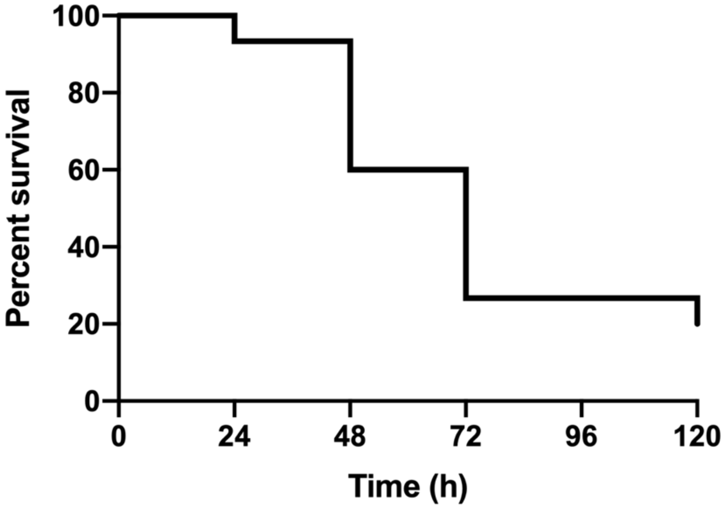


Sau108 (CC8)

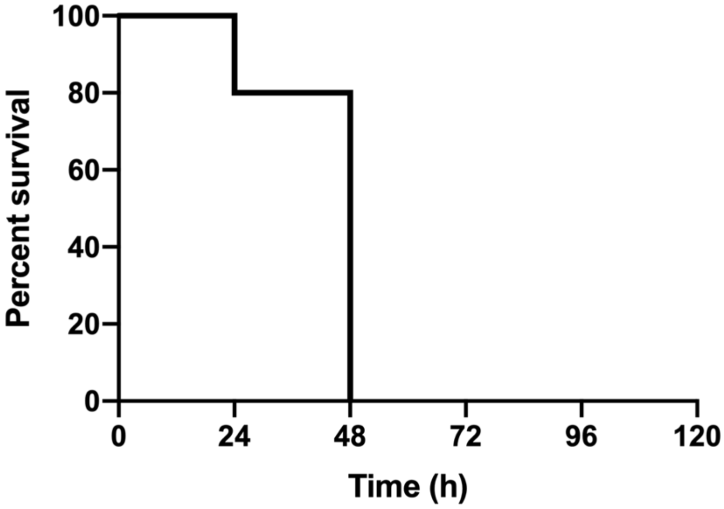

Sau112 (CC45)

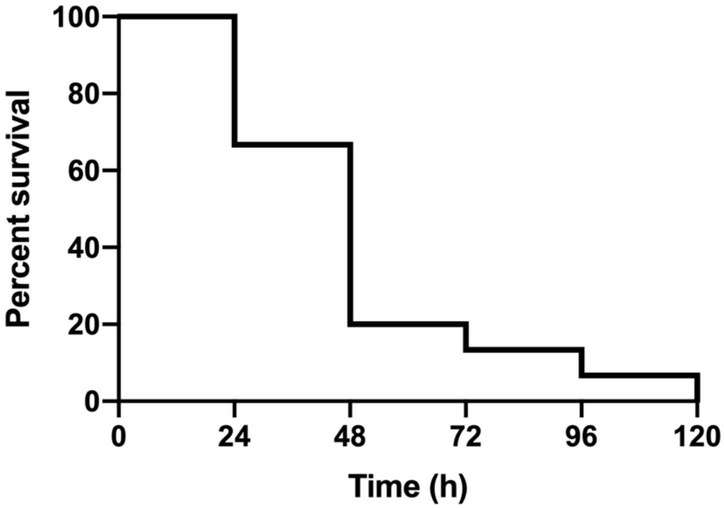


Sau115 (ST398)

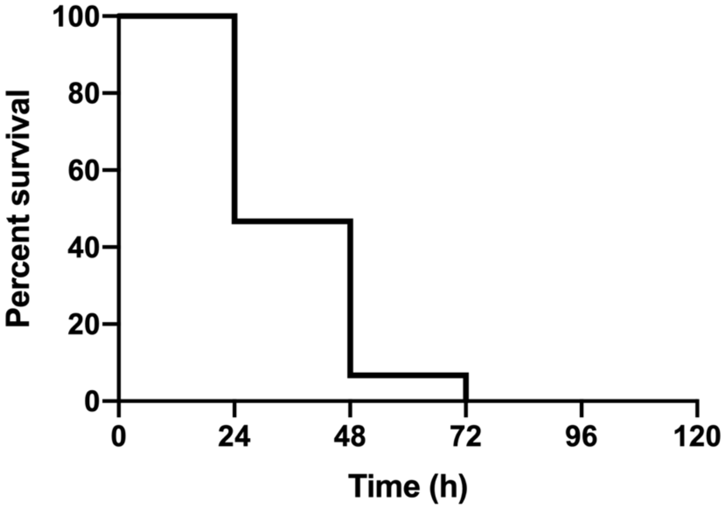


Sau116 (ST398)

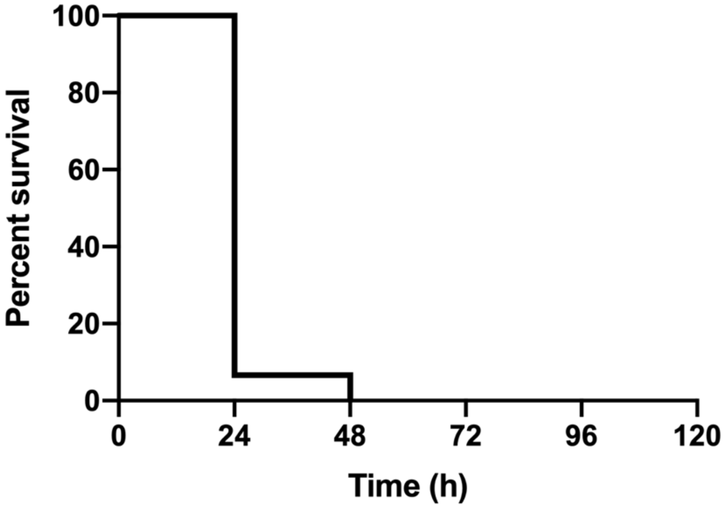

Sau119 (CC30)

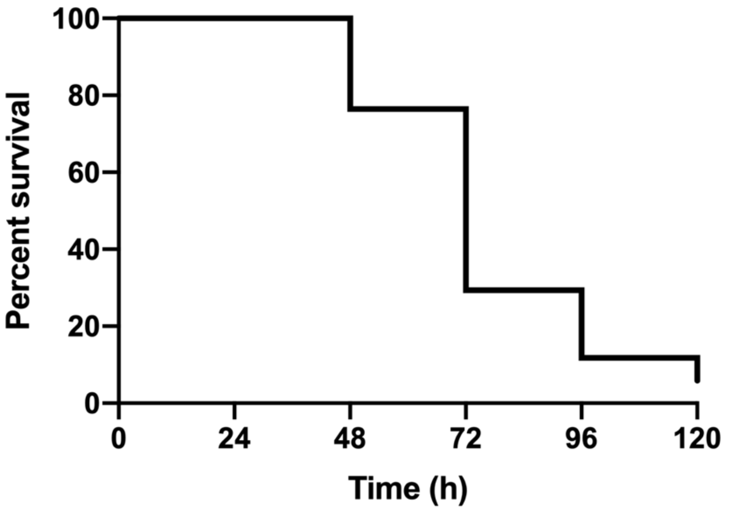


Sau123 (CC15)

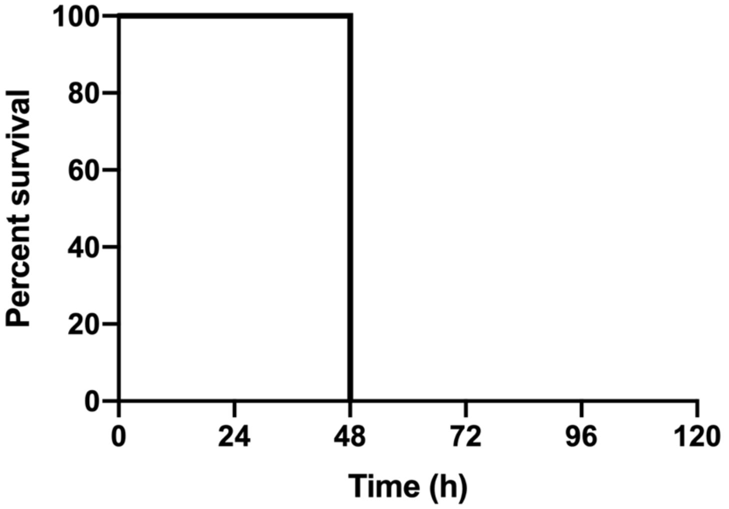


Sau124 (CC30)

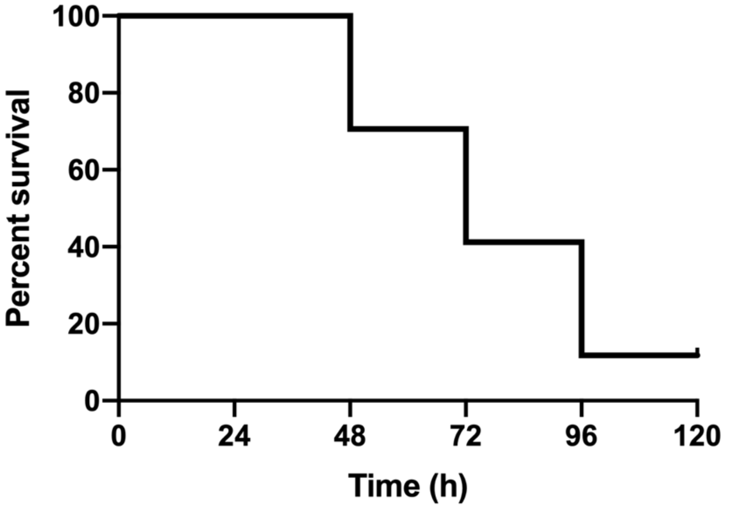

Sau132 (CC30)

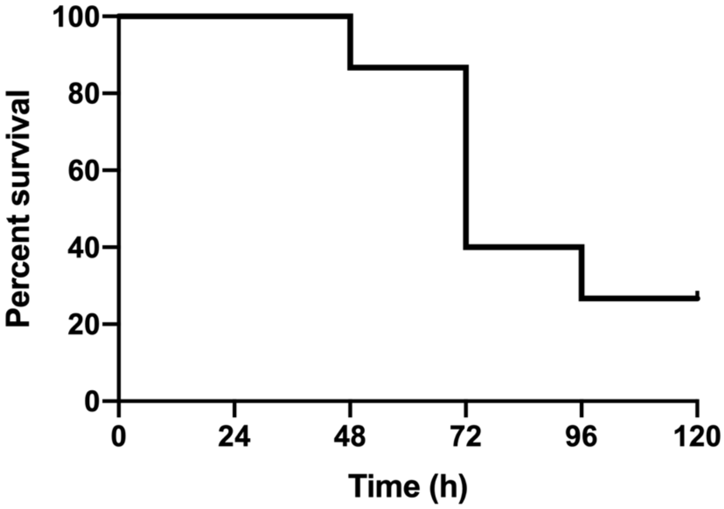


Sau133 (CC30)

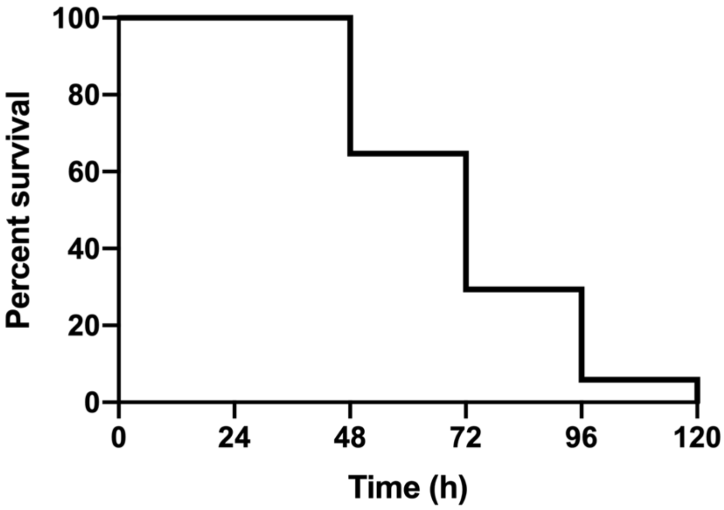


Sau135 (ST398)

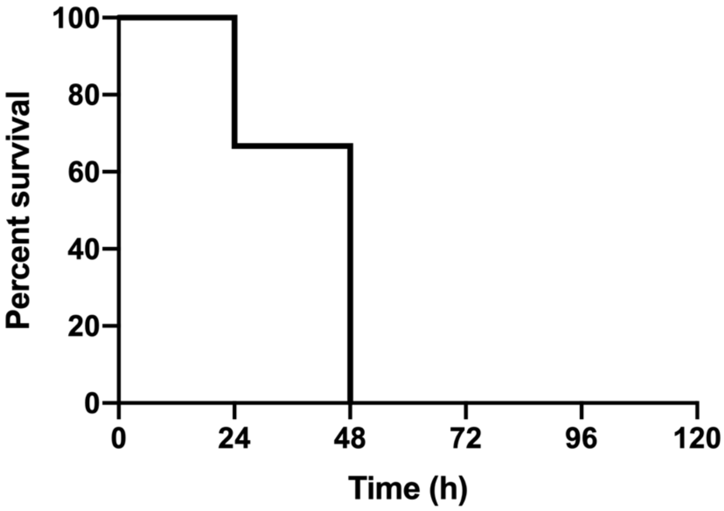

Sau137 (CC22)

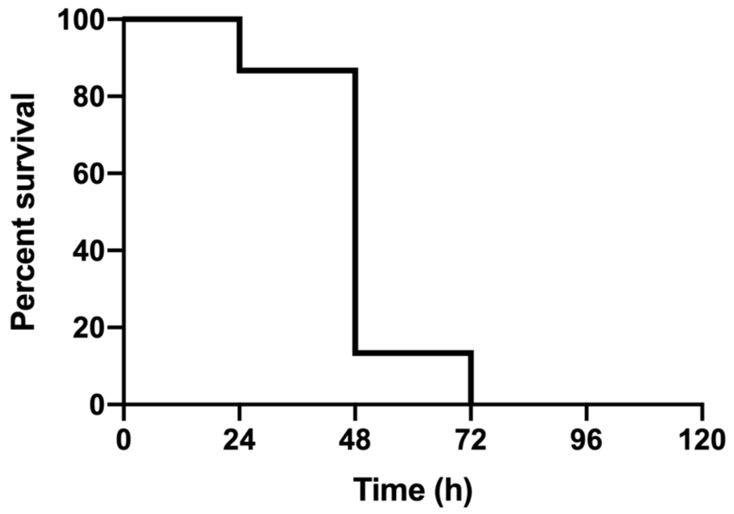


Sau139 (ST398)

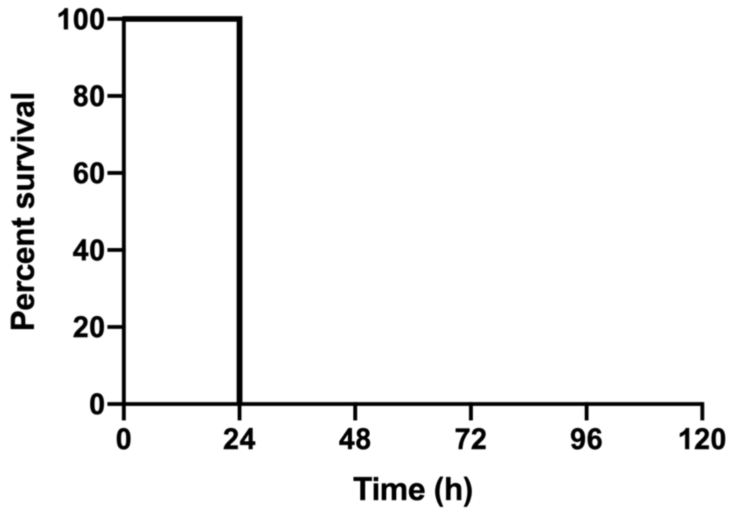


Sau146 (CC30)

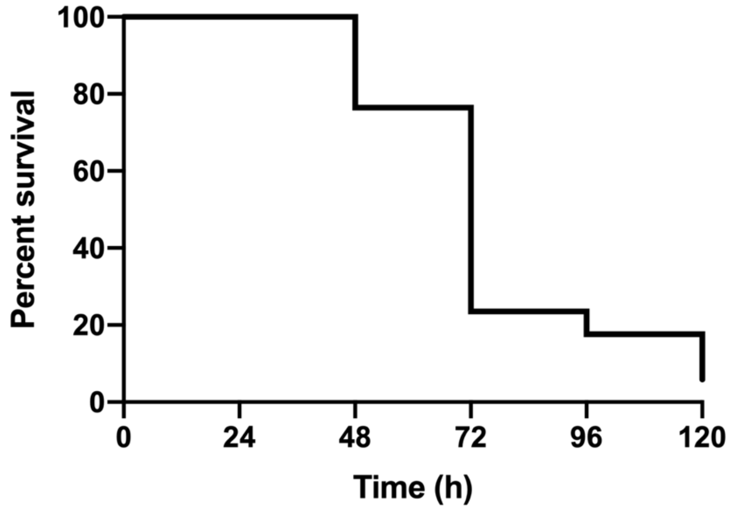

Sau147 (CC1)

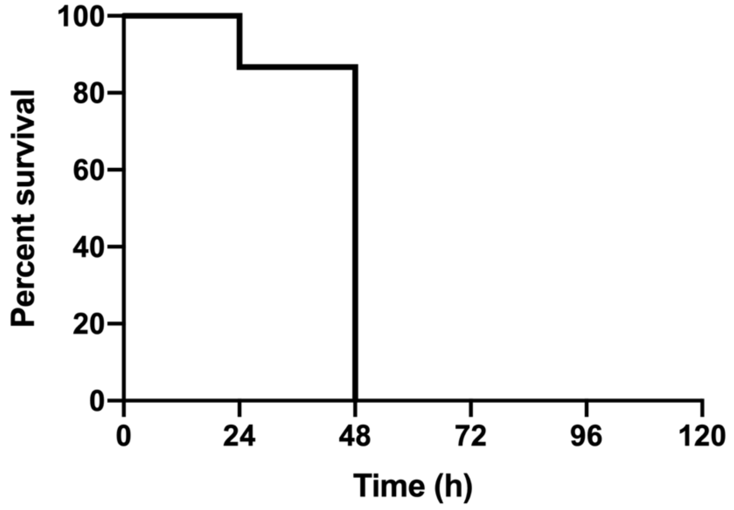


Sau148 (CC45)

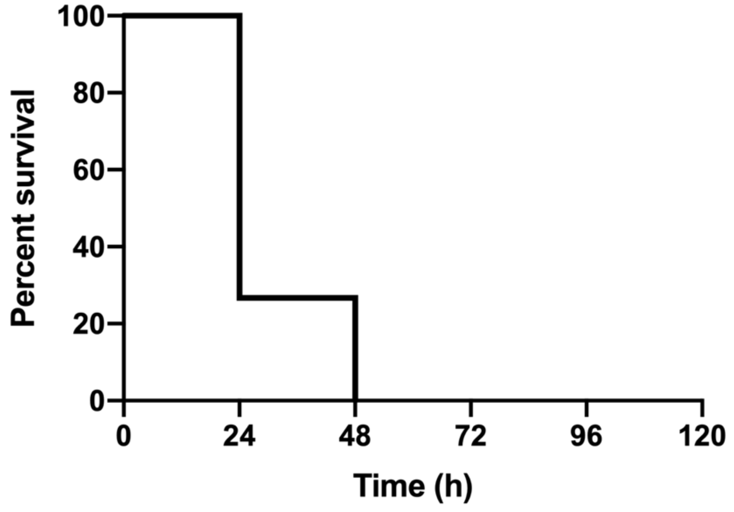


Sau149 (CC30)

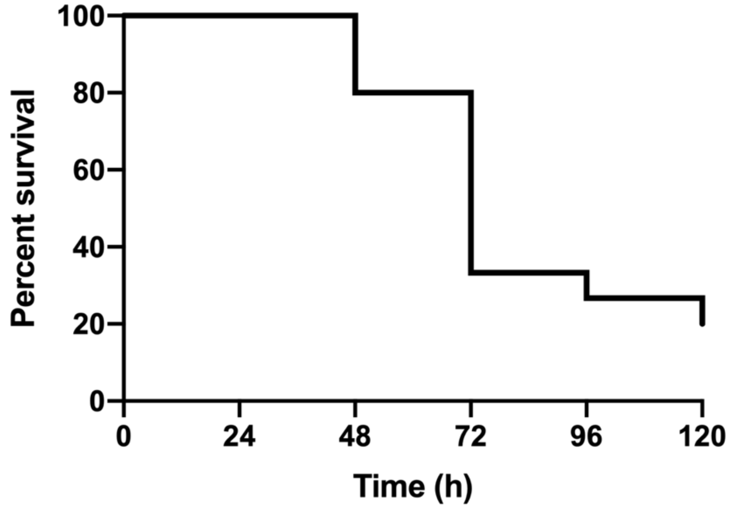

Sau150 (CC8)

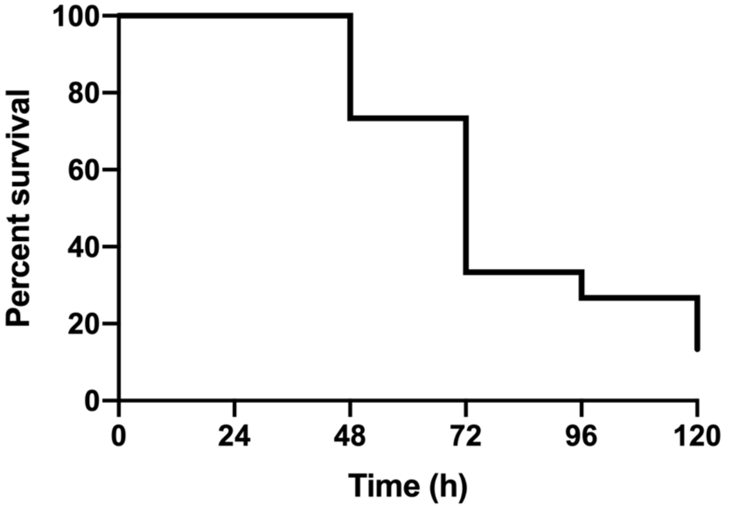


Sau152 (CC121)

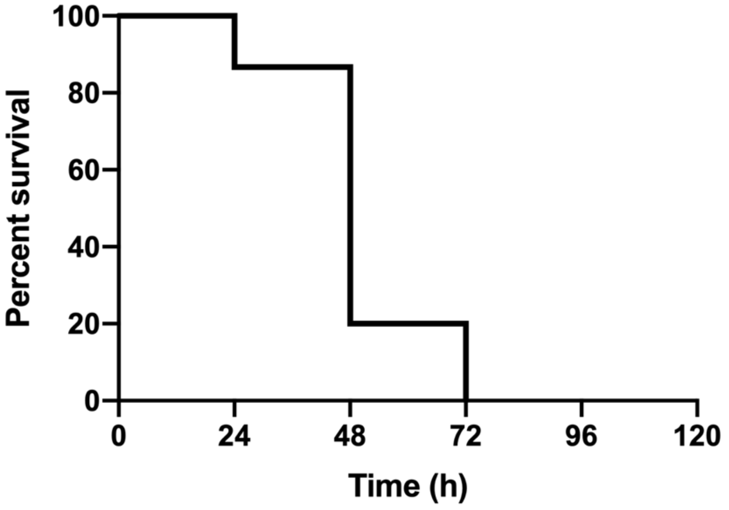


Sau153 (CC30)

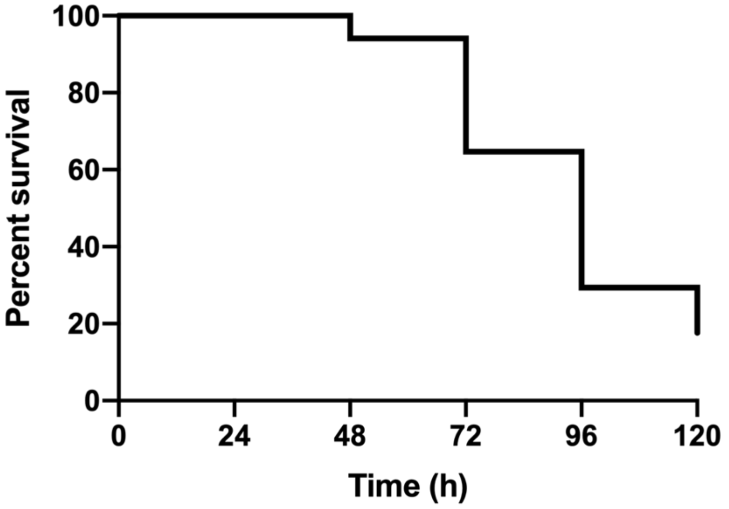

Sau158 (CC8)

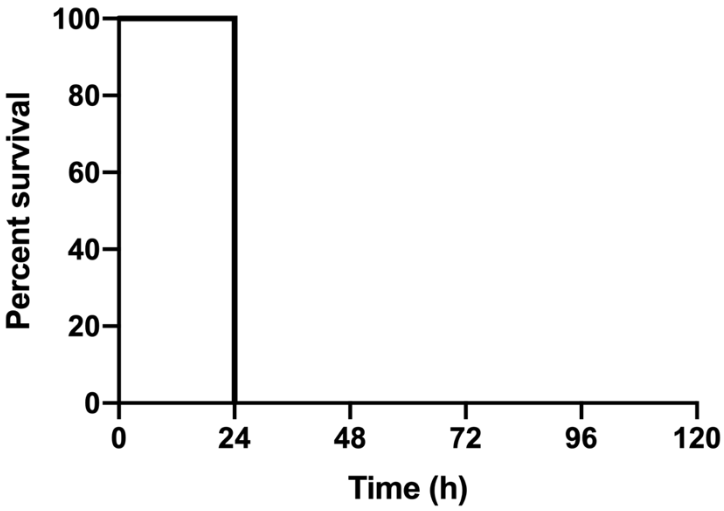


Sau162 (ST7)

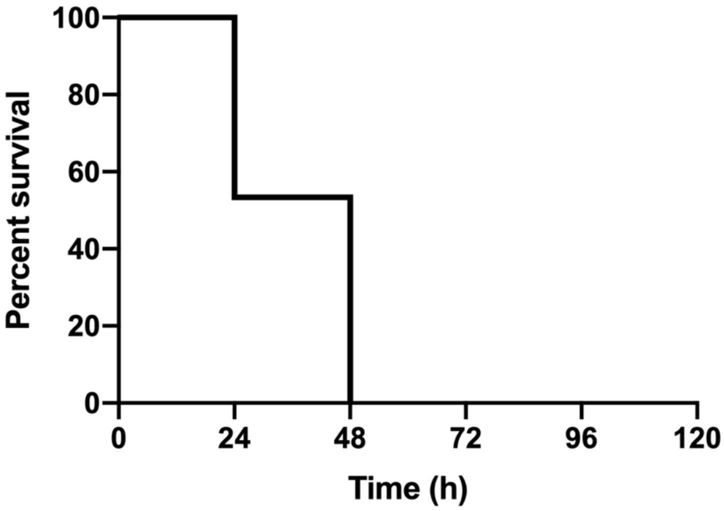


Sau163 (CC8)

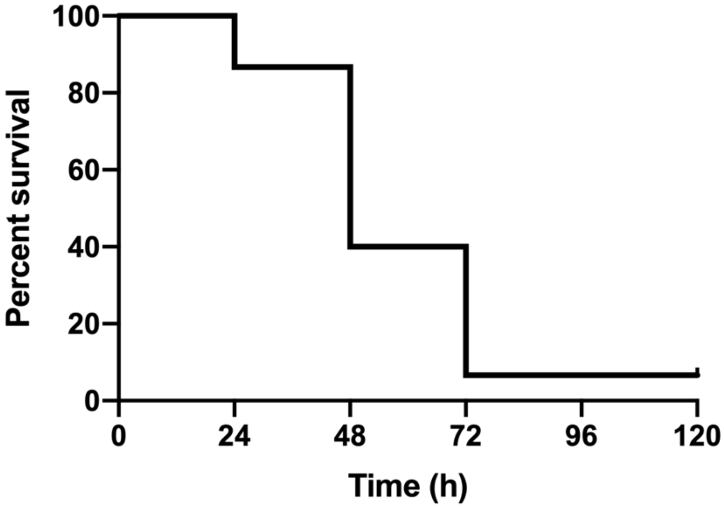

Sau164 (ST398)

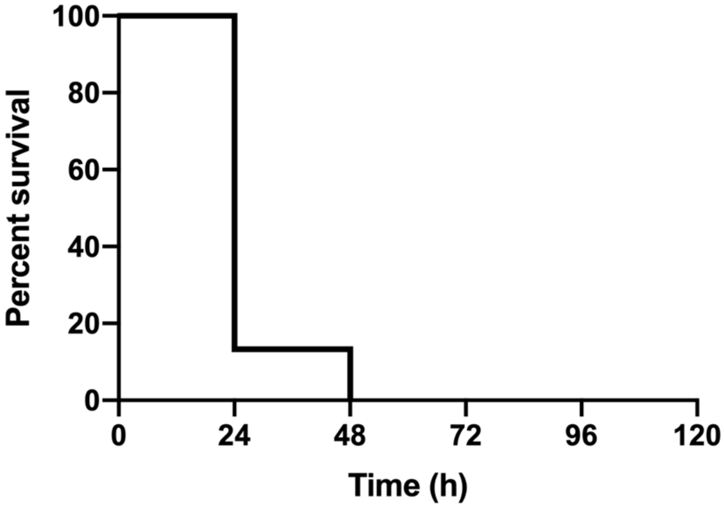


Sau165 (CC45)

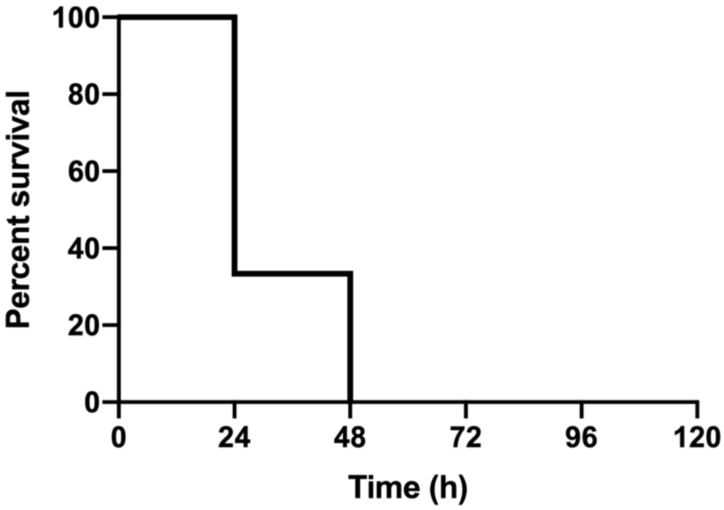


Sau166 (CC5)

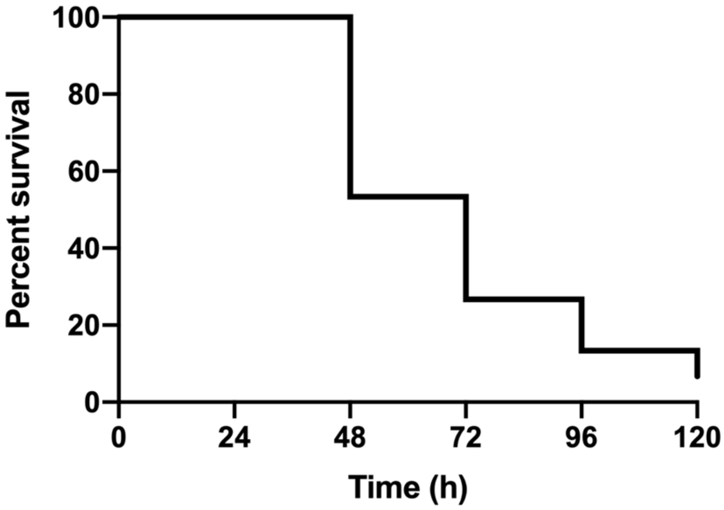

**Sau171 (CC1)**

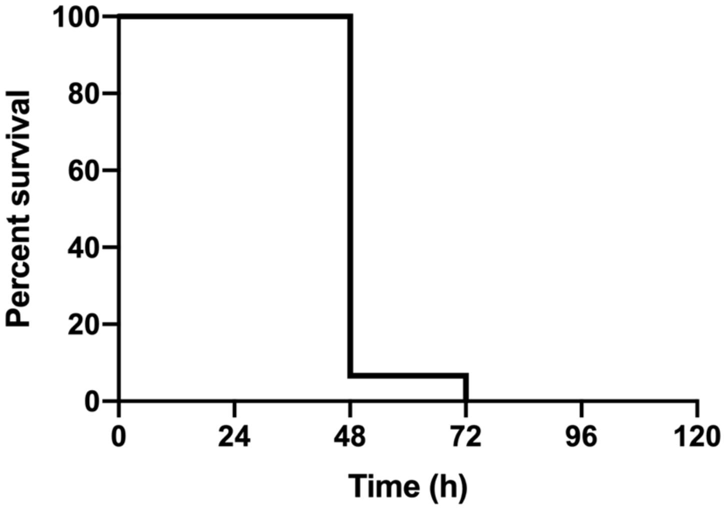


**Sau188 (CC1)**

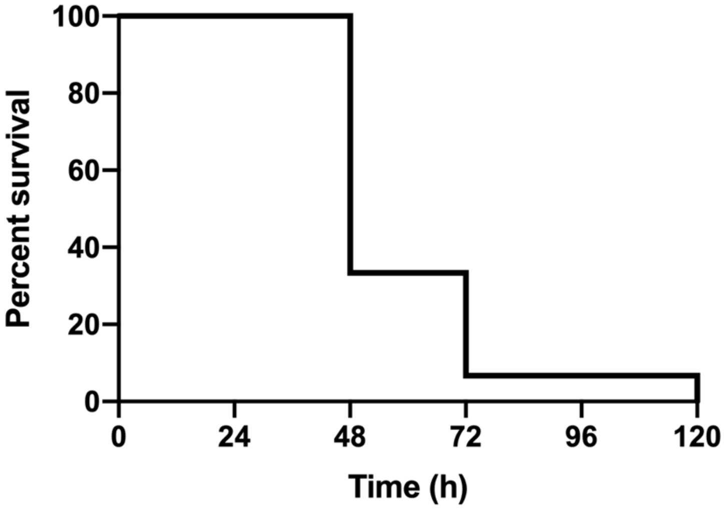


**Sau197 (CC1)**

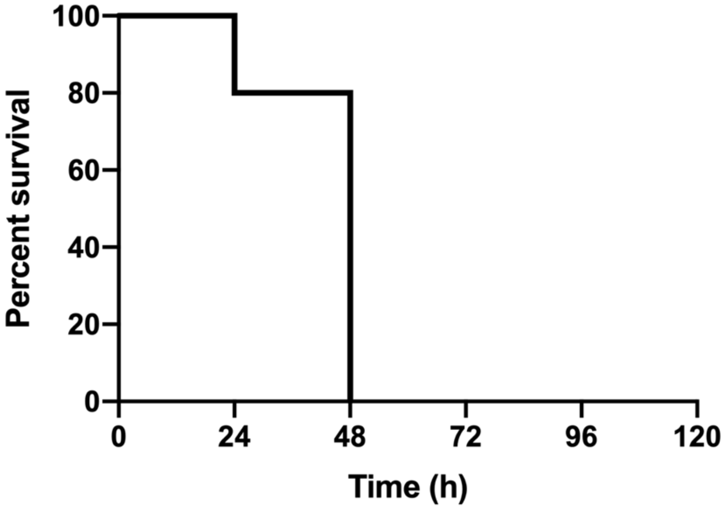

Sau210 (CC1)

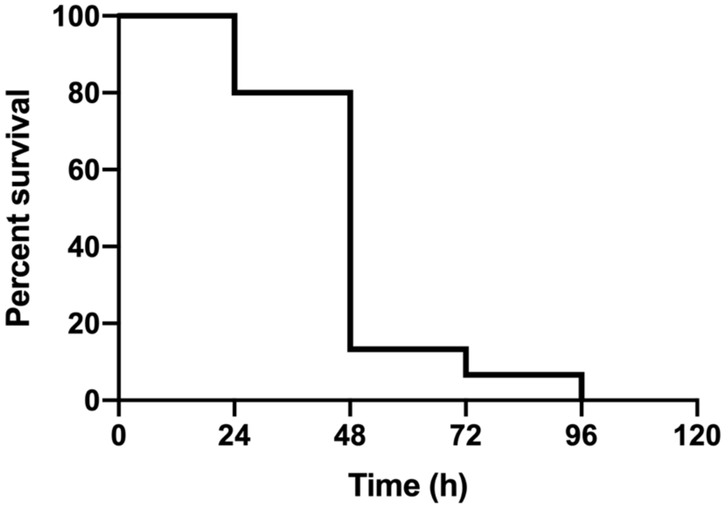


Sau229 (CC1)

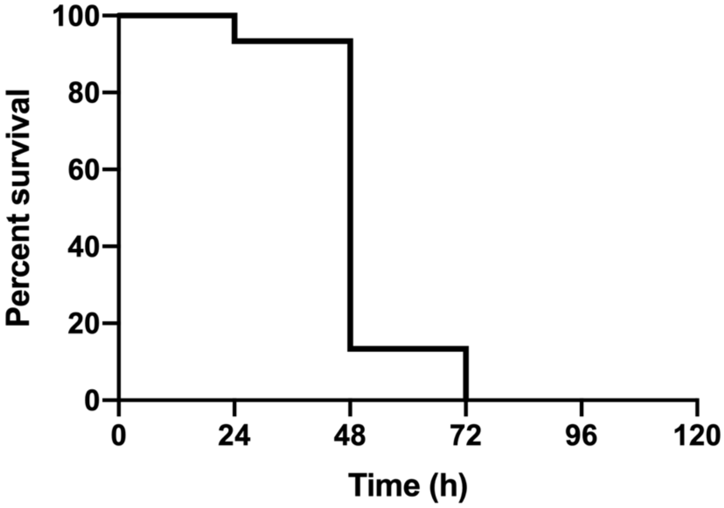


Sau275 (CC1)

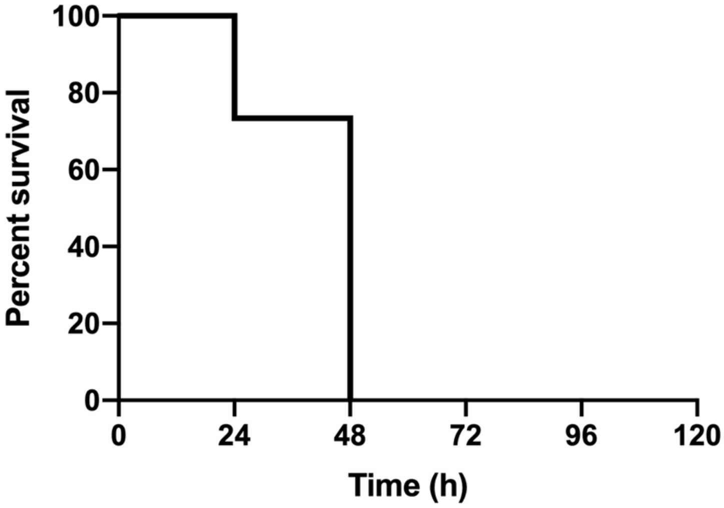

**Sau360 (CC1)**

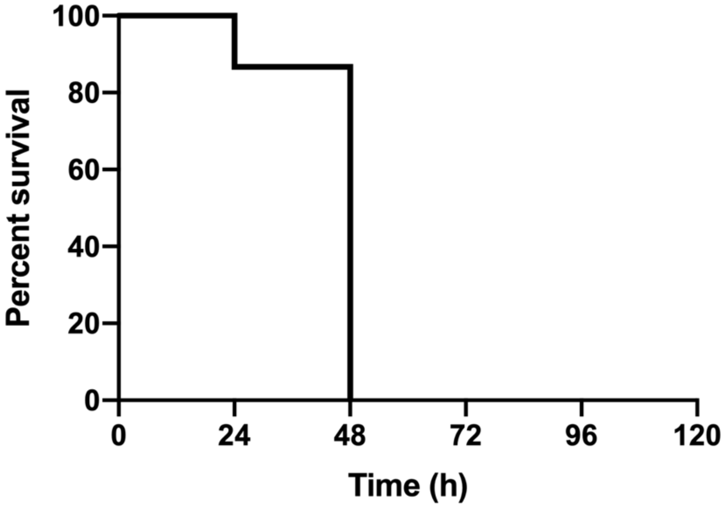


**Sauc6 (CC1)**

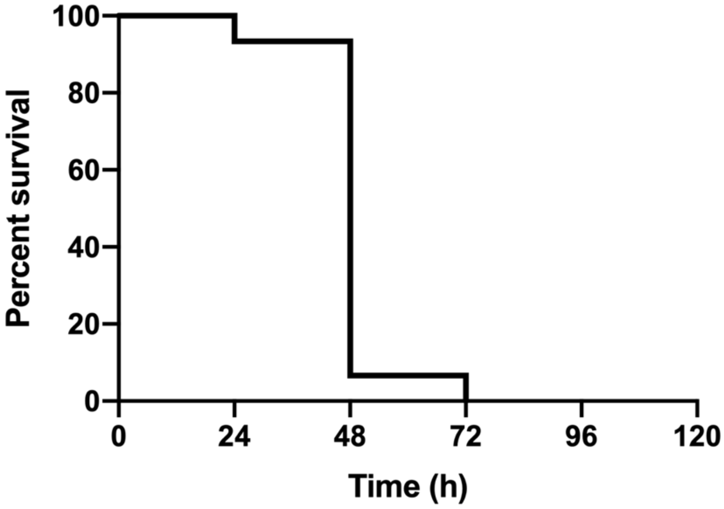
