## Supplementary figures and images for "Comparative mortality of dominant *Staphylococcus aureus* lineages in human bacteremia and animal infection models"

### Figure S3

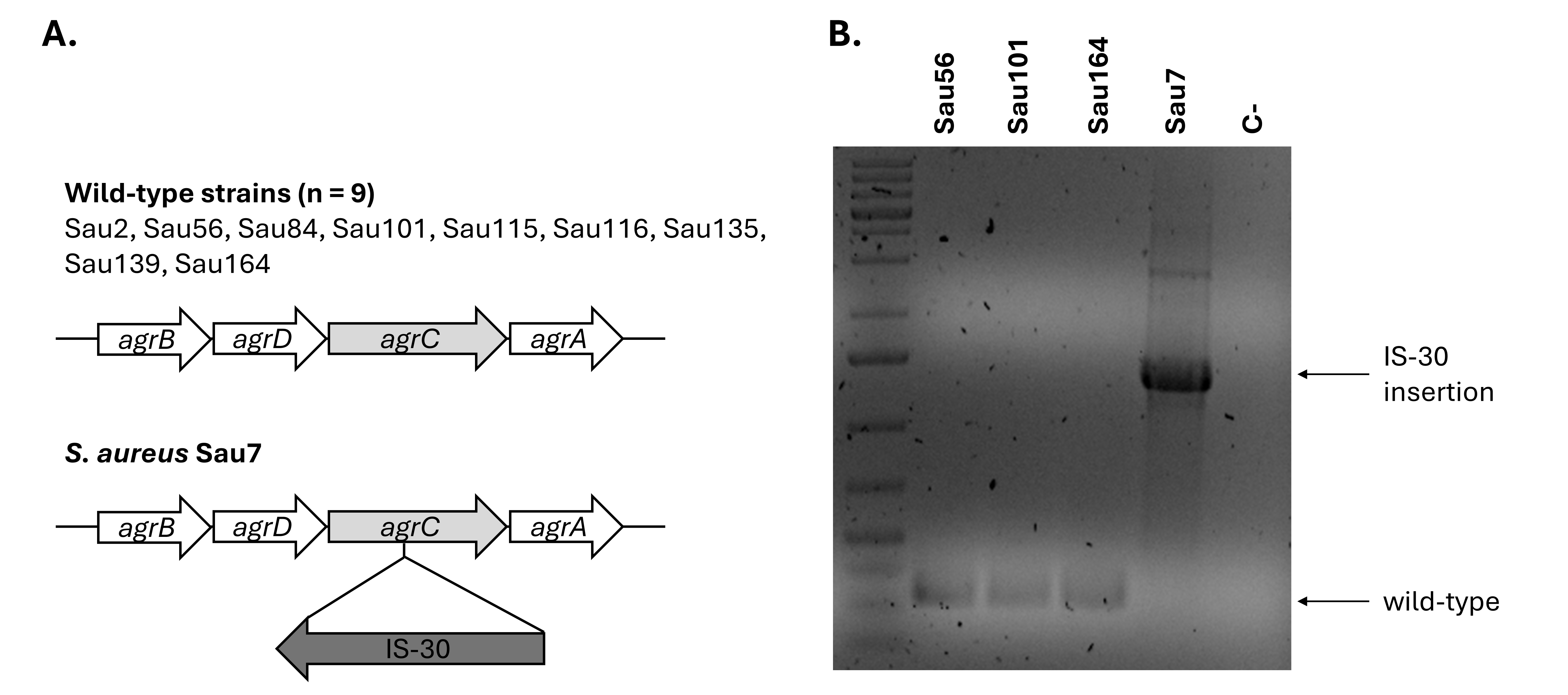

### Figure S4

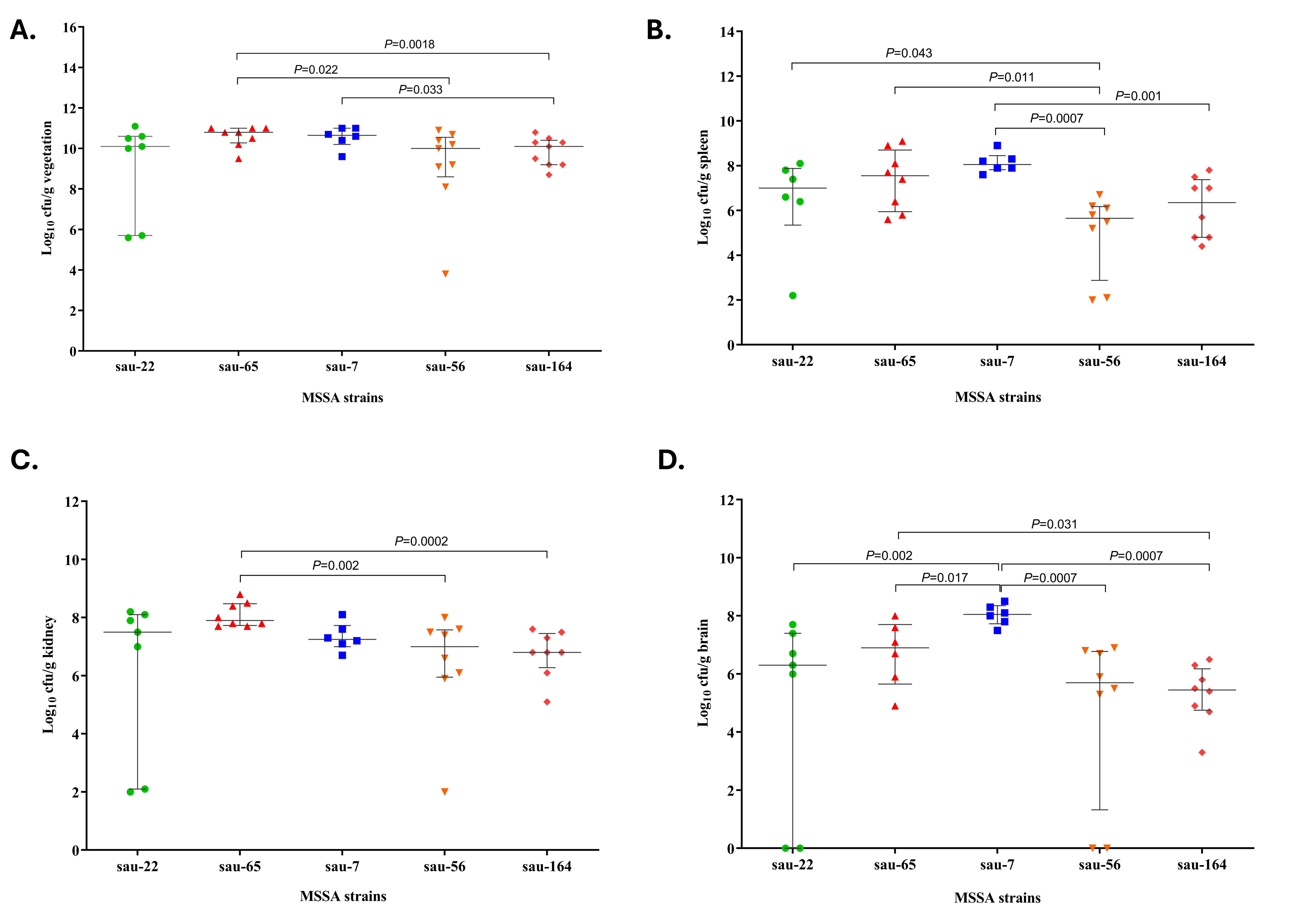
